# Congruent downy mildew-associated microbiomes reduce plant disease and function as transferable resistobiomes

**DOI:** 10.1101/2023.03.14.532520

**Authors:** P. Goossens, J. Spooren, K.C.M. Baremans, A. Andel, D. Lapin, N. Echobardo, C.M.J. Pieterse, G. Van den Ackerveken, R.L. Berendsen

**Author notes:** these authors contributed equally (P. Goossens, J. Spooren). Corresponding author: Roeland L. Berendsen.

## Abstract

Root-associated microbiota can protect plants against severe disease outbreaks. In the model-plant *Arabidopsis thaliana*, leaf infection with the obligate downy mildew pathogen *Hyaloperonospora arabidopsidis* (*Hpa*) results in a shift in the root exudation profile, therewith promoting the growth of a selective root microbiome that induces a systemic resistance against *Hpa* in the above-ground plant parts. Here we show that, additionally, a conserved subcommunity of the recruited soil microbiota becomes part of a pathogen-associated microbiome in the phyllosphere that is vertically transmitted with the spores of the pathogen to consecutively infected host plants. This subcommunity of *Hpa*-associated microbiota (HAM) limits pathogen infection and is therefore coined a “resistobiome”. The HAM resistobiome consists of a small number of bacterial species and was first found in our routinely maintained laboratory cultures of independent *Hpa* strains. When co-inoculated with *Hpa* spores, the HAM rapidly dominates the phyllosphere of infected plants, negatively impacting *Hpa* spore formation. Remarkably, isogenic bacterial isolates of the abundantly-present HAM species were also found in strictly separated *Hpa* cultures across Europe, and even in early published genomes of this obligate biotroph. Our results highlight that pathogen-infected plants can recruit protective microbiota via their roots to the shoots where they become part of a pathogen-associated resistobiome that helps the plant to fight pathogen infection. Understanding the mechanisms by which pathogen-associated resistobiomes are formed will enable the development of microbiome-assisted crop varieties that rely less on chemical crop protection.

## Main

Downy mildews are plant pathogenic oomycetes that cause major damage to a wide variety of plant species^1^. These pathogens have an obligate biotrophic lifestyle and are highly host-specific^2^. Plants have an intricate innate immune system, that relies on the recognition of the pathogen using cell-surface pattern recognition receptors, and R genes that encode intracellular nucleotide-binding leucine-rich repeat receptors (NLRs)^3^. In the model plant *Arabidopsis thaliana* (hereafter: Arabidopsis), resistance to its cognate downy mildew *Hyaloperonospora arabidopsidis* (*Hpa*) is based on the recognition of pathogen-produced immune-modulatory effector proteins by NLRs of the *Resistance to Peronospora parasitica* (*RPP*) family^4,5^.

However, disease severity does not solely depend on the efficacy of the immune system. The plant accommodates diverse microbial communities in virtually all its tissues. The highest population densities are generally observed in the rhizosphere, the part of the soil that is directly influenced by plant roots^6^. The aboveground plant parts, *i.e.* the phyllosphere, hosts microbial communities that are typically less abundant and diverse than those belowground^7^. Microbiomes in the rhizosphere and phyllosphere can protect plants from their attackers^8,9^. Beneficial microbes can inhibit pathogen growth through competition for nutrients and the production of antibiotics, or by increasing competence of the plant immune system, a phenomenon known as induced systemic resistance ^10,11^. Consequently, it has been argued that the microbiome functions as an extension of the plant immune system, providing an additional layer of defense against pathogen attack^12^.

Previously it was shown that leaf infection of Arabidopsis by *Hpa* results in the recruitment of a synergistically acting community of protective bacteria to its roots. The reconstituted three-member consortium consisting of a *Xanthomonas*, a *Stenotrophomonas*, and a *Microbacterium* sp. reduced susceptibility to *Hpa* and promoted plant growth^13^. Plants sown and grown on a soil previously occupied by *Hpa*-infected plants were more resistant to *Hpa* than plants grown on soil conditioned by uninfected plants^13^. Infections by other leaf pathogens such as the bacterium *Pseudomonas syringae*^14,15^, the fungus *Pseudopestalotiopsis camelliae-sinensis*^16^, or the root-feeding insect herbivore *Delia radicum*^17^ were later also shown to result in a beneficial microbial community in the soil that can persist and protect a subsequent population of plants. Thus, plants dynamically steer their microbiota upon pathogen attack to create a persistent protective soil microbiome, called a soil-borne legacy (SBL)^18,19^. This was similarly shown in agricultural fields, where the buildup of beneficial microbes resulted in disease-suppressive soils^20,21^.

On the other hand, pathogens can also modulate neighboring microbiota to their benefit. Like all microbes, pathogens have many mechanisms by which they can antagonize and outcompete their microbial competitors^22,23^. Disease-associated microbiomes are thus likely shaped by a combination of plant- and pathogen-driven selective pressures. Moreover, it is becoming increasingly clear that disease is often not caused by a single microbial species alone. Frequently, multiple pathogen symbionts are found to facilitate the effects of the primary pathogen^24^. It is, thus, argued that diseases are caused by the combined action of pathogens and their microbial entourage, called pathobiomes^25,26^.

As an obligate biotrophic pathogen, *Hpa* laboratory cultures are maintained by weekly passaging of spores from diseased to healthy plants^27^. Different isolates of *Hpa* with distinct effector repertoires are cultured on specific Arabidopsis accessions that lack the corresponding *RPP* gene(s)^5,27^. Therefore, laboratory *Hpa* cultures represent systems in which distinct pathogen isolates have, since their isolation in the early 1990’s^28^, completed many lifecycles on their host in association with a microbiome. We hypothesized that these *Hpa* cultures are rich in microbes that associate with the downy mildew pathogen and initially thought they would benefit the pathogen.

Here, we investigated phyllosphere microbiomes of Arabidopsis plants following the inoculation with distinct *Hpa* isolates using amplicon sequencing of *Hpa* cultures. We found remarkable similarities between the *Hpa*-associated microbiota (HAMs) of distinct *Hpa* strains and cultures. We show that phyllosphere HAM members benefit from *Hpa* infection, but that reversely HAM reduces *Hpa* sporulation when co-inoculated with *Hpa* spores on the leaves. Finally, we show that the plant-protective HAM can survive in soil as a SBL to subsequently colonize both roots and shoots of a subsequent plant population and limit *Hpa* infection. Together our data suggest that diseased plants can assemble a pathogen-associated protective microbiome, coined “resistobiome”, that can be transmitted to subsequent plant populations growing in the same soil.

## Results

### Distinct *Hpa* cultures are dominated by congruent disease-associated microbiomes

To study whether the obligate downy mildew pathogen *Hpa* is associated with a microbiome that potentially modifies the disease outcome, we investigated phyllosphere microbiomes of Arabidopsis plants following inoculation with two distinct isolates of *Hpa*, Noco2^29^ and Cala2^28^. These isolates have been routinely and separately cultured in our laboratory since 1999. Two-week-old plants of the Arabidopsis accessions C24^30^, Col-0, *Ler*, and Pro-0^31^ were inoculated with Noco2 or Cala2, a mix of both isolates, or with sterile water (mock). These accessions were selected as they are susceptible to both *Hpa* isolates, to either Noco2 or Cala2, or resistant to both isolates (Fig. 1A, Supplementary Fig. S1). Seven days post-inoculation (dpi), when *Hpa* had started sporulating on the susceptible accessions, we analyzed phyllosphere microbial community composition of the inoculated plants. The bacterial phyllosphere community composition was strongly affected by *Hpa* inoculation as determined by amplicon sequencing of the 16S rRNA genes (Fig. 1B). Inoculation with either Noco2 or Cala2 spore suspensions led to phyllosphere communities that were significantly different from each other *(P<0.05* in permutational multivariate analysis of variance (PERMANOVA); Supplementary Table S1). Mix-inoculated samples appeared as intermediate between Noco2 and Cala2 inoculated samples (Fig. 1B). Even though also plant genotype had a small but significant effect, the phyllosphere bacterial community changed following inoculation with *Hpa* regardless of plant accession’s susceptibility (Fig. 1B, Supplementary Table S2-S3). This shows that the *Hpa* spores were co-inoculated with a bacterial community and suggests that the co-inoculated bacteria subsequently strongly affected the composition of the phyllosphere microbiome. In contrast, when fungal community compositions of a subset of samples were analyzed by amplicon sequencing of the ribosomal internal transcribed spacer (ITS2) region, no statistically significant differences between mock- and *Hpa*-inoculated plants were observed (Supplementary Fig. S2).

**Figure 1.**
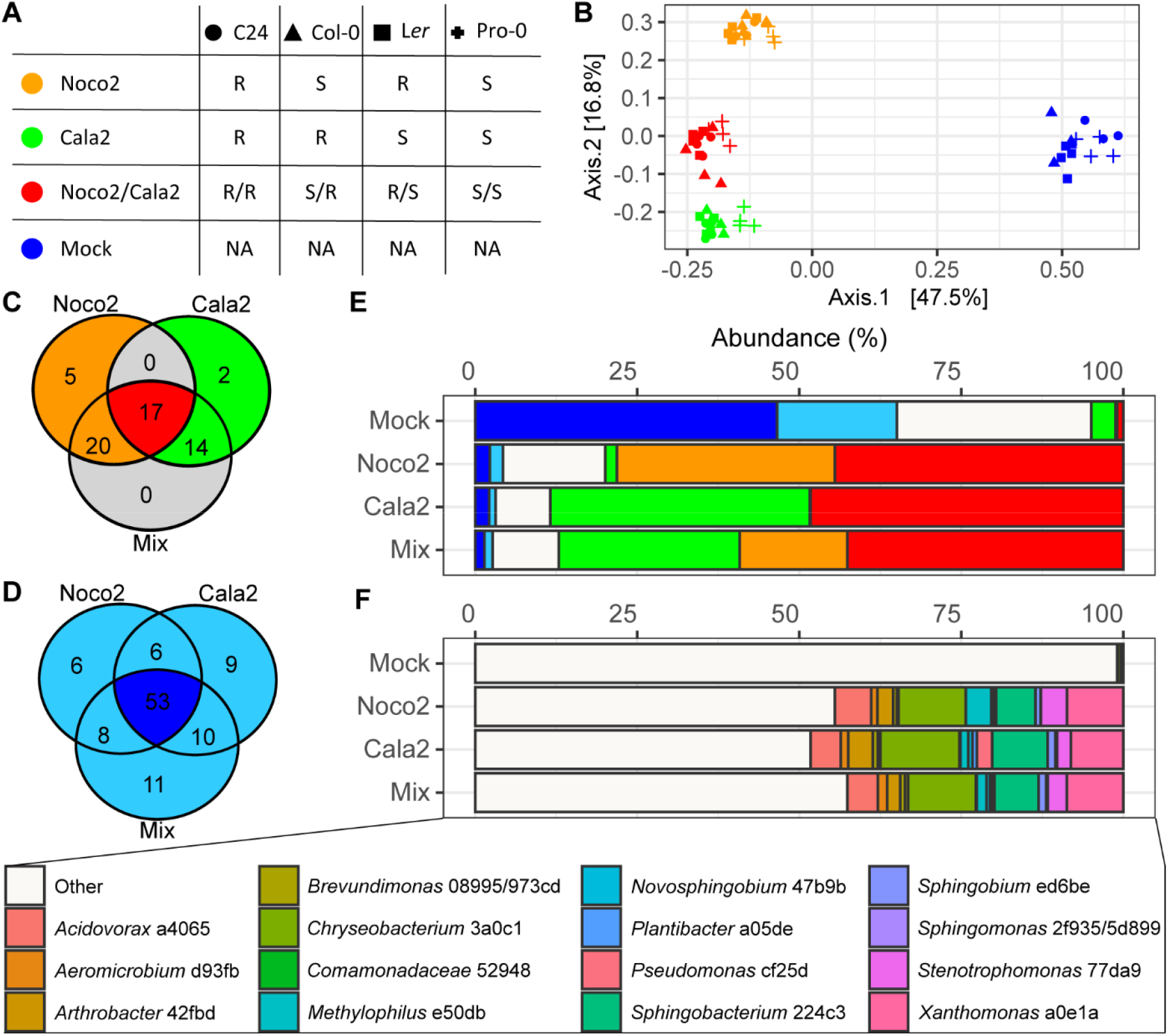
Distinct *Hpa* cultures are enriched for identical ASVs that dominate the phyllosphere bacterial communities. (**A**) Overview of the susceptibility of each accession to Noco2 and Cala2. S: Susceptible. R: Resistant. *Hpa* isolate color codes and *Arabidopsis thaliana* accession shapes also function as a guide to colors and shapes in (**B**); PCoA ordination plot based on Bray-Curtis dissimilarities of the bacterial phyllosphere communities of *Arabidopsis thaliana* accessions Col-0, C24, L*er*, or Pro-0 following inoculation with sterile water (mock) or spore suspensions of *Hpa* isolate Noco2, Cala2 or a mix of both isolates. Venn diagrams of (**C**) significantly enriched and (**D**) significantly depleted ASVs in *Hpa*-inoculated plants compared to mock-inoculated plants. (**E**) Stacked chart with the relative abundance of the 17 ASVs that are enriched in all *Hpa*-inoculated groups (red); those that were enriched in either Noco2-inoculated (orange) or Cala2-inoculated (green) plants; the 53 ASVs that were depleted in all *Hpa*-inoculated groups (dark blue) or ASVs that were depleted in two or less *Hpa*-inoculated groups (lighter blue); and all other ASVs in the data (white) in each inoculation group. (**F**) Stacked chart highlighting the abundances and taxonomies of the ASVs that were significantly enriched in Noco2- and Cala2-inoculated plants. ASVs are colored by taxonomy as indicated in the legend.

We identified 161 bacterial 16S amplicon sequence variants (ASVs) that significantly changed in abundance (*P* < 0.05, Deseq2^32^) as a result of inoculation with at least 1 *Hpa* isolate. Among them, 17 ASVs were enriched on both Noco2- as well as Cala2-inoculated plants (Fig. 1C). These 17 ASVs together occupy 45% of the bacterial phyllosphere communities on the *Hpa*-inoculated plants (Fig. 1C,E,F, Supplementary Fig. S3). Conversely, 53 ASVs were significantly depleted on all *Hpa*-inoculated plants, but constituted 47% of the bacterial communities on mock-treated plants (Fig. 1D-E, Supplementary Fig. S3). Thus, although inoculation with each of the two *Hpa* isolates leads to a distinct phyllosphere microbiome, these microbiomes are dominated by bacteria with identical ASVs (Fig. 1F).

The Noco2 and Cala2 cultures in Utrecht have been maintained separately since their arrival in 1999. Cross contamination has always been monitored using plant accessions that are resistant to the *Hpa* isolate being cultured, but susceptible to other *Hpa* isolates. It is therefore remarkable to see that the same 17 ASVs form such a large part of the bacterial microbiome of both the Noco2 and the Cala2 culture in Utrecht. To test whether the enrichment for these distinct *Hpa-associated* microbiota is a local peculiarity or a genuine phenomenon for Arabidopsis downy mildews, we collected Col-0 and L*er* samples infected with Noco2, Cala2, or inoculated with water (mock) at a different location, the Max Planck Institute for Plant Breeding Research (Cologne, Germany), who independently also maintained the same isolates for many years. Consistent with observations made in Utrecht, *Hpa* inoculation with either Noco2 or Cala2 significantly altered the bacterial phyllosphere community (Fig. 2A). We found 57 ASVs enriched in at least two *Hpa* cultures from either or both geographic locations (hereafter: frequently *Hpa*-enriched ASVs; pink to red in Fig. 2B,C,D) including 4 ASVs enriched in all four *Hpa* cultures across both locations (hereafter: *Hpa*-core ASVs; Fig 2E; red in Fig. 2). These 57 frequently *Hpa*-enriched ASVs occupy up to 75% of the phyllosphere bacterial communities of *Hpa*-inoculated plants at both locations and are low abundant or undetected in mock-treated samples (Fig. 2C). The 57 frequently *Hpa*-enriched ASVs corresponded to taxonomically-diverse taxa representing 9 bacterial classes and 38 genera (Fig. 2D). The *Hpa-core* ASV a0e1a, annotated as a *Xanthomonas* sp., was consistently highly abundant in the four *Hpa* cultures that were investigated and comprised between 5 and 10% of the bacterial phyllosphere population in all inoculated samples (Fig. 2E). The other 3 *Hpa*-core ASVs were consistently enriched but less abundant than *Xanthomonas* a0e1a (Fig. 2E). As the Utrecht and Cologne *Hpa* cultures have been maintained separately for many years on different soil substrates, these results show that repeated passaging of *Hpa* and associated microbes during *Hpa* maintenance resulted in the enrichment of a similar small set of taxonomically-diverse bacterial ASVs.

**Figure 2.**
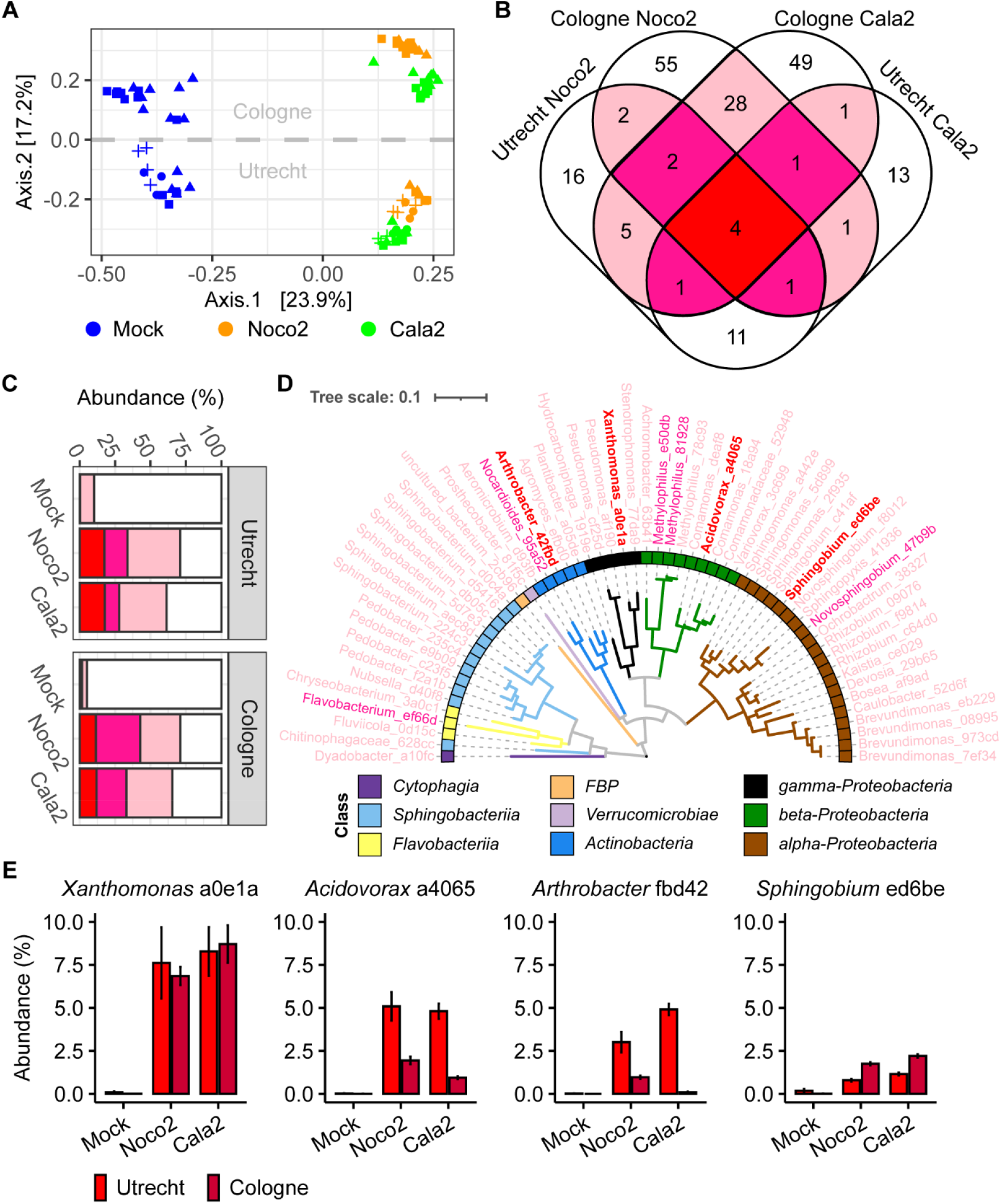
*Hpa* cultures from Utrecht and Cologne are enriched for identical ASVs. (**A**) PCoA ordination plot based on Bray-Curtis dissimilarities of the bacterial phyllosphere communities of *Arabidopsis thaliana* accession Col-0 (triangle symbols), C24 (circle symbols), L*er* (square symbols), or Pro-0 (plus symbols) following inoculation with sterile water (blue symbols) or a spore suspension of *Hpa* isolates Noco2 (orange symbols) or Cala2 (green symbols) that are cultured and maintained at Utrecht (lower half) and Cologne (upper half). (**B**) Venn diagram of frequently *Hpa*-enriched ASVs that are significantly enriched in 2 (pink), 3 (deep pink), or 4 (red) of the *Hpa* cultures. (**C**) Stacked chart with the cumulative relative abundance of the frequently *Hpa*-enriched ASVs that are enriched in 2 (pink), 3 (deep pink), or 4 (red) *Hpa* cultures. (**D**) Dendrogram based on MAFFT alignment of frequently *Hpa*-enriched ASVs that were significantly enriched in at least two out of four *Hpa* cultures. Colors of taxonomy labels indicate significant enrichment in 2 (pink), 3 (deep pink), or 4 (red) *Hpa* cultures. Colored squares indicate class-level taxonomy of ASVs in accordance with the legend. (**E**) Bar charts of mean abundances of the 4 ASVs that were consistently enriched in the Noco2 and Cala2 cultures at Utrecht (red) and Cologne (dark red). Error bars represent standard error.

### Distinct *Hpa* cultures contain isogenic HAM bacteria

To further characterize the HAM, we generated a collection of 702 bacterial isolates from Arabidopsis leaves that were infected by Noco2 or Cala2 in either Utrecht or Cologne and characterized the individual isolates by sequencing 16S rRNA amplicons. We subsequently sequenced the whole genome of 31 isolates that represented 8 HAM ASVs, including the 4 *Hpa-core* ASVs. The 31 genomes matched *Xanthomonas* HAM ASV a0e1a*, Acidovorax*, HAM ASV a4065, *Sphingobium* HAM ASV ed6be*, Arthrobacter* HAM ASV 42fbd*, Aeromicrobium* HAM ASV d93fb*, Methylobacterium* HAM ASV 15da8*, Microbacterium* HAM ASV f0c76, and *Rhizobium* HAM ASV 2569b. Genomes of isolates that matched with the same ASV were mostly isogenic (Average Nucleotide Identity (ANI)>99,99%)^33^, even when the isolates were obtained from distinct and geographically-separated *Hpa* cultures (Supplementary Fig. S4). Only the 2 *Microbacterium* isolates *(Microbacterium* f0c76-1 and *Microbacterium* f0c76-2, respectively) that represented HAM ASV f0c76 were found not to be isogenic (ANI = 88,5%, Fig S4). These isolates were both used in subsequent analyses.

We used these 9 distinct HAM bacterial genomes to investigate the presence of HAM bacteria in 7 additional independent *Hpa* cultures. To this end, we made use of 7 publicly-available *Hpa* genomes that had all been produced from spores of *Hpa*-infected plants and, thus, the underlying raw sequencing data essentially represent *Hpa* metagenomes of *Hpa* and its associated microbiome. These seven metagenomes were either obtained by Sanger sequencing of selected Bacterial Artificial Chromosome (BAC) clones derived from *Hpa* isolate Emoy2^34^ or by Illumina sequencing of the *Hpa* isolates Emoy2, Hind2, Cala2, Emco5, Emwa1, and Maks9^35^. They were all originally isolated in the UK and were maintained by regular transfer of *Hpa* spores from infected plants to a new batch of uninfected plants in laboratories located in East-Maling (now Warwick University)^34^ and Norwich (Sainsbury laboratory)^35^, respectively.

Reads from the *Hpa* metagenomes were pseudo-aligned^36^ to a genome index containing the 9 distinct HAM bacterial genomes and all available unique genomes (<98% ANI) of the 8 corresponding genera (in total 1128 genomes; Supplementary Table S4; Supplementary Fig. S5 – S13). In this way, the majority of bacterial genomes within the index were assigned only a few reads from every *Hpa* metagenome (considered background noise). Contrastingly, we generally observed a few genomes per genus that were assigned far more reads than the background noise (Fig S5 – S13). Using this method, we detected the presence of 7 out of 9 HAM bacterial genomes in multiple of the 7 *Hpa* metagenomes (Fig. 3A, Fig S5 – S13). Only the *Arthrobacter* and *Methylobacterium* HAM genomes were not detected in any of the *Hpa* metagenomes. In each of the investigated metagenomes, at least 2 of these 7 bacteria were detected and all 7 HAM bacteria were detected in the metagenome of the *Hpa* Hind2 isolate.

**Figure 3.**
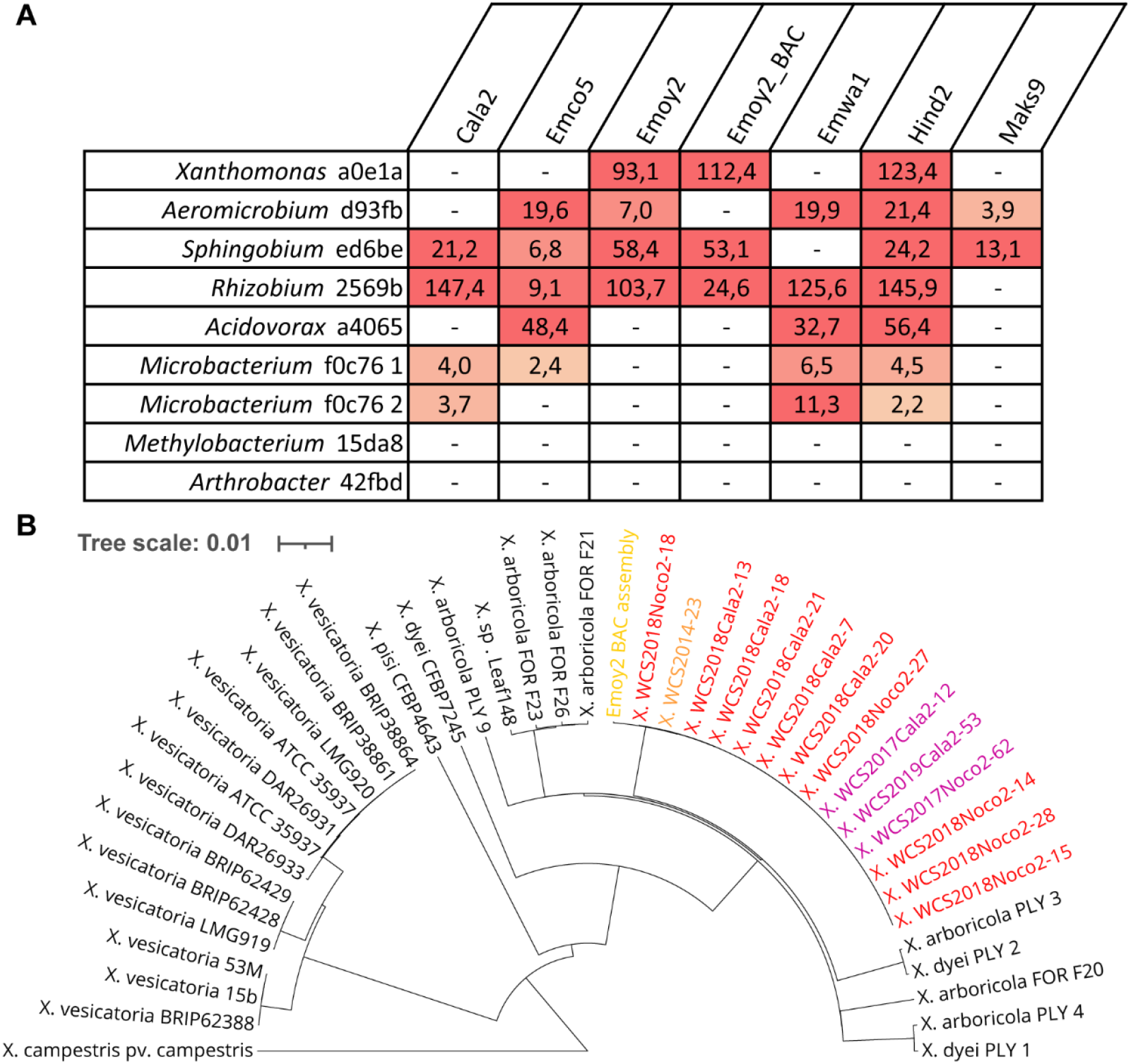
Isogenic HAM bacterial genomes are present in metagenomes of geographically separated *Hpa* cultures. (**A**) Heatmap indicating the presence of specific HAM bacterial genomes in publicly-available *Hpa* metagenomes. The numbers indicate signal-to-noise ratios, in which signal represents the number of reads that was assigned to a specific genome, and background noise was calculated as the total number of reads that were assigned to all genomes within a specific genus, divided by the number of genomes within that genus. Genomes with a signal-to-noise ratio below 2 were considered undetected (-). (**B**) Dendrogram based on UPGMA hierarchal clustering of a (1 – ANI) distance matrix for all *Xanthomonas* a0e1a isolates that were sequenced (red indicates isolates from Utrecht, purple indicates isolates from Cologne), *Xanthomonas* sp. WCS2014-23 (indicated in orange), a *Xanthomonas* sp. genome assembly using Sanger reads from an Emoy2-derived BAC-library (indicated in yellow), the 25 most related *Xanthomonas* genomes in the refseq database, and *Xanthomonas campestris* pv. *campestris* as outgroup. Tree scale represents branch length corresponding to the proportion of non-identical nucleotides between genomes.

The Emoy2 metagenome that was derived from a BAC library^34^ that was published in 2010 consisted of longer high-quality reads than the Illumina-based metagenomes^35^ and a large number of BAC-derived reads were assigned to the *Xanthomonas* a0e1a genome (Supplementary Fig. S13). We were able to partially re-assemble the *Xanthomonas* BAC clones from which these reads originated. This resulted in the assembly of 7 contigs ranging from 9844 bp to 62269 bp, with a total length of approximately 233 kb, and these contigs shared an ANI of 99,97% with the *Xanthomonas* a0e1a genomes from the Utrecht and Cologne *Hpa* isolates from this study (Fig. 3B, Supplementary Fig. S14). This confirms that the *Hpa* metagenome reads assigned to the *Xanthomonas* a0e1a genome are indeed derived from a bacterium that is isogenic to *Xanthomonas* a0e1a genomes isolated in this study. Intriguingly, the *Xanthomonas* a0e1a isolates were also isogenic to *Xanthomonas* sp. WCS2014-23 (Fig. 3B), previously isolated in our lab as part of a plant-protective consortium of microbes from the roots of downy-mildew infected plants^13^.

Together these results show that, although none of the investigated HAM bacteria are obligately associated with *Hpa*, different combinations of these HAM bacteria were always present in each of the 11 distinct *Hpa* cultures tested. Although the HAM comprises bacteria of diverse taxonomy, isogenic representatives of the HAM taxa are enriched in strictly separated *Hpa* cultures maintained in British, Dutch and German laboratories. These data suggest that independent cultures of distinct *Hpa* isolates have independently acquired similar consortia of isogenic HAM bacteria, and that *Hpa*-infected leaves thus selectively favor the recruitment and selection of specifically these bacteria.

### HAMs benefit from *Hpa* infection

We then questioned whether HAM enrichment on the infected leaves is driven by the interaction with *Hpa* or simply by an intrinsic ability of HAM members to outcompete other microbes in the Arabidopsis phyllosphere. To investigate HAM development in absence of *Hpa*, we made use of the gene *RESISTANT TO PERONOSPORA PARASITICA 5 (RPP5)*, which provides resistance to Noco2 in wild-type L*er* plants and in transgenic Col-0 *RPP5* plants^29^, rendering the latter resistant to both Noco2 and Cala2. Since Noco2 and Cala2 have distinct HAMs (Fig. 1), spore suspensions of these 2 cultures were mixed in equal proportions to make an *Hpa* inoculant with a uniform HAM (uHAM). This uniform inoculant was used to inoculate three lineages of plants: 1) wild-type Col-0 (resistant to Cala2, susceptible to Noco2, so microbial wash-offs from this lineage only contain Noco2 spores and microbiota from uHAM), 2) L*er* plants (resistant to Noco2, susceptible to Cala2, so microbial wash-offs from this lineage only contain Cala2 spores and microbiota from uHAM), and 3) transgenic Col-0 *RPP5* plants (resistant to both isolates, so wash-offs do not contain *Hpa* spores, only the microbiota from uHAM). One week after inoculation, leaf wash-offs from these 3 lineages of plants were prepared and used to inoculate L*er rpp5* plants^37^, which are susceptible to both Noco2 and Cala2 (Supplementary Fig. S15). Subsequently, the phyllosphere wash-offs from Lineage 1 (Noco2+uHAM), Lineage 2 (Cala2+uHAM only), and Lineage 3 (only uHAM) were passaged every week to a new population of susceptible L*er rpp5* plants, such that 8 independent phyllosphere cultures were passaged per lineage. This was repeated for in total 9 consecutive passages. Lineage 3 L*er rpp5* plants did not develop *Hpa* infections, indicating that the experiment remained free from cross-contamination. Untreated L*er rpp5* plants were included as an additional negative control (Supplementary Fig. S15). We analyzed the bacterial phyllosphere microbiomes from all 3 lineages and untreated plants at the end of the 1^st^, 5^th^ and 9^th^ passage on L*er rpp5* by 16S rDNA amplicon sequencing.

At each of these timepoints, phyllosphere microbiome communities of untreated controls were clearly distinct from the inoculated phyllosphere samples of Lineages 1-3 (Supplementary Fig. S16A, Fig. 4A, Supplementary Table S5; *P*<0.05 in PERMANOVA), suggesting the uHAM persists to a certain extent even in absence of *Hpa.* However, while uHAM microbiomes associated with either Noco2 or Cala2 were similar, they differed significantly from the *Hpa*-depleted uHAM communities (Fig S16B-D, Tables S5, S6). These *Hpa* effects were detectable after the first L*er rpp5* passage but were more evident after passages 5 and 9 (Supplementary Table S6, Supplementary Fig. S16B-D). This shows that *Hpa* infection significantly affects phyllosphere microbiome composition.

**Figure 4.**
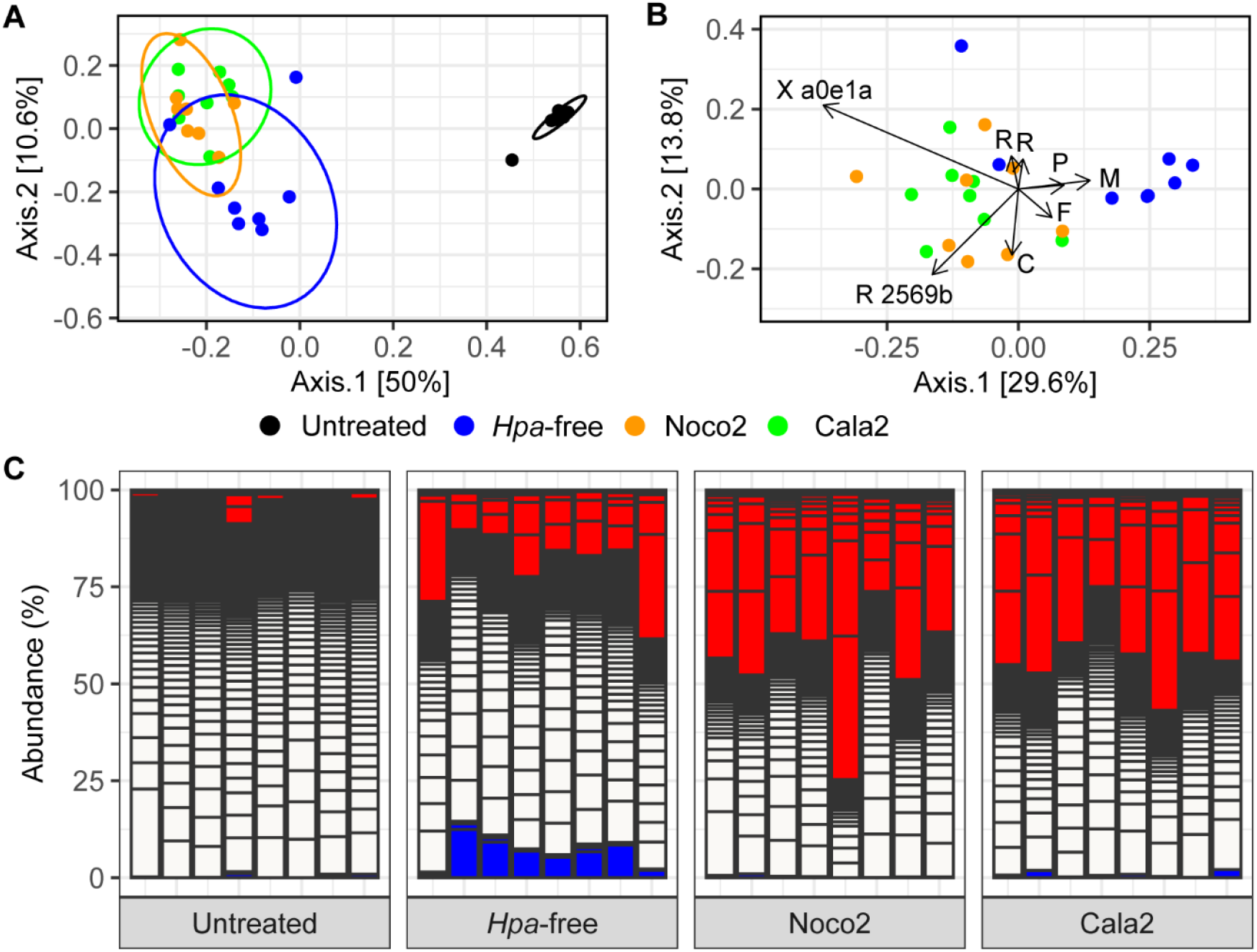
HAM ASV abundances diminish in absence of *Hpa.* (**A**) PCoA ordination plot based on Bray-Curtis dissimilarities of phyllosphere bacterial communities following 9 HAM passages over *Arabidopsis thaliana Ler/rpp5* plants in presence of *Hpa* isolate Noco2 (orange symbols) or Cala2 (green symbols), or in absence of *Hpa* (blue symbols). Black symbols show phyllosphere microbiomes of plants that were left untreated. Ellipses represent multivariate t-distributions with a 95% confidence level. (**B**) PCoA ordination biplot similar to (A), without untreated samples. Arrows indicate the relative contribution of individual ASVs to the first two principal coordinate axes. Displayed are the top five contributors for each axes (8 ASVs as 2 ASVs were top contributors to both axes). X a0e1a: *Xanthomonas* ASV a0e1a; R 2569b, *Rhizobium* ASV 2569; P, *Pseudomonas;* M, *Methylophilus;* F, *Flavobacterium;* C, *Chryseobacterium.* (**C**) Stacked bar chart with the relative abundances of the ASVs that are significantly enriched (red), depleted (blue), or unaffected (white) in Noco2- (Lineage 1) and Cala2-infected (Lineage 2) samples compared to *Hpa*-free cultures (Lineage 3). Each bar represents the ninth passage of an independent lineage of passages or an untreated sample (Lineage 4) grown at the same time.

Next, we focused specifically on passage 9 to further study the effect of the removal of *Hpa* from its associated microbial community (Fig. 4A). Interestingly, a PCoA biplot (Fig. 4B) shows that the previously-observed *Xanthomonas* ASV a0e1a is the strongest contributor to the separation of the Noco2- and Cala2-infected plants from the *Hpa*-free cultures, highlighting that this bacterium is most-strongly associated with downy mildew infection. Differential abundance testing revealed 18 ASVs that were enriched in the phyllosphere of infected plants compared to *Hpa*-free inoculated plants (Supplementary Table S7). Whereas the 18 enriched ASVs occupy on average 44% of the total phyllosphere communities of *Hpa*-infected plants, they represent approximately 20% of the communities in *Hpa*-free cultures, while they are largely absent in untreated control plants (Fig. 4C). Strikingly, the *Hpa*-infected-plant phyllospheres of passage 9 were dominated by *Xanthomonas* ASV a0e1a (Supplementary Fig. S17), whereas it diminished in the *Hpa*-free phyllospheres. Thus, the presence of *Hpa* benefits specific HAM bacteria in the phyllosphere, among which most prominently *Xanthomonas* ASV a0e1a.

### HAM bacteria are promoted by *Hpa* and form a resistobiome that reduces *Hpa* sporulation

We then wondered whether the abundance of individual HAM species on leaves would indeed increase as a result of infection with *Hpa.* To test this, we generated a gnotobiotic culture of *Hpa* isolate Noco2 (henceforth referred to as gno*Hpa*) by carefully touching healthy Arabidopsis seedlings, grown axenically on agar-solidified medium, with sporangiophores of *Hpa* extending from a detached and infected Arabidopsis leaf. After several weeks of subsequent careful passaging of *Hpa* in this gnotobiotic system, leaf wash-offs from this *in vitro* Arabidopsis-*Hpa* culture were plated on different cultivation media. No microbial growth was observed, indicating that the *gnoHpa* culture was cleared from its HAM.

Next, *gnoHpa* spores were co-inoculated with the individual HAM isolates (Fig. 3; Supplementary Table S8) or with 4 individual isolates that represent ASVs that were not enriched on *Hpa*-infected plants (hereafter: phyllosphere-resident isolates; Supplementary Table S8). These phyllosphere-resident isolates were obtained from Arabidopsis leaves simultaneously with the *Hpa*-associated isolates. We found that most of the HAM isolates reached significantly higher abundances when individually co-inoculated with *gnoHpa* (Fig. 5A). Particularly the core HAM isolate represented by *Sphingobium* ASV ed6be benefitted greatly from *gnoHpa* presence as its abundance increased significantly to more than 1000-fold in presence of gno*Hpa*. The phyllosphere resident isolates, however, were unaffected or even declined in abundance in the presence of *gnoHpa.* These results highlight that the HAM is specifically promoted in the phyllosphere by downy mildew infection.

**Figure 5.**
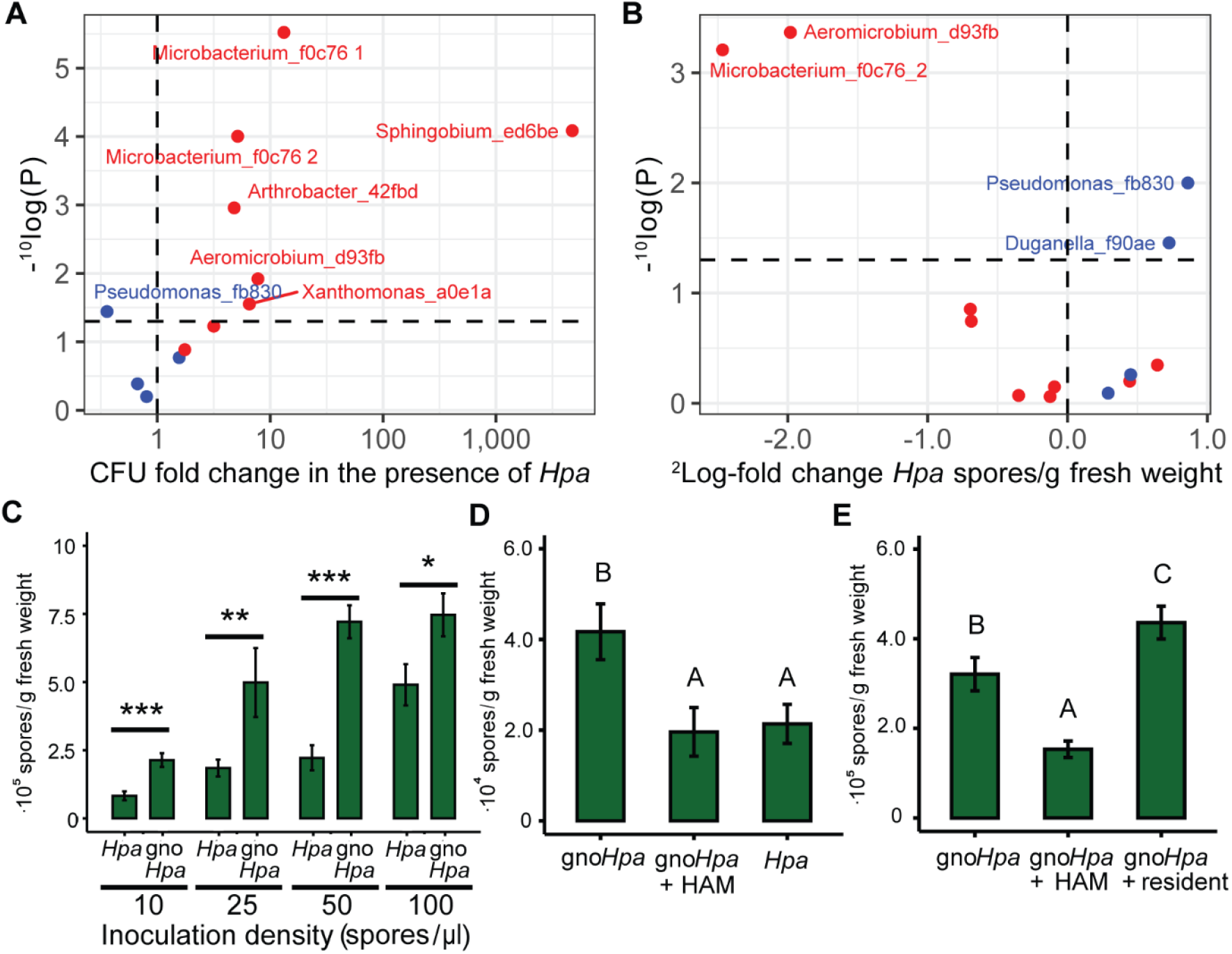
HAM bacteria benefit from the presence of *Hpa* in a gnotobiotic system and can reduce disease. Fold change in (**A**) bacterial abundance of HAM isolates 7 days after inoculation of each isolate separately on axenic plants with or without *gnoHpa* spores. (**B**) Fold change in *Hpa* spore production in the presence or absence of single bacterial isolates. Scatterplots show relation between fold change of *Hpa* spore production on gnotobiotic Arabidopsis plants growing on agar-solidified MS and log-transformed *P*-values (Wilcoxon tests). Red dots denote HAM bacterial isolates that represent *Xanthomonas* HAM ASV a0e1a, *Acidovorax* HAM ASV a4065, *Sphingobium* HAM ASV ed6be, *Arthrobacter* HAM ASV 42fbd, *Aeromicrobium* HAM ASV d93fb, *Methylobacterium* HAM ASV 15da8 and 2 isolates representing *Microbacterium* HAM ASV f0c76. Blue dots represent phyllosphere-resident isolates (Table S8). The vertical dashed lines separate negative (left) and positive (right) effects on (**A**) bacterial abundance and (**B**) *Hpa* spore production. The horizontal line indicates the significance threshold (*P* = 0.05) above which significant differences are displayed. Isolate names are only included in the plots if they are above this significance threshold.(**C**) Bar graph of *Hpa* and *gnoHpa* spore production 1-week post-inoculation with different inoculum spore densities. *N* = 10; error bars indicate standard error; *: *P* < 0.05; **: *P* < 0.01; ***: *P* < 0.001 in Student’s *t*-test. (**D**) Bar graph of *Hpa* spore production following inoculation of 14-day-old Arabidopsis plants with *gnoHpa* spores, *gnoHpa* amended with HAM filtrate, or regular *Hpa* spores suspensions. (**E**) Bar graph of *Hpa* spore production following inoculation of 14-day-old Arabidopsis plants with *gnoHpa* spores, *gnoHpa* amended with HAM filtrate, or *gnoHpa* amended with the microbiome filtrate of healthy plants (resident). Bars in D and E show the mean of >11 replicate pots. Error bars depict the standard error of the mean. Capital letters show significant difference (ANOVA with Tukey’s posthoc test).

In similar experiments we quantified *gnoHpa* spore production and observed that HAM-isolates *Aeromicrobium* d93fb and *Microbacterium* f0c76_2 significantly reduced *gnoHpa* spore production, whereas phyllosphere-resident isolates *Pseudomonas* fb830 and *Duganella* f90ae significantly promoted the production of gno*Hpa* spores (Fig. 5B). These results suggest that some HAM bacteria can aid the plant in reducing downy mildew disease. To further explore this, we quantified *Hpa* sporulation after inoculating Arabidopsis seedlings with *Hpa* or gno*Hpa* at multiple starting inoculum densities (Fig. 5C), with *gnoHpa* that was supplemented with either bacterial leaf wash-offs from *Hpa*-infected plants from which *Hpa* spores were removed through filtration (gno*Hpa* + HAM; Fig. 5D), or with similarly treated leaf wash-offs from healthy plants (gno*Hpa* +resident; Fig. 5E). Seven days after inoculation, HAM-free *gnoHpa* produced more spores compared to regular HAM-containing *Hpa* (Fig. 5C, D). Interestingly, supplementation of *gnoHpa* with HAM wash-offs reduced spore production to the same level as regular HAM-containing *Hpa* (Fig. 5D). When we compared the effect of HAM with that of phyllosphere resident microbiota from healthy plants on *gnoHpa* performance (Fig. 5E), we again observed a negative effect of HAM on *gnoHpa* spore production, while the phyllosphere resident microbiota yielded even higher sporulation than HAM-free *gnoHpa* on its own (Fig. 5E). Thus, HAM members reduce *Hpa* sporulation and thereby promote host health. We propose to call such a disease suppressive, pathogen-associated microbiome a ‘resistobiome’.

### Phyllosphere HAM resistobiome is recruited from the rhizosphere

Previously, we demonstrated that conditioning of soil by *Hpa*-infected Arabidopsis makes the next planting growing on that soil more resistant to *Hpa*^13^. This phenomenon is called a soil-borne legacy (SBL)^18^ and is associated with a shift in the rhizosphere microbiome that mediates an induced systemic resistance against *Hpa* in a next population of plants^13,38^. Interestingly, a number of the ASVs that became enriched in the rhizosphere upon foliar *Hpa* infection^13^ are also found in the phyllosphere HAM resistobiome in the present study. Therefore, we hypothesized that colonization of HAM bacteria is first promoted in the rhizosphere upon foliar infection with *Hpa*, resulting in the creation of a SBL. When a next population of plants subsequently germinates in SBL soil, their phyllospheres acquire the HAM bacteria, which upon foliar *Hpa* infection are further stimulated and become members of the HAM resistobiome in the phyllosphere. To test this hypothesis, we first set up a SBL experiment in which we conditioned soil with Arabidopsis Col-0 plants inoculated with mock, regular HAM-containing *Hpa* Noco2, or HAM-free *gnoHpa* Noco2, after which we quantified susceptibility of the successive (response) population of Col-0 plants on that soil to Noco2 (Supplementary Fig. S18). Interestingly, both conditioning of the soil with *Hpa*- and gno*Hpa*-infected plants reduced *Hpa* sporulation in a next population of *Hpa*-inoculated Col-0 plants (Fig 6A, Supplementary Fig. S19). This suggests that *Hpa* infection by itself, even without its associated HAM, is sufficient for the creation of a SBL. To verify this further we tested whether conditioning of the soil with *Hpa*-inoculated Col-0 *RPP5* plants, which are resistant to *Hpa* Noco2, would lead to the creation of a SBL. This was not the case (Fig. 6B), confirming that *Hpa* infection is required for the establishment of a SBL, and that only the introduction of HAM bacteria to the conditioning population of plants does not create a SBL.

**Figure 6.**
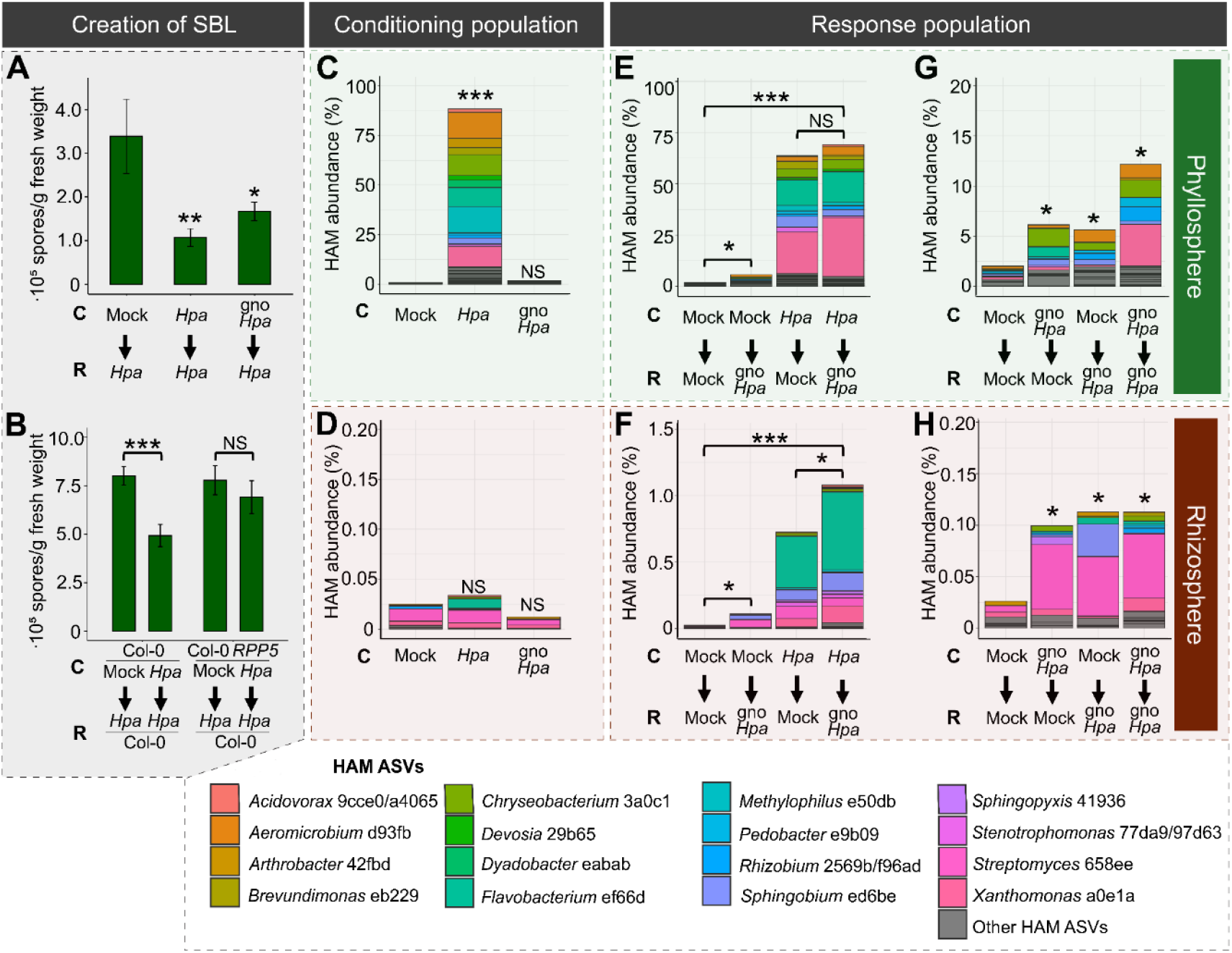
HAM ASVs are selectively promoted in response to *Hpa* infection and associated with SBL. (**A**) *Hpa* spore production on response (R) populations of Arabidopsis Col-0 plants growing on soil conditioned (C) by populations of Col-0 plants inoculated with either mock-, *Hpa*-, or gno*Hpa*-spore suspensions. Asterisks indicate significant differences compared to mock-conditioned plants in a Dunnet’s test. (**B**) *Hpa* spore production of Arabidopsis Col-0 plants growing in soil conditioned by populations of Col-0 or transgenic Col-0 *RPP5* plants that had been mock inoculated or inoculated with a *Hpa*-spore suspensions. Spore production was quantified 7 days post inoculation and normalized to shoot fresh weight. Asterisks indicate significance level in Student’s *t*-test compared to mock conditioned plants per plant genotype. (**C-H**) Cumulative relative abundance of 52 HAM ASVs in the phyllosphere (**C,E,G**) or rhizosphere (**D,E,F**) of a conditioning (**C,D**) or response population of Arabidopsis Col-0 plants. Colors indicate the taxonomy of 19 distinct HAM ASVs that together comprise the top-15 most-abundant HAM ASVs in the phyllosphere and rhizosphere. Colors correspond to single HAM ASVs, except for the genera *Acidovorax, Rhizobium* and *Stenotrophomonas* which are represented by 2 HAM ASVs. Asterisks indicate significance level in *fdr-corrected* Student’s *t*-test compared to mock-treated (**C**), or mock-treated mock-conditioned (**E-H**) plant populations. All bars and error bars indicate the average and standard error, respectively, of ≥ 10 replicate pots. *: *P* < 0.05; **: *P* < 0.01; ***: *P* < 0.001. NS: Not significant.

Next, we tested whether the HAM bacteria were promoted by *Hpa*-infected plants were subsequently picked up by a succeeding population of plants growing in the conditioned soil. To this end, we monitored the buildup of HAM bacteria in the rhizospheres and the phyllospheres of conditioning phase and response-phase Col-0 plants in the SBL experimental setup (Supplementary Fig. S18). As expected, one week after inoculation of the conditioning population of Col-0 plants with HAM-containing *Hpa*, we observed a significant shift in the phyllosphere bacterial community (Supplementary Fig. S20A; PERMANOVA: *R*^2^ = 0.58, *P* < 0.001), while upon inoculation with HAM-free *gnoHpa* we did not (Supplementary Fig. S20A; PERMANOVA: *R*^2^ = 0.05, *P* = 0.45). Using Deseq2^32^, we identified 52 ASVs that were uniquely enriched (*P* < 0.05, Supplementary Table S9) and together comprised 90% of the total bacterial community in the phyllosphere of *Hpa*-inoculated plants (Fig. 6C). The enriched ASVs were largely consistent with our earlier observations (Figs. 1, 2 and 4). In the remainder of this experiment these 52 ASVs are collectively referred to as HAM ASVs. The HAM ASVs were lowly abundant or below the detection limit in the phyllosphere of mock- or gno*Hpa*-inoculated plants (Fig. 6C) and in the rhizospheres of all conditioning-phase plants (Fig. 6D), confirming that in the conditioning phase they established in the phyllosphere as a result of co-inoculation with HAM-containing *Hpa.*

Next, we monitored the buildup of the HAM bacterial phyllosphere and rhizosphere communities of SBL response-phase Col-0 plants, one week after inoculation of this second population of plants with HAM-free *gnoHpa* or a mock-solution (Supplementary Fig. S18). Although no HAM bacteria were introduced during the inoculation of response-phase plants with *gnoHpa*, we found that the cumulative abundance of the 52 predefined HAM ASVs was significantly higher in the phyllospheres (Fig. 6E) and rhizospheres (Fig. 6F) of plants growing on *Hpa*-conditioned soils than on mock-conditioned soils. In the phyllosphere, *Xanthomonas* ASV a0e1a was among the most-dominant HAM ASVs that were transferred via the *Hpa*-condidioned soil to the response population of Col-0 plants (Fig. 6E), whereas in the rhizosphere *Flavobacterium* ASV ef66d most-dominantly increased in abundance (Fig. 6F). These results indicate that phyllosphere HAM bacteria are soil-borne and that they can readily colonize the phyllosphere from *Hpa*-conditioned soil.

Remarkably, we observed that in the response population of gno*Hpa*-inoculated Col-0 plants growing on mock-conditioned soil, the cumulative relative abundance of HAM ASVs was significantly higher in both the rhizosphere and phyllosphere in comparison to mock-inoculated response-phase plants (Fig. 6E-F). Because in these *gnoHpa* treatments HAM bacteria were never introduced with the inoculum, we concluded that gno*Hpa*-inoculated plants indeed specifically recruited and promoted the abundance of HAM bacteria in both the rhizosphere and phyllosphere. To further examine this, we focused on all treatments in this experiment to which the HAM was never co-inoculated. The cumulative relative abundance of HAM ASVs was significantly higher in both the rhizospheres (Fig. 6G) and the phyllospheres (Fig. 6H) of response-phase plants that either grew on gno*Hpa*-conditioned soil or were themselves inoculated with gno*Hpa*. These results further support the notion that downy mildew-infected plants selectively recruit HAM bacteria from the rhizosphere to the phyllosphere, where they are subsequently further promoted by downy mildew infections.

### *Aeromicrobium* ASV d93fb and *Xanthomonas* ASV a0e1a are important contributors to the resistobiome

To investigate which of the individual HAM ASVs were promoted by *gnoHpa* infection, we compared the abundance of ASVs between gno*Hpa*-inoculated plants on gno*Hpa*-conditioned soil to mock-inoculated plants on mock-conditioned soil. Remarkably, 10 HAM ASV were among the top 30 most strongly enriched ASVs in the phyllosphere of gno*Hpa*-infected plants (Supplementary Fig. S21). This again confirms the specific recruitment of these HAM ASVs by these diseased plants. Remarkably, the 4^th^ most-strongly responding ASV is *Xanthomonas* HAM ASV a0e1a, which increased 17-fold in phyllosphere abundance and 2.7 fold in the rhizosphere of gno*Hpa*-infected plants. Previously, we showed that *Xanthomonas* sp. WCS2014-13, representative of HAM ASV a0e1a, can contribute to suppression of *Hpa*^13^. Although the enrichment of the predefined HAM ASVs was evident, the increase of most HAM ASVs was not statistically significant as a result of their irregular occurrence and consequential low statistical power. In the rhizosphere, ANCOM-BC^39^ identified only 2 ASVs that were significantly enriched (*P* < 0.05, *fdr*-corrected) following infection, but both were lowly abundant and not predefined as HAM ASVs. Similarly in the phyllosphere, the enrichment following infection of only 1 of 452 phyllosphere ASVs was deemed statistically significant by ANCOM-BC. However, this singled significantly responding ASV was *Aeromicrobium* HAM ASV d93fb (Supplementary Fig. S22). Extraordinarily, the isolate representing specifically this HAM ASV consistently reduced spore production when co-inoculated with gno*Hpa* on axenic Arabidopsis plants (Fig. 5B). The observed phyllosphere enrichment of *Aeromicrobium* ASV d93fb resulting from *gnoHpa* infection, and also that of *Xanthomonas* ASV a0e1a, could thus be sufficient to explain the increased resistance associated with SBL. Together these results again highlight that HAM ASVs are specifically promoted as a result of *gnoHpa* infection and suggest that the increased resistance of SBL results from the increased abundance of HAM bacteria and their combined actions as a resistobiome.

## Discussion

In recent years, evidence has been mounting that upon perceiving environmental stress, plants actively shape their microbiome to recruit microbes that help alleviate such harmful conditions^18^, as has been reported for plants responding to nutrient deficiency^40,41^, drought^42–44^ or salinity^45^, but also to stress caused by microbial pathogens^13,14,20,46,47^. The disease-induced recruitment and buildup of beneficial microbes is thought to result in the creation of disease-suppressive soils^18–21,48,49^.

In this study, we investigated whether infection by a foliar pathogen similarly leads to changes in the phyllosphere microbial community, either to the benefit of the pathogen or the plant. We found that laboratory cultures of distinct isolates of the obligate downy mildew pathogen *Hpa*, strictly separately maintained in laboratories in Germany, the Netherlands, and the U.K., are dominated by a similar set of isogenic *Hpa*-associated bacteria. Most prevalent among these phyllosphere HAM bacteria is a *Xanthomonas* sp. that was found in 7 out of 11 *Hpa* cultures, but isogenic representatives of at least 6 out of 9 HAM isolates were frequently represented in the *Hpa* cultures that we investigated. Our results show that a HAM community is selectively promoted in the phyllosphere of *Hpa*-infected plants, and that this can be reconstructed on axenic Arabidopsis plants on which HAM-free *gnoHpa* and individual HAM isolates are co-inoculated. Interestingly, inoculation of Arabidopsis plants growing on natural soil with HAM-free *gnoHpa* resulted in increased relative abundances of HAM ASVs in both the rhizosphere and phyllosphere of infected plants. We provide evidence that HAM bacteria are recruited from the soil surrounding downy mildew-infected plants to the phyllosphere of a next population of plants growing in the conditioned soil, where they are subsequently further promoted by *Hpa* infection. The fact that similar bacterial taxa are member of HAM communities in physically and geographically separated *Hpa* cultures suggests that these specific HAM members are independently selected from different soils by Arabidopsis in response to *Hpa* infection. In free analogy of the Baas-Becking hypothesis *“Everything is everywhere, but the environment selects”*^50^, it thus appears that HAM bacteria are everywhere, but the infected plant selects.

While HAM bacterial growth is promoted on *Hpa*-infected leaves, vice versa *Hpa* spore production is reduced by the HAM or specific individual HAM isolates. Because the HAM limits *Hpa* infection, we coin the term “resistobiome” for this pathogen-associated microbiome. Moreover, foliar *Hpa* infection can stimulate a SBL in the soil that results in increased resistance to *Hpa* of a subsequent population of plants growing on the same soil. This increased resistance also coincides with an increased cumulative abundance of HAM ASVs in the rhizosphere and phyllosphere of these plants. As application of HAM isolates to leaves (this study) or to roots^13^ is sufficient to reduce *Hpa* spore production, it is likely the HAM is responsible for the increased protection that results from SBL. The mechanism by which this HAM either directly or indirectly protects against *Hpa* should be further investigated. However, we previously found that mutant plants impaired in defense signaling that involves the plant hormone salicylic acid, are not protected by a *Hpa*-induced SBL, suggesting the SBL and associated HAMs work at least partly through activation of salicylic acid-dependent immunity^38^.

In sum, our data show that subsequent populations of Arabidopsis plants that are infected by Arabidopsis’ cognate downy mildew pathogen develop disease-associated microbiomes in both the phyllosphere and the rhizosphere. This resistobiome hinders pathogen development and is thus the opposite of a pathobiome^25,26^, which promotes pathogen infection. Our results suggest that it is not the mere presence of the pathogen that leads to resistobiome assembly. The creation of a SBL requires successful infection with the pathogen, as resistant plants that prevent *Hpa*-infection did not lead to a SBL. Moreover, we previously reported that foliar *Hpa* infection changes the root exudation profile, and that mutant plants impaired in the biosynthesis of root-secreted coumarins are not able to create a SBL even though they are just as susceptible as the wild type^38^. This indicates that it is the plant that actively assembles its resistobiome in response to attack. Future research should further elucidate the plant genetic mechanisms by which plants in this way ‘cry for help’. Fundamental understanding of such mechanisms would allow the breeding of crop varieties that enhance the build-up and protective function of resistobiomes. Ultimately this could contribute to sustainable agriculture that relies less on chemical inputs for crop resilience.

## Online Methods

### Plant materials and growth conditions

In this study, we used the Arabidopsis accessions Col-0, L*er*, C24^30^ and Pro-0^31^, transgenic Col-0 *RPP5* plants^29^ and the mutants L*er rpp5*^37^ and *eds1*^4,51^, respectively. For the experiments performed at UU that correspond to results shown in Fig. 1, 2, 4, 60-mL pots were filled with approximately 90 g of a mix of river sand and potting soil (5:12) supplemented with 10 mL half-strength Hoagland solution^52^ and further saturated with water. Twenty-one seeds were sown on top of the soil using toothpicks (Fig. 1, 2, 3) or denser fields of seedlings were sown through gentle dispersion from seed bags (9-passages experiment, Fig. 4). Pots withs seeds were subsequently submitted to stratification for three days in the dark at 4 °C. Seeds were then allowed to germinate and develop in a climate-controlled chamber (21 °C, 70% relative humidity, 12 h light/12 h dark, light intensity 100 μmol m^-2^ s^-1^). For the experiments at MPIPZ, Cologne, seeds were stratified for three days in the dark at 4 °C and sown on water-saturated Jiffy pellets (J-7) and allowed to develop at slightly shorter day cycles (10 h light/14 h dark). For experiments in which multiple genotypes of Arabidopsis were used simultaneously, seeds were surface sterilized before sowing by vapor-phase sterilization as described previously^53^.

For bioassays with both *Hpa* and gno*Hpa* and for SBL experiments (Fig. 5,6), a natural soil from the Reijerscamp nature reserve (52°01’02.55”, 5°77’99.83”) in the Netherlands^13^ was used. The soil was air dried and sieved twice (1×1 cm^2^) to remove rocks and plant debris. One day prior to sowing, the soil was watered in a 1:10 v/w ratio. Pots with a volume of 60 mL were filled with 120 g of soil (+/- 2.5 g), placed in 60-mm Petri dishes and the soil surface was covered by a circular cut-out of plastic micro pipette tip holders (Greiner Bio-One, 0.5-10 μL, Item No: 771280) to prevent growth of algae and to ensure that seeds are equally distributed during sowing^54^. Pots were stored at 4°C overnight before sowing. Arabidopsis accession Col-0 seeds were suspended in 0.2% (w/v) agar solution and stratified in dark conditions at 4°C for 48-72 hours. Seeds were sown by pipetting two or three seeds per hole of the plastic cover, resulting in approximately 30 seeds per pot. After sowing, pots were stored in trays with transparent lids and put in a growth chamber (21°C, 70% relative humidity, 10h light/14 h dark, light intensity 100 μmol m^-2^ s^-1^). Pots were watered two times a week with 3 mL water. After one week, closed lids were replaced with mash lids to reduce humidity and plants were once watered with 5 mL of ½-strength Hoagland nutrient solution.

### Preparation of *Hpa* spore suspensions and inoculation of plants

*Hpa* spore suspensions were prepared by harvesting shoot material 7-14 days after inoculation with spore suspensions of *Hpa* isolate Noco2^29^ or Cala2^28^ and by vigorously shaking the material in 20 mL of autoclaved tap water. Spores were counted in 3 separate 1-μL droplets using a transmitted light microscope (Carl Zeiss Microscopy, Standard 25 ICS, Item No. 450815.9902) and subsequently the spore suspensions were diluted to obtain appropriate spore densities as specified below.

For the experiment performed in Utrecht that corresponds to data shown in figures 1 & 2, 10-day-old Arabidopsis seedlings were spray-inoculated with spore suspensions in autoclaved tap water containing *Hpa* isolate Noco2 or Cala2 (50 spores/μL), a mix of both spore suspensions (50 spores/μL of each isolate), or mock-treated with autoclaved tap water. In Cologne (experiment corresponding to Fig. 2), 16-day-old Arabidopsis seedlings were spray-inoculated with Noco2 or Cala2 spores in MilliQ water or mock-inoculated with MilliQ. In bioassays and SBL experiments (Fig. 5,6), 14-day-old seedlings were spray-inoculated with *Hpa* or *gnoHpa* spore suspensions in tap water (Fig. 5C: 10, 25, 50, 100 spores/μL, Fig. 5D,E: 25 spores/μL, Fig. 6A: 50 spores/μL, Fig. 6B: 85 spores/μL, respectively). Pots with inoculated plants were airdried and randomly placed in trays in a climate chamber (16 °C Fig. 1-4, 21°C Fig. 5,6, 10 h light/14h dark, light intensity 100 μmol m^-2^ s^−1^) and were covered with moisturized transparent lids to increase humidity.

### Arabidopsis sample collection and disease quantification

Seven days after inoculation with *Hpa* spore suspensions, infected Arabidopsis shoot material was collected in 2-mL Eppendorf tubes (Utrecht, Fig. 1,2, Fig. 4, 6) or 15-mL conical tubes (Cologne, Fig. 2) containing 3 glass beads, carefully avoiding the sampling of roots or soil.

For DNA isolation, the material was immediately snap-frozen in liquid nitrogen and stored at −80 °C until further processing. Rhizosphere samples were taken by gently sieving the pots under running tap water until the loosely adhering soil was washed away and only roots with attached soil were left. For bulk soil samples, the upper soil layer (+/- 2 cm) was removed and the exposed soil was sampled. All sample types were collected in 2-mL Eppendorf tubes, snap frozen in liquid nitrogen, and stored at −80°C until further processing.

For disease quantification, the weighted shoot material was suspended in 3-6 mL of water depending on fresh weight and level of observed sporulation. Greiner tubes were vortexed for 15 s and spores were counted in 3 separate 1-μL droplets using a Transmitted Light Microscope (Carl Zeiss Microscopy, Standard 25 ICS, Item No. 450815.9902) and the average spore count was normalized by shoot fresh weight.

### Genomic DNA (gDNA) extractions

Total gDNA was extracted from Arabidopsis leaves and resident microbiomes using the PowerLyzer PowerSoil DNA isolation kit (Qiagen) modified for leaf material. In brief, frozen leaf samples were mechanically lysed twice for 60 s at 30 Hz using the Tissuelyser II (Qiagen). Per sample, 750 μL Powerbead solution and 60 μL C1 solution were added, samples were mixed by inverting tubes and subsequently incubated for 10 min at 65 °C. Total solutions were then transferred to Powerbead tubes and submitted to bead beating, twice, for 10 min at 30 Hz. Samples were then centrifuged for 4 min at 10.000x *g* and supernatant was transferred to new 2-mL collection tubes. The protocol was subsequently completed without further modifications according to the manufacturer’s instructions. DNA extractions for phyllosphere samples from the SBL experiment were performed using the QIAGEN MagAttract PowerSoil DNA KF Kit® and a ThermoFisher KingFisher® with the same modifications for leaf material as described above. Rhizosphere and bulk soil DNA was extracted according to the protocol provided by the QIAGEN MagAttract PowerSoil DNA KF Kit®. All DNA concentrations were quantified using a NanoDrop 2000®.

### *Hpa* quantification by qRT-PCR

*Hpa* was quantified from gDNA obtained from *Hpa*-infected and uninfected Arabidopsis shoot material according to Anderson *et al.*^55^. Two-step qRT–PCR reactions were performed in optical 96-well plates with a ViiA 7 real time PCR system (Applied Biosystems), using Power SYBR® Green PCR Master Mix (Applied Biosystems) with 0.8 μM primers (Supplementary Table S10). A standard thermal profile was used: 50 °C for 2 min, 95 °C for 10 min, 40 cycles of 95 °C for 15 s and 60 °C for 1 min. Amplicon dissociation curves were recorded after cycle 40 by heating from 60 to 95 °C with a ramp speed of 1.0 °C ^-1^. *Hpa* abundance was calculated by 2-^(*CtHPaACTIN–CtArabidopsisACTIN*)^.

### 16S rDNA amplicon library preparation for microbiome analysis

For the amplicon sequencing of the experiment in Fig. 1, library preparations for Illumina 16S rDNA amplicon sequencing were performed on gDNA samples using standard materials and methods as described in the Illumina protocol. For amplicon sequencing of experiments corresponding results shown in Fig. 2,4, we used primers with heterogeneity spacers for the 16S amplicon PCR^56^. This change was implemented to increase nucleotide diversity in the first 25 bases as analyzed by MiSeq and lower the required amount of PhiX^56^ spike-in, thereby increasing read depth per sample (Supplementary Table S10). All 16S rDNA library preparations were performed with the addition of PNA PCR clamps to the PCR1 reaction mixtures, at 0.25 μM per reaction, to prevent the amplification of plant-derived sequences^57^. ITS2 amplicon libraries were prepared in similar fashion, using primers^58^ and blocking oligonucleotides as specified in Supplementary Table S10, with altered numbers of PCR cycles (PCR1, 10 cycles; PCR2, 25 cycles), to accommodate the use of the ITS2 blocking oligonucleotides for the prevention of plant-derived reads^59^. Sequencing was performed with MiSeq V3 chemistry (2×300 base pairs paired-end) at the Utrecht Sequencing Facility (USEQ; Utrecht, the Netherlands, Figs. 1-2, Fig. 4)

For amplicon sequencing of the SBL experiment (Fig. 6), library preparation and sequencing with NovaSeq chemistry (2×250 base pairs paired-end sequencing) was performed by Genome Quebec (Quebec, Montreal, Canada). The primers used were 16S-B341F (5’-CCTACGGGNGGCWGCAG) and 16S-B806 (5’-GACTACHVGGGTATCTAATCC) according to Genome Quebec’s standard operating protocols. Plastid- and mitochondrial-blocking peptide nucleic acid (pPNA - 5’-GGCTCAACCCTGGACAG; mPNA - 5’-GGCAAGTGTTCTTCGGA) were used in the PCR reactions to prevent amplification of plant-derived sequences.

### 16S rDNA and ITS2 amplicon sequencing analyses

Preprocessing of sequencing data was performed in the Qiime2 environment (version 2019.7)^60^, using Cutadapt^61^ to remove primer sequences from reads and DADA2^62^ for quality filtering, error-correction, chimera removal, and dereplication to ASVs. Samples from the SBL experiment (Fig. 6) were sequenced twice and the resulting two datasets were merged after cutadapt- and DADA2-processing. ASV id’s were assigned based on the sequence-specific MD5-sums using the --p-hashed-feature-ids parameter, of which we used the first 5 characters in the text above to designate the individual ASVs. For 16S rDNA data sets, taxonomic assignment was performed using the VSEARCH plugin and the SILVA database (QIIME compatible 132 release, 99% clustering identity, 7-level RDP-compatible consensus taxonomies) from which the 16S V3/V4 regions were extracted based on the 16S rDNA specific primer sequences used in library preparation. Plant-derived sequences were identified and removed based on the annotation of “D_4 Mitochondria” at the family level or “D_2 Chloroplast” at the class level. Additionally, ASVs with unassigned taxonomies were removed. In the experiments sequenced with the MiSeq platform, ASVs in the lowest 1% of total cumulative abundances were removed. For the sequencing data generated with the NovaSeq platform, ASVs that contribute to the lowest 3% of total cumulative abundance or that were detected in less than 5 samples were removed from the data. Fungal ITS amplicons were taxonomically classified using a fitted classifier trained on the UNITE fungal database (QIIME release, version 7.2, 99% similarity clustering, 7-level taxonomies). ITS sequences (ASVs) were filtered based on minimum occurrence in two samples (Supplementary Fig. S2). Samples with less than 500 reads were removed from the dataset.

All beta-diversity-related calculations, graphs and differential abundance tests were performed in R (version 4.0.3). All PCoA ordinations and PERMANOVA tests were performed on Bray-Curtis dissimilarity matrices calculated for relative abundance data, using either vegan (version 2.5.7) or vegan functionalities within the phyloseq package (version 1.34.0). The pairwiseAdonis package (version 0.0.1) was employed for PERMANOVA tests involving multiple comparisons. For the SBL experiment (Fig. 6), rhizosphere samples, R-G-G1-4 and R-G-G1-8, clustered away from the other rhizosphere samples towards bulk soil samples. Hence, these samples were considered as not representative for the rhizosphere and removed for downstream analysis. Differential abundance testing was performed with either DESeq2^32^ (version 1.30.1) or ANCOM-BC^39^, employing a separate function to calculate the geometric mean used for the estimateSizeFactor step, namely: function(x, na.rm=TRUE) exp(sum(log(x[x > 0]), na.rm=na.rm) / length(x)) (described here: https://bioconductor.org/packages/devel/bioc/vignettes/phyloseq/inst/doc/phyloseq-mixture-models.htmL). This circumvents errors related to the sparsity in 16S rDNA amplicon data. Dendrogram visualization for Fig. 3B was achieved using ggtree (version 2.4.1). All other graphs were made using ggplot2 (version 3.3.3) or ggpubr (version 0.4.0). Data wrangling was done with packages from the Tidyverse suite.

### Bacterial isolate collection

Infected shoot material was harvested from Noco2 and Cala2-infected plants in Utrecht in 2018 and from Cologne in 2017 and 2019. Shoot material of *Hpa*-infected plants was stored in 1 mL of 10 mM MgSO_4_ with 25% glycerol (v/v) at −80 °C. For isolation, shoots were defrosted at room temperature, and plants were crushed using sterile pestles. In order to maximize the isolation of culturable bacterial species, a dilution series of the Utrecht samples was plated on agar-solidified medium containing either 1/10 strength tryptic soy broth (TSB; Difco), King’s medium B^63^ amended with 13 mg/L chloramphenicol and 40mg/L ampicillin (KB+), Luria Bertani (LB, Difco), R2A (Difco), yeast-extract mannitol (YEM, per L: 0.5 g yeast extract (Difco), 5 g mannitol, 0.5 g K2HPO4, 0.2 g MgSO_4_.7H_2_O, 0.1 g NaCl (Difco), pH 7.0), Nutrient agar (NA, per L: 5 g peptone (Difco), 3 g beef extract (Difco), pH 6.8); and glucose nutrient agar (GNA, per L: 5 g peptone, 3 g beef extract (Difco), 10 g glucose (Difco). All media were amended with 200 μg/mL Delvocid (DSM; active compound: natamycin) to prevent fungal growth. Plates were incubated at 21 °C for 2 – 11 days. A total of 654 bacterial colonies, selected on morphology and/or time of emergence, were streaked on the same type of agar medium from which it was selected. Single colonies from pure cultures were inoculated in 1/10 strength tryptic soy broth (TSB; Difco), incubated overnight at 20 °C at 180 rpm, and stored at −80 °C in 25% (v/v) glycerol. Pure cultures were labeled based on the sample it originated from, the type of medium it was isolated from, and numbered according to the order of selection on that medium. Similar methods were applied to isolate Xanthomonads from the Cologne samples, but only 1/10 strength TSA was used and 48 colonies were selected for yellow color of colonies. Isolates that were used in further experiments and analyses were given a unique code and absorbed in the Willy Commelin Scholten collection WCS (Supplementary Fig. S4).

### Initial characterization of bacterial isolates

All isolates were processed for simultaneous 16S rDNA sequencing using Illumina MiSeq V3 chemistry, using the multiplexing strategy described by De Muinck *et al.*^56^. All isolates were grown for 2 days in 1/10^th^ strength TSB and 1 μL of each culture was added directly to a PCR reaction in 96-well plates, containing 0.2 μM of column-specific forward primers and row-specific reverse primers (Supplementary Table S10), and 2 KAPA HiFi Hotstart Ready Mix (Roche) to a total volume of 15 μL. PCR was performed by 10 min incubation at 95 °C followed by 30 cycles of 30 s at 95 °C, 30 s at 55 °C and 30 s at 72 °C, followed by a final elongation step of 5 min at 72°C. PCR products were purified using AMPure XP beads with 9 μL of bead solution per 15 μL PCR mixture and washing with 80% ethanol. PCR products were quantified using Nanodrop 2000 spectrophotometer (Thermo Fisher Scientific), concentrations were normalized to 1 ng/μL and each 96-well plate of samples was combined into a pooled sample. Per pooled sample, 1 μL was then submitted to a second PCR reaction, wherein each pooled sample was assigned its unique pair of Nextera indexing primers (Supplementary Table S10). These PCR-reactions were performed with 2x KAPA HiFi Hotstart Ready Mix and 0.4 μM forward and reverse primers, in total volumes of 25 μL. The PCR-products were purified as described above, quantified using the Qubit dsDNA BR Assay kit according to the manufacturer’s instructions, normalized to 1 ng/ μL and again pooled together into a single 16S library. The 16S library was submitted to sequencing at USEQ. Between-plate demultiplexing was performed by USEQ, whereas within-plate demultiplexing and removal of adapters was achieved using Cutadapt^61^. Sequence data per bacterial isolate was then processed in the Qiime2^60^ environment using DADA2^62^ as described above. The resulting ASVs were matched to ASVs from the 16S rDNA analyses of bacterial communities based on their sequence specific MD5-sum identifiers.

### Whole-genome sequencing of bacterial isolates

Whole-genome sequencing (WGS) of bacterial isolates was performed on gDNA that was extracted using the GenElute Bacterial Genomic DNA Kit according to the manufacturer’s instructions. gDNA samples were processed for WGS at the Microbial Genome Sequencing Center (Pittsburgh, USA) and sequenced to an estimated coverage of 60x using 2×150 paired-end sequencing. Sequences from paired FASTQ files were quality filtered using Trimmomatic^64^ (version 0.39) with LIDINGWINDOW:4:20 as the only set parameter. Genomes were then assembled using Spades^65^ (version 3.11.1) with ‘–careful’ as the only set parameter. Assemblies were checked using Quast^66^. Contigs shorter than 1000 bp were then removed using the Galaxy webserver (https://usegalaxy.eu) using the ‘filter fasta’ function. If genomes for multiple isolates per ASV were obtained, their average nucleotide identities were calculated using the python3 module pyani. Dendrograms based on whole genomes were calculated using mashtree and visualized using the ITOL webserver (https://itol.embl.de/).

### Analysis of publicly available *Hpa* metagenomes

Sequence data described in the publication by Baxter *et al.*^34^, that was obtained from Emoy2 spores collected in water (similar to the *Hpa* inocula used in the present study), were obtained in fasta format from the NCBI TRACE archive using query ‘species_code=‘HYALOPERONOSPORA PARASITICA’’. The first and the last 100 base pairs of all reads were removed using Trimmomatic, resulting in reads with an average length of approximately 600 bp.

Sequence data described in the publication by Asai *et al.*^35^ was obtained from the European Nucleotide Archive under project number PRJEB22892. These reads we filtered based on quality using Trimmomatic, using; SLIDINGWINDOW:4:20 and MINLEN:45 for fastq files with 2 x 60 bp and 2 x 75 bp reads, and SLIDINGWINDOW:4:20 MINLEN:30 for fastq files with 2 x 35 bp reads.

To quantify specific genomes of interest within their respective genera, first we obtained non-redundant genomes of genera of interest, using bacsort (https://github.com/rrwick/Bacsort), which picks the best genome assembly from clusters of genomes that are within 98% average nucleotide identity (with parameter ‘--threshold 0.02’) of each other. Kallisto indices^36,67^ were then built using all these non-redundant genomes, the genomes of the bacteria identified and sequenced in the study, the *Hpa* Emoy2 genome, and the *Arabidopsis thaliana* genome. All reads per *Hpa* metagenome were then pseudo-aligned^36^ against these indices, and the proportion of reads that pseudo-aligned to the genome of interest among all reads mapped to genomes within that genus was calculated (Supplementary Fig. S5 – S12). Signal-to-noise ratios were calculated (Fig. 3A), in which signal represents the number of reads that were assigned to a specific genome, and noise was calculated as the total number of reads that were assigned to all genomes within a specific genus, divided by the number of genomes within that genus (as included in the genome index).

### Nine-passages experiment

Ten pots with ten-day-old seedlings of Arabidopsis accessions Col-0 and L*er*, and transgenic Col-0 *RPP5*^29^, kindly provided by Jane Parker (Max Planck Institute for Plant Breeding Research, Cologne, Germany) were inoculated with a mix of Noco2 and Cala2 spore suspensions, prepared as described above. These pots were placed in one tray covered with a transparent plastic lid, which was sprayed with water on the inside to raise humidity inside the tray, and placed in a climate chamber (16 °C, 10 h light/14 h dark, light intensity 100 μmol m^-2^ s^-1^). *Hpa* started to sporulate on Col-0 and L*er* plants approximately 5 days after inoculation.

After 7 days, leaf wash-offs were obtained from each pot (30 pots in total) and used to spray-inoculate a population of 10-day-old L*er rpp5*^37^ (also kindly provided by Jane Parker, Cologne) plants growing on 60-mL pots, each of which was placed in an individual Eco2box. After 7 days, and visual and microscopic confirmation that sporulation only occurred in plants inoculated with Noco2 or Cala2, half of the seedlings per pot were collected, snap frozen in liquid nitrogen, and stored for further processing. The remaining plants from each pot were used to generate leaf wash-offs in 5 mL of autoclaved water, of which approximately 600 μl was spray-inoculated onto a new population of 10-day-old L*er rpp5* seedling using 2-mL spray units, such that the leaf wash-off from one pot was used to inoculate the plants on one new pot only. These pots were again placed in new individual Eco2boxes to prevent cross-contamination and ensure the propagation of separated phyllosphere cultures. In total this process was repeated 8 times, thus passaging the *Hpa*-associated microbiome over 9 consecutive L*er rpp5* plant populations in presence of either Noco2 (Lineage 1), Cala2 (Lineage 2), or neither of the *Hpa* isolates (Lineage 3). For each plant population in the 9-passages experiment, 8 additional replicate pots with L*er rpp5* plants were left untreated and harvested as untreated controls. A schematic overview of this experiment is presented in Supplementary Fig. S15.

### Creation of a gnotobiotic *Hpa* culture

Microbial contaminant-free *Hpa* (gno*Hpa*) was generated by successive passaging of *Hpa* isolate Noco2 to susceptible Arabidopsis Col-0 seedlings grown on Murashige & Skoog (MS; Duchefa Biochemie)^68^ agar-solidified medium without sucrose. Vapor-phase sterilized^53^ Col-0 seeds were sown on MS agar medium. After 2 days of stratification at 4 °C, Petri dishes were placed vertically in a growth chamber (21 °C, 70% relative humidity, 12 h light/12 h dark, light intensity 100 μmol m^-2^ s^-1^). Ten-day-old seedlings were then inoculated with *Hpa* (Noco2) by gently touching leaves of the new host plants with sporangiophores extending from *Hpa*-infected leaves in a sterile-laminar-flow cabinet. This initial infection of axenically grown Col-0 seedlings was performed using *Hpa*-infected leaves from the standard (microbe-rich) Utrecht laboratory culture of Noco2. Petri dishes with infected seedlings were then placed in a growth chamber with optimal conditions for *Hpa* infection (16 °C, 10 h light/14 h dark, light intensity 100 μmol m^-2^ s^-1^). Using the same gentle touch-inoculation method, this *Hpa* culture was then passaged weekly to new axenically grown Col-0 seedlings on MS agar-solidified medium (stratified, germinated and grown as described above). After *Hpa* touch inoculations, newly infected axenically grown plants were placed in a growth chamber (16 °C, 10 h light/14 h dark, light intensity 100 μmol m^-2^ s^-1^). After each disease cycle, absence of microbial contaminants was tested by serial dilution plating on 1/10 TSA. Following the observation that there were no culturable microbes present in the gnotobiotic culture, which required 3 passages, we double checked the gnotobiotic nature of these cultures by amplicon sequencing the 16S rRNA gene as described above (data not shown). Upon verification of the absence of microbial contaminants, weekly passaging was performed on axenically grown hyper-susceptible *eds1* (enhanced disease susceptibility 1) mutant seedlings^4,51^, germinated and grown on MS agar medium as described above for Col-0, to increase spore inoculum densities.

### Co-inoculation of *gnoHpa* and individual bacterial isolates on axenic plants

For gnotobiotic bioassays, vapor-phase sterilized^53^ Col-0 seeds were sown on agar-solidified Hoagland medium^52^ (pH = 5.5, 0.6% agarose w/v) in 24-well microtiter plates (one seedling per well), stratified at 4 °C for 2 days and subsequently cultivated at 21 °C, 70% relative humidity, 12 h light/12 h dark, light intensity 100 μmol m^-2^ s^-1^.

Bacterial isolates were grown on 1/10 strength TSA plates for 2-3 days, depending on growth speed, at 28 °C. Bacterial suspensions were then prepared by scraping colonies from the TSA agar medium into sterile MgSO_4_ (10 mM), optical density was measured at 600 nm, and diluted to OD600 = 0.2 in MgSO_4_. *GnoHpa* suspensions were prepared by shaking 10-20 sporulating *eds1* mutant plants in 2 mL MgSO_4_ and then transferred to a new 2-mL tube. Mixtures of gno*Hpa/*MgSO_4_, gno*Hpa*/bacteria, and MgSO_4_/bacteria were prepared at 9:1 ratio so that *gnoHpa* spore densities were ~150 spores/μL and bacterial densities were OD600 = 0.02. Leaves of 10-day old Col-0 seedlings were then inoculated with these mixtures. For each seedling, both cotyledons and the first two true leaves were inoculated with a 0.3-μL droplet of suspension. Bacterial densities were quantified 7 days-post inoculation by 10-fold serial dilution plating on 1/10 strength TSA plates and counting colony forming units. *GnoHpa* spore production was quantified 7 days post inoculation as described above. Although all 9 representative HAM bacterial isolates (Supplementary Table S8) were tested, no useful data on bacterial densities were obtained for *Rhizobium* (ASV 2569b) isolate WCS2018Hpa-8 due to contamination of the assay.

### Complementation of *gnoHpa* spores with the Hpa-associated microbiome

*Hpa* spore suspensions were prepared from *Hpa* and *gnoHpa* of isolate Noco2 as describe above. Half of the *Hpa* suspension was filtered using a sterile 10-μm filter, moistened in advance with autoclaved demineralized water, to remove *Hpa* spores and allow the passage of HAM bacteria. The absence of spores in the HAM filtrate was confirmed by microscopy and by spraying the filtrate directly onto susceptible plants, following which no sporulation was observed. Using an equal amount of leaf material from healthy Arabidopsis plants, we also obtained a suspension of phyllosphere resident bacteria. The bacterial HAM and phyllosphere-resident filtrates were subsequently supplemented with equal densities of *gnoHpa* spores.

We then spray inoculated 10 replicate 60-mL pots with 14-day-old Arabidopsis Col-0 plants with 12.5 mL spore suspensions of *Hpa* or of *gnoHpa* mixed with water, HAM filtrate or resident filtrate. Plants were airdried for 2 h, covered with transparent lids to ensure high humidity and incubated (21°C, 10 h light/14 h dark, light intensity 100 μmolm^-2^ s^-1^). Seven days post inoculation, *Hpa* and *gnoHpa* sporulation was quantified as described above.

### Soil-borne legacy experiment

Fourteen-day old Col-0 or transgenic Col-0 *RPP5* plants growing on Reijerscamp soil were inoculated with spores of regular *Hpa* cultures or *gnoHpa* cultures of isolate Noco2 or mock-inoculated as described above. Plants were airdried for 2 h, covered with transparent lids to ensure high humidity and incubated (21°C, 10 h light/14 h dark, light intensity 100 μmol m^-2^ s^-1^). Seven days post inoculation of this conditioning population of plants, shoots were cut off and a new population of Col-0 plants (response population) was sown directly on top of the soil after which the experimental cycle was repeated^54^. For both the conditioning and response population of plants, *Hpa* and *gnoHpa* sporulation was quantified as described above. Phyllosphere, rhizosphere and bulk soil samples were taken at the end of the growth period of the conditioning and response populations, respectively, as described above. An overview of the experimental setup described above is shown in Supplementary Fig. S18.

## Acknowledgements

This study was sponsored by the TopSector Horticulture and Starting Materials (TKI grant no. 1605-106) and by the Dutch Research Council (NWO) through the Gravitation program MiCRop (grant no. 024.004.014), and XL program “Unwiring beneficial functions and regulatory networks in the plant endosphere” (grant no. OCENW.GROOT.2019.063). The TKI project was carried out in collaboration with four industrial partners; DSM, Enza Zaden, Pop Vriend Seeds, and RijkZwaan Breeding B.V. We would like to thank Jane Parker for providing Col-0 *RPP5* and L*er rpp5* Arabidopsis seeds, Ronnie de Jonge for valuable input on bio-informatic approaches, and Joyce Elberse, Deniz Duijker, Lotte Pronk, Cindy Molina Ruiz, Xinya Pan, Lisa Wagenaar and Tilda Tarrant for excellent technical assistance.

