## Supplementary figures and tables for "Congruent downy mildew-associated microbiomes reduce plant disease and function as transferable resistobiomes"

**The following Supplementary information is available for this article:**

<u>Supplementary Figures</u>

Figure S1. **Compatibility of Noco2 and Cala2 with susceptible and resistant Arabidopsis accessions**.

Figure S2. **Phyllosphere fungal community composition is unaffected by *Hpa* inoculation.**

Figure S3. ***Hpa*-associated bacteria dominate the phyllosphere regardless of the plant's susceptibility to and the proliferation of *Hpa*.**

Figure S4. **Genome comparisons between bacterial HAM isolates.**

Figure S5. **Assignment of reads from *Hpa* metagenomes to unique *Acidovorax* genomes**.

Figure S6. **Assignment of reads from Hpa metagenomes to unique *Aeromicrobium* genomes.**

Figure S7. **Assignment of reads from Hpa metagenomes to unique *Arthrobacter* genomes.**

Figure S8. **Assignment of reads from Hpa metagenomes to unique *Sphingobium* genomes.**

Figure S9. **Assignment of reads from Hpa metagenomes to unique *Rhizobium* genomes.**

Figure S10. **Assignment of reads from Hpa metagenomes to unique *Methylobacterium* genomes**

Figure S11. **Assignment of reads from Hpa metagenomes to unique *Microbacterium* genomes.**

Figure S12. **Assignment of reads from Hpa metagenomes to unique *Xanthomonas* genomes.**

Figure S13. **Summary of Hpa metagenome read mapping against HAM isolate genomes and non-redundant genomes of genera**

Figure S14. **Genome comparisons between Xanthomonas a0e1a isolates and a Hpa-metagenome-derived assembly.**

Figure S15. **Schematic overview of the '9-passages experiment', which tests the effect of the removal of *Hpa* on the associated microbiome**.

Figure S16. **The *Hpa*-culture bacterial community is largely unaffected by removal of *Hpa*, but nonetheless there are community shifts in the absence of *Hpa***

Figure S17**. Boxplot of the abundance of *Xanthomonas* ASV a0e1a in 16S rDNA amplicon sequencing data in the 9$^{th}$ plant population from the 9-passages experiment.**

Figure S18. **Setup of soil-borne legacy experiments.**

Figure S19. **Gno*Hpa* inoculated Col-0 plants grown on soils conditioned with *Hpa*- or gno*Hpa* inoculated Col-0 plants can perceive soil-borne legacy.**

Figure S20. ***Hpa* inoculation of a conditioning population of Col-0 plants drastically alters the phyllosphere microbial community composition, whereas the rhizosphere remains largely unaffected**

Figure S21. **HAM ASVs are represented among the top-30 most-strongly enriched ASVs in the phyllosphere, but not the rhizosphere of gno*Hpa*-infected plants.**

Figure S22. ***Aeromicrobium* ASV d93fb is promoted in the phyllosphere by gno*Hpa* infection.**

### Supplementary Tables

Table S1. **Statistical differences (PERMANOVA) between bacterial phyllosphere communities in mock-, Noco2-, and Cala2-treated Arabidopsis plants.**

Table S2. **PERMANOVA of treatment (mock, Noco2, Cala2) and Arabidopsis genotype (C24, Col-0, Ler, Pro-0) on bacterial phyllosphere communities.**

Table S3. **Statistical differences (PERMANOVA) between bacterial phyllosphere communities in mock-, Noco2-, and Cala2-treatment per Arabidopsis genotype.**

Table S4. **Number of genomes included in genome index per genus, as used for metagenome analyses**

Table S5. **Statistical differences (PERMANOVA) in bacterial phyllosphere communities of L*er rpp5* lineages 1-3 (Noco2, Cala2, *Hpa*-free) and untreated L*er rpp5* plants in the 9-passages experiment.**

Table S6. **Statistical differences (PERMANOVA) in bacterial phyllosphere communities of L*er rpp5* lineages 1-3 (Noco2, Cala2, Hpa-free) per passage (1, 5, and 9) in the 9-passages experiment.**

Table S7. **Bacterial ASVs that that are affected by the removal of Hpa from the HAM in the 9-passages experiment.**

Table S8. **Bacterial isolates screened for modulation of *Hpa* reproduction and the effect of *Hpa* on bacterial abundance in a gnotobiotic system.**

Table S9. **ASVs that were significantly enriched in *Hpa*-treated phyllosphere samples in the conditioning phase of the soil-borne legacy experiment.**

Table S10. **Primers used in this study.**

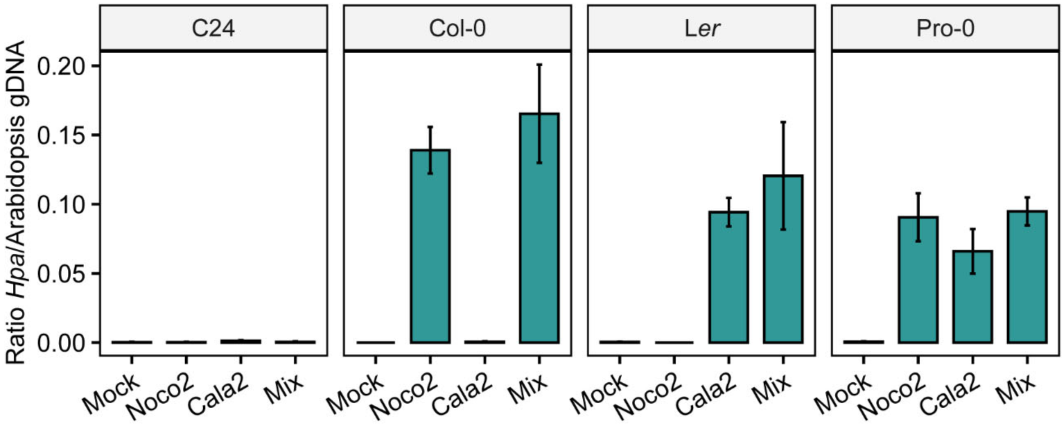


Figure S1. **Compatibility of Noco2 and Cala2 with susceptible and resistant Arabidopsis accessions**. qPCR quantification of *Hpa* abundance in the susceptible and resistant *Hpa*-Arabidopsis interactions indicated, confirming that C24 is resistant to both Noco2 and Cala2, that Col-0 is susceptible to Noco2, that L*er* is susceptible to Cala2, and that Pro-0 is susceptible to both Noco2 and Cala2. qPCR quantification was performed on total genomic DNA from inoculated leaves that were also used for 16S rDNA amplicon sequencing (Fig. 1). *Hpa* abundance was calculated as a ratio of the levels of *ACTIN* in *Hpa* and Arabidopsis. Error bars represent standard error.

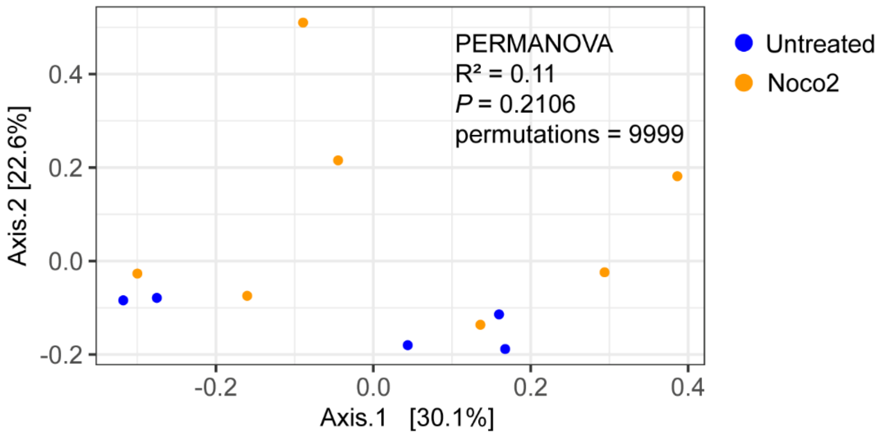


Figure S2. **Phyllosphere fungal community composition is unaffected by *Hpa* inoculation.** PCoA ordination plot of fungal (ITS2 amplicon-based) communities based on Bray-Curtis dissimilarities between untreated (blue symbols), and Noco2-inoculated (orange symbols) Col-0 leaves.

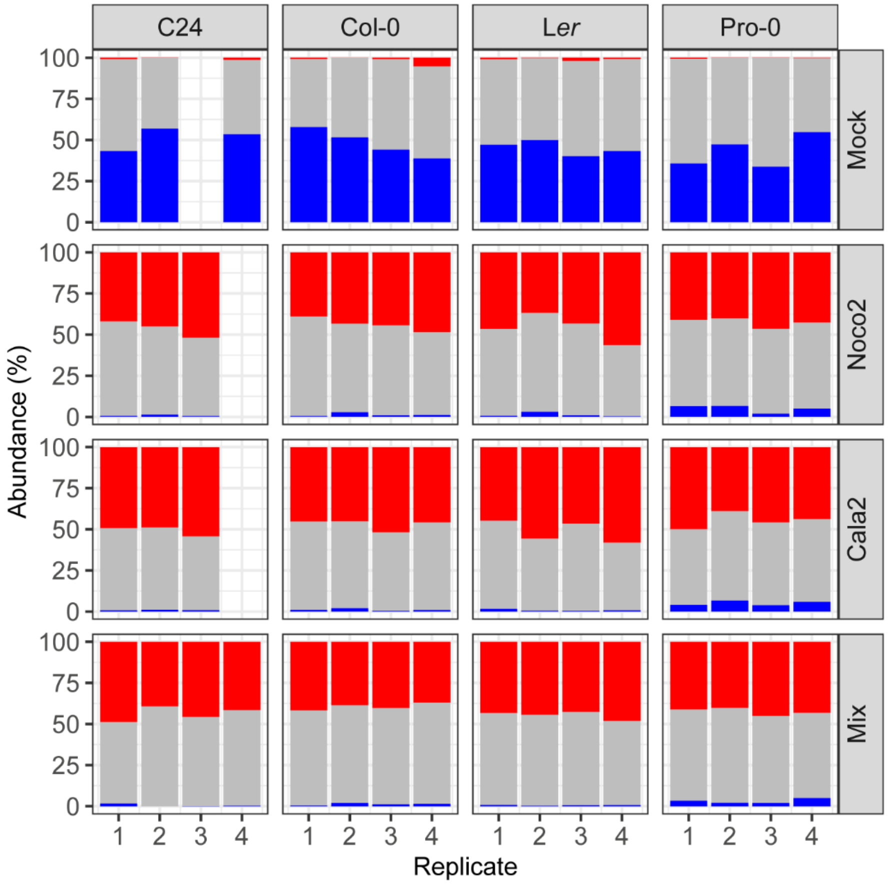


Figure S3. ***Hpa*-associated bacteria dominate the phyllosphere regardless of the plant's susceptibility to and the proliferation of *Hpa*.** Stacked chart with the relative abundance of the 17 ASVs that are enriched in all *Hpa*-inoculated groups (red), the 53 ASVs that are depleted in all *Hpa*-inoculated groups (blue), and all other ASVs in the data (grey) in each inoculation treatment. Each stack represents one individual sample.

| ASV | Genome | Acidovorax_WCS2018Cala2-16 (Utrecht) | Acidovorax_WCS2018Cala2-18 (Utrecht) | Acidovorax_WCS2018Noco2-12 (Utrecht) | Acidovorax_WCS2018Noco2-16 (Utrecht) | **Acidovorax_WCS2018Noco2-17 (Utrecht)** | Acidovorax_WCS2018Noco2-34 (Utrecht) | Acidovorax_WCS2018Noco2-43 (Utrecht) | **Aeromicrobium_WCS2018Hpa-31 (Utrecht** | Aeromicrobium_WCS2018Hpa-33 (Utrecht) | Arthrobacter_WCS2018Hpa-5a (Utrecht) | Arthrobacter_WCS2018Hpa-5b (Utrecht) | **Arthrobacter_WCS2018Hpa-7 (Utrecht)** | **Methylobacterium_WCS2018Hpa-22 (Utre** | **Microbacterium_WCS2018Hpa-23 (Utrecht** | **Microbacterium_WCS2018Hpa-9 (Utrecht)** | **Rhizobium_WCS2018Hpa-16 (Utrecht)** | Rhizobium_WCS2018Hpa-8 (Utrecht) | **Sphingobium_WCS2017Hpa-17 (Cologne)** | Xanthomonas_WCS2017Cala2-12 (Cologne) | Xanthomonas_WCS2017Noco2-62 (Cologne | Xanthomonas_WCS2018Cala2-13 (Utrecht) | Xanthomonas_WCS2018Cala2-18 (Utrecht) | Xanthomonas_WCS2018Cala2-20 (Utrecht) | **Xanthomonas_WCS2018Cala2-21 (Utrecht)** | Xanthomonas_WCS2018Cala2-7 (Utrecht) | Xanthomonas_WCS2018Noco2-14 (Utrecht | Xanthomonas_WCS2018Noco2-15 (Utrecht | Xanthomonas_WCS2018Noco2-18 (Utrecht | Xanthomonas_WCS2018Noco2-27 (Utrecht | Xanthomonas_WCS2018Noco2-28 (Utrecht | Xanthomonas_WCS2019Cala2-53 (Cologne) |
|---|---|---|---|---|---|---|---|---|---|---|---|---|---|---|---|---|---|---|---|---|---|---|---|---|---|---|---|---|---|---|---|---|
| a4065 | Acidovorax_WCS2018Cala2-16 (Utrecht) | 100% | 100% | 100% | 100% | 100% | 100% | 100% | 82% | 82% | 78% | 78% | 78% | 83% | 80% | 81% | 82% | 82% | 82% | 82% | 82% | 82% | 82% | 82% | 82% | 82% | 82% | 82% | 82% | 82% | 82% | 82% |
| a4065 | Acidovorax_WCS2018Cala2-18 (Utrecht) | 100% | 100% | 100% | 100% | 100% | 100% | 100% | 82% | 82% | 78% | 78% | 78% | 83% | 80% | 81% | 82% | 82% | 82% | 82% | 82% | 82% | 82% | 82% | 82% | 82% | 82% | 82% | 82% | 82% | 82% | 82% |
| a4065 | Acidovorax_WCS2018Noco2-12 (Utrecht) | 100% | 100% | 100% | 100% | 100% | 100% | 100% | 82% | 82% | 78% | 78% | 78% | 83% | 80% | 81% | 82% | 82% | 82% | 82% | 82% | 82% | 82% | 82% | 82% | 82% | 82% | 82% | 82% | 82% | 82% | 82% |
| a4065 | Acidovorax_WCS2018Noco2-16 (Utrecht) | 100% | 100% | 100% | 100% | 100% | 100% | 100% | 82% | 82% | 78% | 78% | 78% | 83% | 80% | 81% | 82% | 82% | 82% | 82% | 82% | 82% | 82% | 82% | 82% | 82% | 82% | 82% | 82% | 82% | 82% | 82% |
| **a4065** | **Acidovorax_WCS2018Noco2-17 (Utrecht)** | 100% | 100% | 100% | 100% | 100% | 100% | 100% | 82% | 82% | 78% | 78% | 78% | 83% | 80% | 81% | 82% | 82% | 82% | 82% | 82% | 82% | 82% | 82% | 82% | 82% | 82% | 82% | 82% | 82% | 82% | 82% |
| a4065 | Acidovorax_WCS2018Noco2-34 (Utrecht) | 100% | 100% | 100% | 100% | 100% | 100% | 100% | 82% | 82% | 78% | 78% | 78% | 83% | 80% | 81% | 82% | 82% | 82% | 82% | 82% | 82% | 82% | 82% | 82% | 82% | 82% | 82% | 82% | 82% | 82% | 82% |
| a4065 | Acidovorax_WCS2018Noco2-43 (Utrecht) | 100% | 100% | 100% | 100% | 100% | 100% | 100% | 82% | 82% | 78% | 78% | 78% | 83% | 80% | 81% | 82% | 82% | 82% | 82% | 82% | 82% | 82% | 82% | 82% | 82% | 82% | 82% | 82% | 82% | 82% | 82% |
| **d93fb** | **Aeromicrobium_WCS2018Hpa-31 (Utrecht)** | 82% | 82% | 82% | 82% | 82% | 82% | 82% | 100% | 100% | 83% | 83% | 83% | 82% | 82% | 82% | 82% | 82% | 80% | 80% | 80% | 80% | 80% | 80% | 80% | 80% | 80% | 80% | 80% | 80% | 80% | 80% |
| d93fb | Aeromicrobium_WCS2018Hpa-33 (Utrecht) | 82% | 82% | 82% | 82% | 82% | 82% | 82% | 100% | 100% | 83% | 83% | 83% | 82% | 83% | 82% | 82% | 82% | 80% | 80% | 80% | 80% | 80% | 80% | 80% | 80% | 80% | 80% | 80% | 80% | 80% | 80% |
| 42fbd | Arthrobacter_WCS2018Hpa-5a (Utrecht) | 78% | 78% | 78% | 78% | 78% | 78% | 78% | 83% | 83% | 100% | 100% | 100% | 82% | 84% | 84% | 82% | 82% | 80% | 80% | 80% | 80% | 80% | 80% | 80% | 80% | 80% | 80% | 80% | 80% | 80% | 80% |
| 42fbd | Arthrobacter_WCS2018Hpa-5b (Utrecht) | 78% | 78% | 78% | 78% | 78% | 78% | 78% | 83% | 83% | 100% | 100% | 100% | 82% | 83% | 84% | 82% | 82% | 80% | 80% | 80% | 80% | 80% | 80% | 80% | 80% | 80% | 80% | 80% | 80% | 80% | 80% |
| **42fbd** | **Arthrobacter_WCS2018Hpa-7 (Utrecht)** | 78% | 78% | 78% | 78% | 78% | 78% | 78% | 83% | 83% | 100% | 100% | 100% | 82% | 84% | 84% | 82% | 82% | 80% | 80% | 80% | 80% | 80% | 80% | 80% | 80% | 80% | 80% | 80% | 80% | 80% | 80% |
| **15da8** | **Methylobacterium_WCS2018Hpa-22 (Utrecht)** | 83% | 83% | 83% | 83% | 83% | 83% | 83% | 82% | 82% | 82% | 82% | 82% | 100% | 81% | 81% | 85% | 85% | 82% | 81% | 81% | 81% | 81% | 81% | 81% | 81% | 81% | 81% | 81% | 81% | 81% | 81% |
| **f0c76** | **Microbacterium_WCS2018Hpa-23 (Utrecht)** | 80% | 80% | 80% | 80% | 80% | 80% | 80% | 82% | 83% | 84% | 83% | 84% | 81% | 100% | 89% | 81% | 81% | 80% | 81% | 82% | 81% | 81% | 81% | 81% | 81% | 81% | 81% | 81% | 81% | 81% | 81% |
| **f0c76** | **Microbacterium_WCS2018Hpa-9 (Utrecht)** | 81% | 81% | 81% | 81% | 81% | 81% | 81% | 82% | 82% | 84% | 84% | 84% | 81% | 89% | 100% | 81% | 81% | 80% | 81% | 82% | 81% | 81% | 81% | 81% | 81% | 81% | 81% | 81% | 81% | 81% | 81% |
| **2569b** | **Rhizobium_WCS2018Hpa-16 (Utrecht)** | 82% | 82% | 82% | 82% | 82% | 82% | 82% | 82% | 82% | 82% | 82% | 82% | 85% | 81% | 81% | 100% | 100% | 85% | 86% | 84% | 83% | 83% | 83% | 83% | 83% | 83% | 83% | 83% | 83% | 83% | 86% |
| 2569b | Rhizobium_WCS2018Hpa-8 (Utrecht) | 82% | 82% | 82% | 82% | 82% | 82% | 82% | 82% | 82% | 82% | 82% | 82% | 85% | 81% | 81% | 100% | 100% | 85% | 86% | 84% | 83% | 83% | 83% | 83% | 83% | 83% | 83% | 83% | 83% | 83% | 86% |
| **ed6be** | **Sphingobium_WCS2017Hpa-17 (Cologne)** | 82% | 82% | 82% | 82% | 82% | 82% | 82% | 80% | 80% | 80% | 80% | 80% | 82% | 80% | 80% | 85% | 85% | 100% | 81% | 81% | 81% | 81% | 81% | 81% | 81% | 81% | 81% | 81% | 81% | 81% | 81% |
| a0e1a | Xanthomonas_WCS2017Cala2-12 (Cologne) | 82% | 82% | 82% | 82% | 82% | 82% | 82% | 80% | 80% | 80% | 80% | 80% | 81% | 81% | 81% | 86% | 86% | 81% | 100% | 100% | 100% | 100% | 100% | 100% | 100% | 100% | 100% | 100% | 100% | 100% | 100% |
| a0e1a | Xanthomonas_WCS2017Noco2-62 (Cologne) | 82% | 82% | 82% | 82% | 82% | 82% | 82% | 80% | 80% | 80% | 80% | 80% | 81% | 82% | 82% | 84% | 84% | 81% | 100% | 100% | 100% | 100% | 100% | 100% | 100% | 100% | 100% | 100% | 100% | 100% | 100% |
| a0e1a | Xanthomonas_WCS2018Cala2-13 (Utrecht) | 82% | 82% | 82% | 82% | 82% | 82% | 82% | 80% | 80% | 80% | 80% | 80% | 81% | 81% | 81% | 83% | 83% | 81% | 100% | 100% | 100% | 100% | 100% | 100% | 100% | 100% | 100% | 100% | 100% | 100% | 100% |
| a0e1a | Xanthomonas_WCS2018Cala2-18 (Utrecht) | 82% | 82% | 82% | 82% | 82% | 82% | 82% | 80% | 80% | 80% | 80% | 80% | 81% | 81% | 81% | 83% | 83% | 81% | 100% | 100% | 100% | 100% | 100% | 100% | 100% | 100% | 100% | 100% | 100% | 100% | 100% |
| a0e1a | Xanthomonas_WCS2018Cala2-20 (Utrecht) | 82% | 82% | 82% | 82% | 82% | 82% | 82% | 80% | 80% | 80% | 80% | 80% | 81% | 81% | 81% | 83% | 83% | 81% | 100% | 100% | 100% | 100% | 100% | 100% | 100% | 100% | 100% | 100% | 100% | 100% | 100% |
| **a0e1a** | **Xanthomonas_WCS2018Cala2-21 (Utrecht)** | 82% | 82% | 82% | 82% | 82% | 82% | 82% | 80% | 80% | 80% | 80% | 80% | 81% | 81% | 81% | 83% | 83% | 81% | 100% | 100% | 100% | 100% | 100% | 100% | 100% | 100% | 100% | 100% | 100% | 100% | 100% |
| a0e1a | Xanthomonas_WCS2018Cala2-7 (Utrecht) | 82% | 82% | 82% | 82% | 82% | 82% | 82% | 80% | 80% | 80% | 80% | 80% | 81% | 81% | 81% | 83% | 83% | 81% | 100% | 100% | 100% | 100% | 100% | 100% | 100% | 100% | 100% | 100% | 100% | 100% | 100% |
| a0e1a | Xanthomonas_WCS2018Noco2-14 (Utrecht) | 82% | 82% | 82% | 82% | 82% | 82% | 82% | 80% | 80% | 80% | 80% | 80% | 81% | 81% | 81% | 83% | 83% | 81% | 100% | 100% | 100% | 100% | 100% | 100% | 100% | 100% | 100% | 100% | 100% | 100% | 100% |
| a0e1a | Xanthomonas_WCS2018Noco2-15 (Utrecht) | 82% | 82% | 82% | 82% | 82% | 82% | 82% | 80% | 80% | 80% | 80% | 80% | 81% | 81% | 81% | 83% | 83% | 81% | 100% | 100% | 100% | 100% | 100% | 100% | 100% | 100% | 100% | 100% | 100% | 100% | 100% |
| a0e1a | Xanthomonas_WCS2018Noco2-18 (Utrecht) | 82% | 82% | 82% | 82% | 82% | 82% | 82% | 80% | 80% | 80% | 80% | 80% | 81% | 81% | 81% | 83% | 83% | 81% | 100% | 100% | 100% | 100% | 100% | 100% | 100% | 100% | 100% | 100% | 100% | 100% | 100% |
| a0e1a | Xanthomonas_WCS2018Noco2-27 (Utrecht) | 82% | 82% | 82% | 82% | 82% | 82% | 82% | 80% | 80% | 80% | 80% | 80% | 81% | 81% | 81% | 83% | 83% | 81% | 100% | 100% | 100% | 100% | 100% | 100% | 100% | 100% | 100% | 100% | 100% | 100% | 100% |
| a0e1a | Xanthomonas_WCS2018Noco2-28 (Utrecht) | 82% | 82% | 82% | 82% | 82% | 82% | 82% | 80% | 80% | 80% | 80% | 80% | 81% | 81% | 81% | 83% | 83% | 81% | 100% | 100% | 100% | 100% | 100% | 100% | 100% | 100% | 100% | 100% | 100% | 100% | 100% |
| a0e1a | Xanthomonas_WCS2019Cala2-53 (Cologne) | 82% | 82% | 82% | 82% | 82% | 82% | 82% | 80% | 80% | 80% | 80% | 80% | 81% | 81% | 81% | 86% | 86% | 81% | 100% | 100% | 100% | 100% | 100% | 100% | 100% | 100% | 100% | 100% | 100% | 100% | 100% |

Figure S4. **Genome comparisons between bacterial HAM isolates.** Heatmap with average nucleotide identities (%) between genomes of 31 bacterial isolates matching with 8 HAM bacterial ASVs, including the 4 *Hpa*-core ASVs. Isolates matching the same ASV that allowed for comparisons between Noco2- and Cala2 culture-derived isolates are indicated in orange and green, respectively. Genomes that were used as representatives for further analyses (Fig. S5 – S12) are highlighted in bold text.

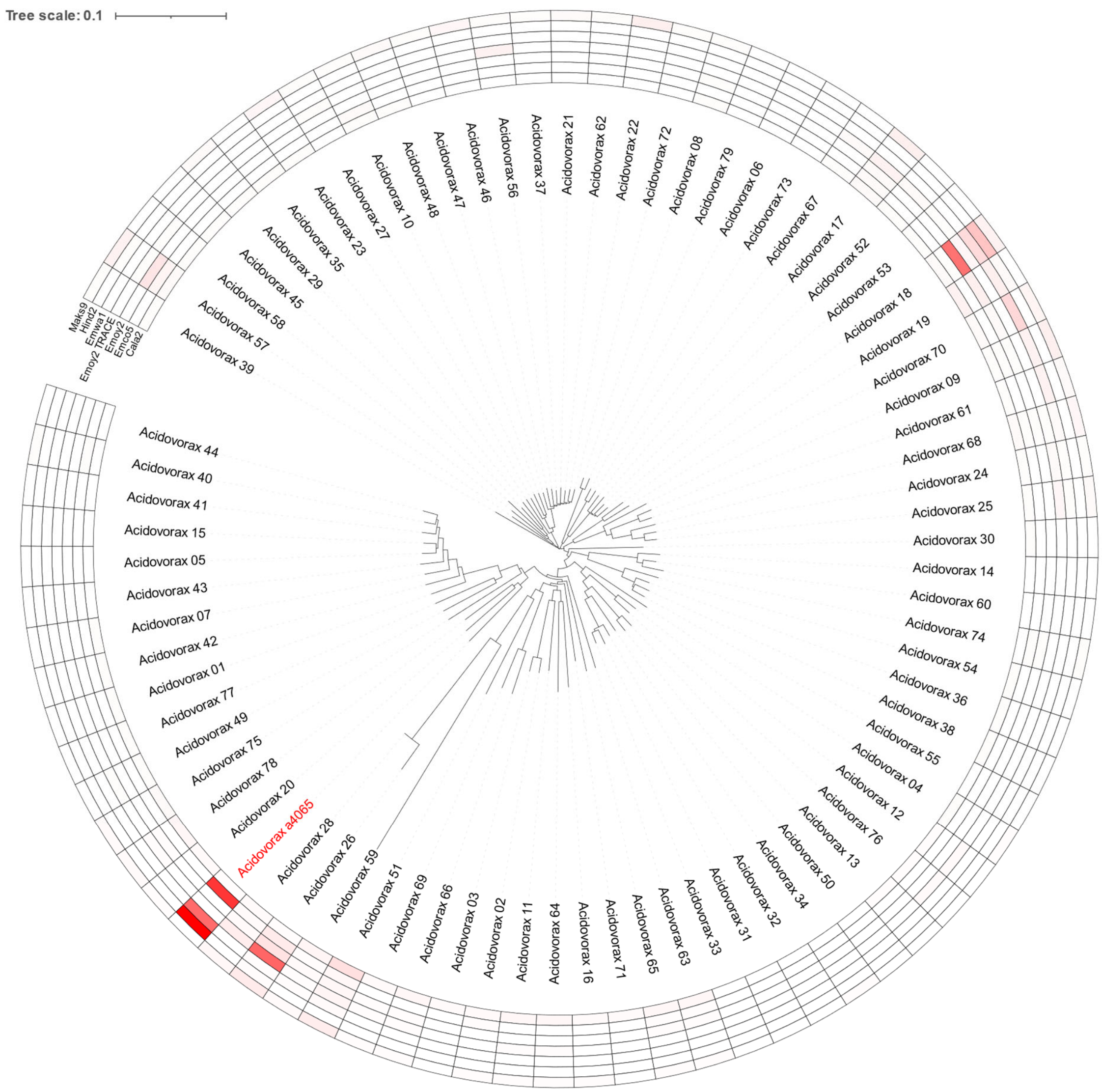


Figure S5 **Assignment of reads from *Hpa* metagenomes to unique *Acidovorax* genomes.** Mashtree dendrogram of non-redundant *Acidovorax* genomes and a heatmap of read-assignment from *Hpa* metagenomes to the bacterial genomes. The genome of *the* isolate representing *Acidovorax* HAM ASV a4065 is designated with a red label. Each row in the heatmap represents one of the seven *Hpa* metagenomes. Cells are colored based on the percentage of reads within that metagenome that pseudo-aligned to that specific genome (dendrogram leaf), as a proportion of the total number of reads that were pseudo-aligned to all genomes present in the dendrogram. Colors are a gradient from 0% (white) to the maximum observed percentage within a metagenome (70%, red).

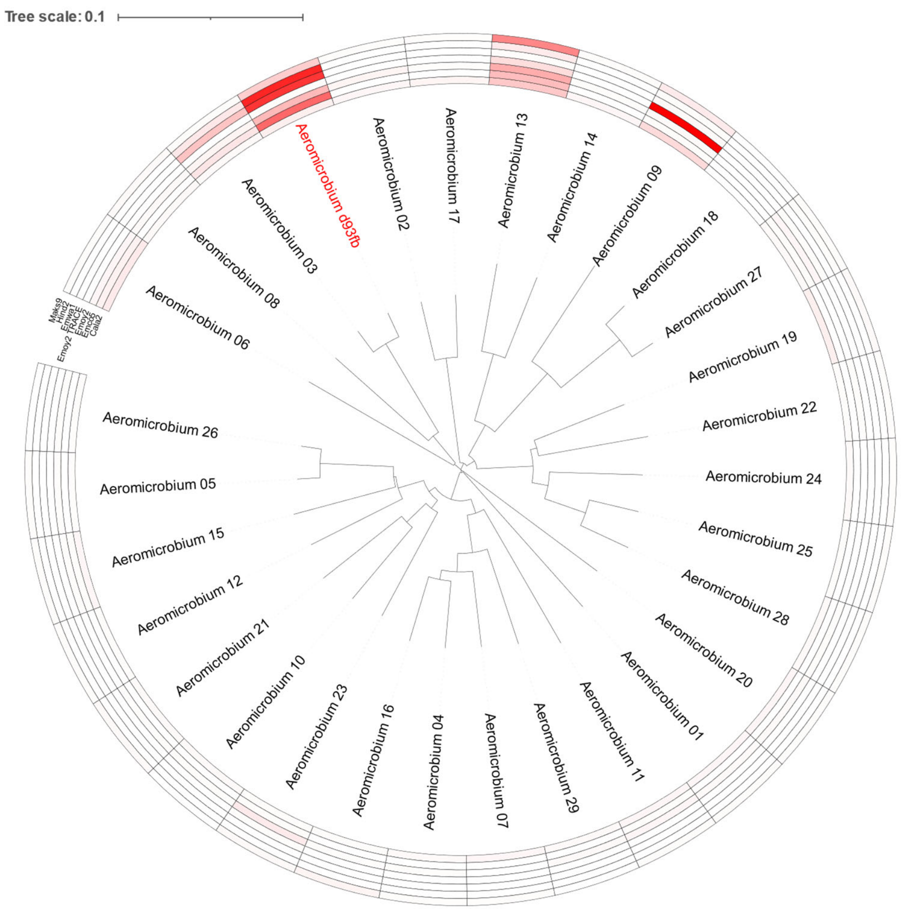


Figure S6 **Assignment of reads from *Hpa* metagenomes to unique *Aeromicrobium* genomes.** Mashtree dendrogram of non-redundant *Aeromicrobium* genomes and a heatmap of read-assignment from *Hpa* metagenomes to the bacterial genomes The genome of *the* isolate representing *Aeromicrobium* HAM ASV d93fb is designated with a red label. Each row in the heatmap represents one of the seven *Hpa* metagenomes. Cells are colored based on the percentage of reads within that metagenome that pseudo-aligned to that specific genome (dendrogram leaf), as a proportion of the total number of reads that were pseudo-aligned to all genomes present in the dendrogram. Colors are a gradient from 0% (white) to the maximum observed percentage within a metagenome (83%, red).

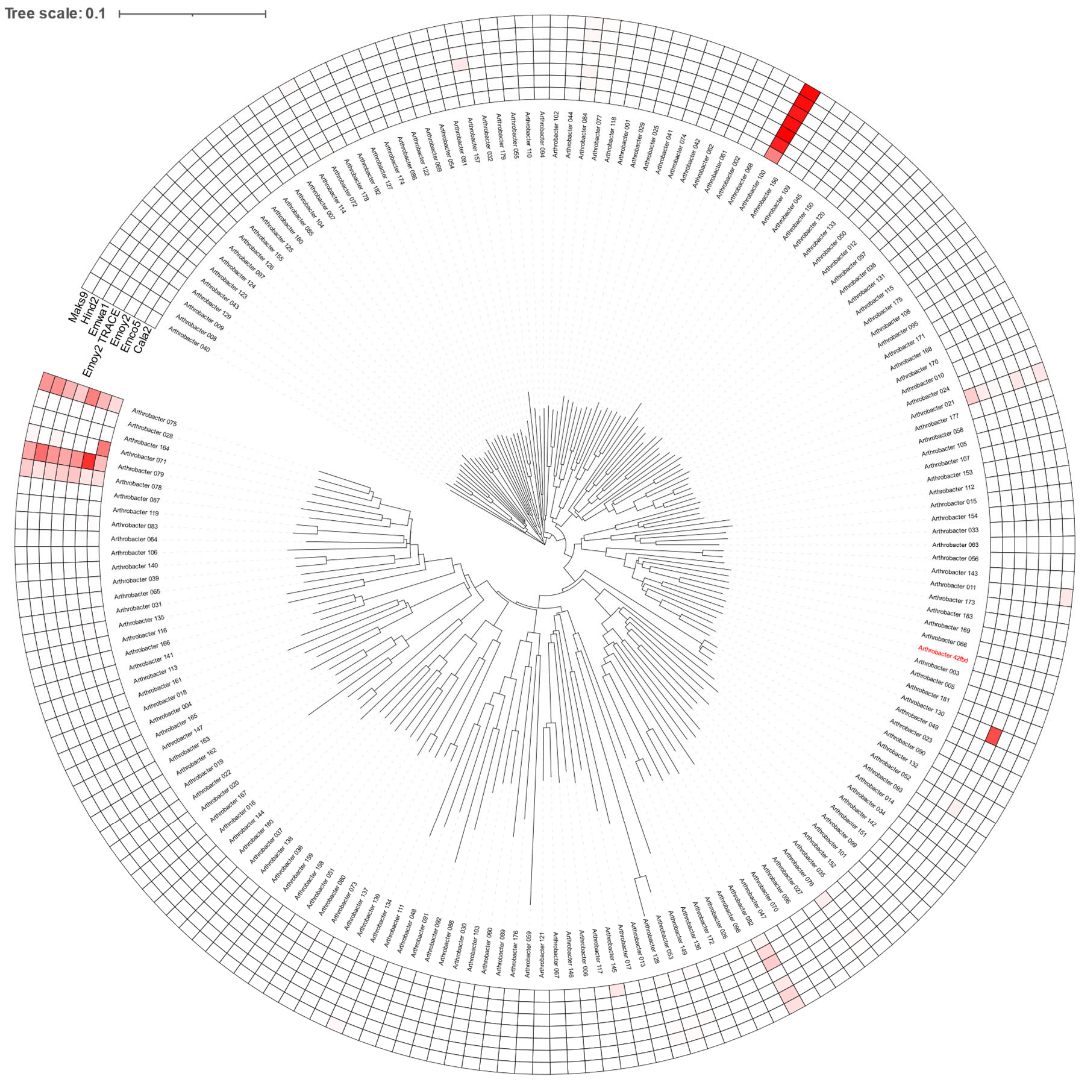


Figure S7 **Assignment of reads from *Hpa* metagenomes to unique *Arthrobacter* genomes.** Mashtree dendrogram of non-redundant *Arthrobacter* genomes and a heatmap of read-assignment from *Hpa* metagenomes to the bacterial genomes. The genome of *the* isolate representing *Arthrobacter* HAM ASV 42fbd is designated with a red label. Each row in the heatmap represents one of the seven *Hpa* metagenomes. Cells are colored based on the percentage of reads within that metagenome that pseudo-aligned to that specific genome (dendrogram leaf), as a proportion of the total number of reads that were pseudo-aligned to all genomes present in the dendrogram. Colors are a gradient from 0% (white) to the maximum observed percentage within a metagenome (40%, red).

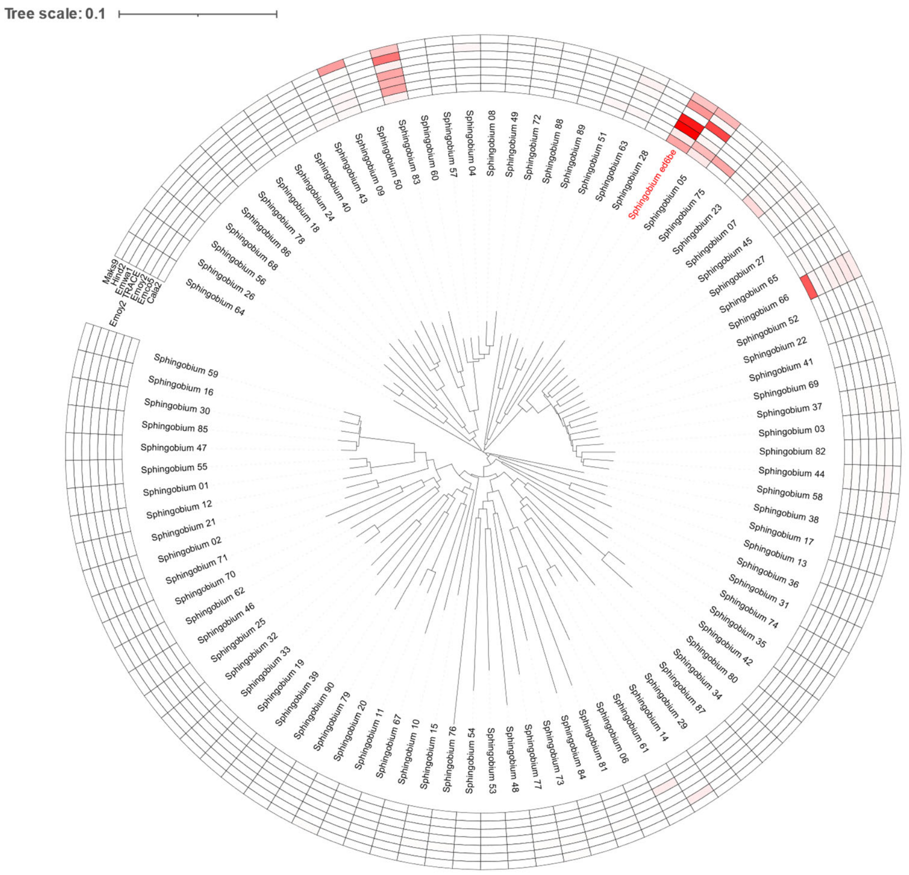


Figure S8 **Assignment of reads from *Hpa* metagenomes to unique *Sphingobium* genomes.** Mashtree dendrogram of non-redundant *Sphingobium* genomes and a heatmap of read-assignment from *Hpa* metagenomes to the bacterial genomes The genome of *the* isolate representing *Sphingobium* HAM ASV ed6be is designated with a red label. Each row in the heatmap represents one of the seven *Hpa* metagenomes. Cells are colored based on the percentage of reads within that metagenome that pseudo-aligned to that specific genome (dendrogram leaf), as a proportion of the total number of reads that were pseudo-aligned to all genomes present in the dendrogram. Colors are a gradient from 0% (white) to the maximum observed percentage within a metagenome (64%, red).

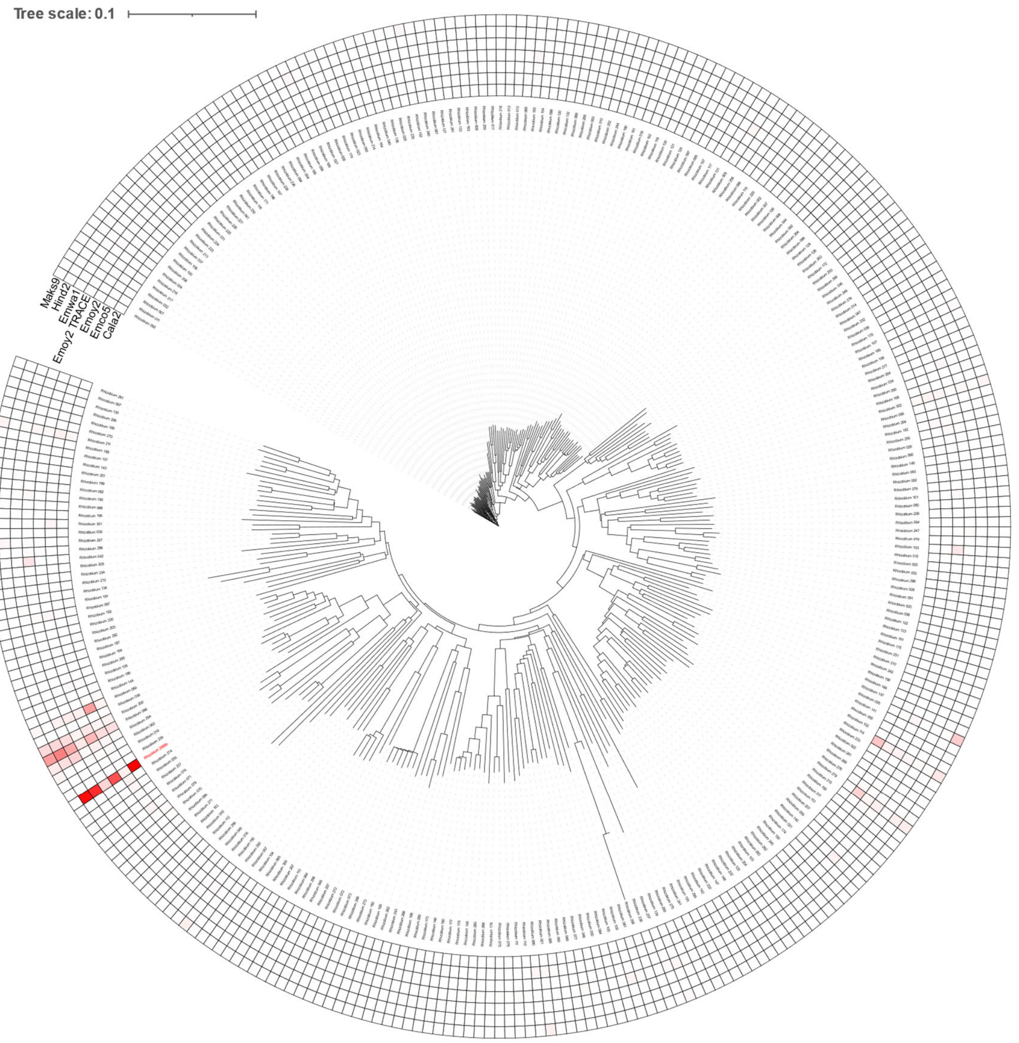


Figure S9 **Assignment of reads from *Hpa* metagenomes to unique *Rhizobium* genomes.** Mashtree dendrogram of non-redundant *Rhizobium* genomes and a heatmap of read-assignment from *Hpa* metagenomes to the bacterial genomes The genome of *the* isolate representing *Rhizobium* HAM ASV 2569b is designated with a red label. Each row in the heatmap represents one of the seven *Hpa* metagenomes. Cells are colored based on the percentage of reads within that metagenome that pseudo-aligned to that specific genome (dendrogram leaf), as a proportion of the total number of reads that were pseudo-aligned to all genomes present in the dendrogram. Colors are a gradient from 0% (white) to the maximum observed percentage within a metagenome (50%, red).

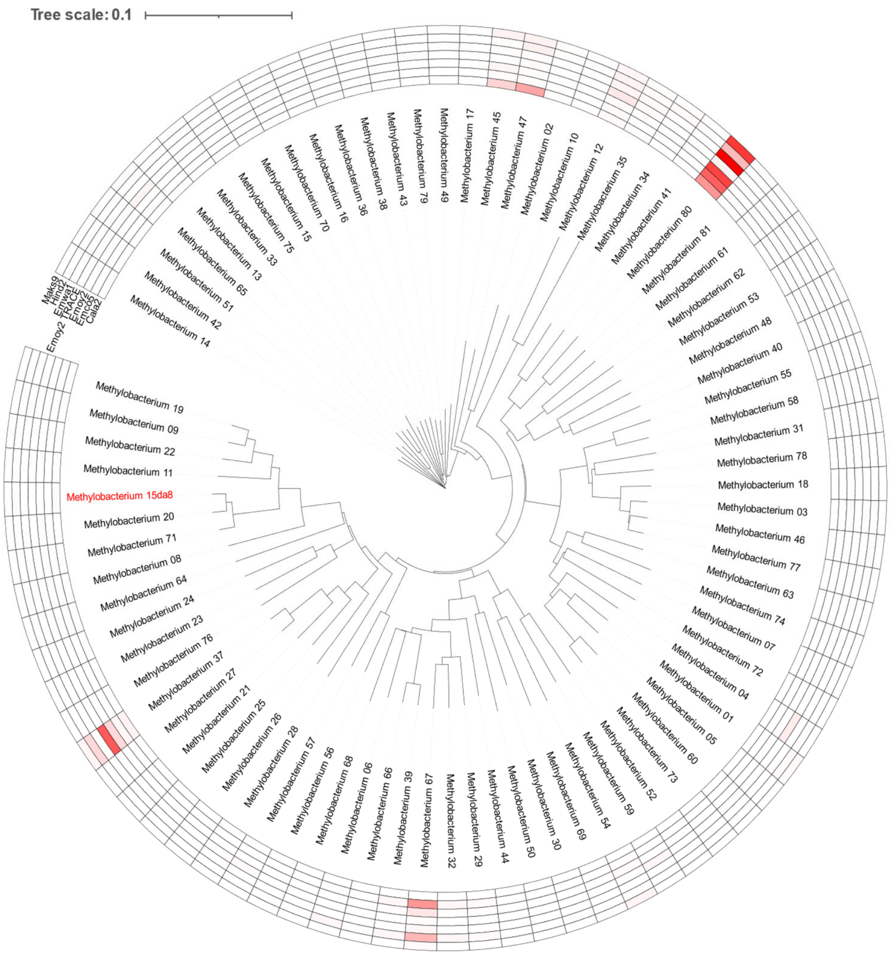


Figure S10 **Assignment of reads from *Hpa* metagenomes to unique *Methylobacterium* genomes.** Mashtree dendrogram of non-redundant *Methylobacterium* genomes and a heatmap of read-assignment from *Hpa* metagenomes to the bacterial genomes The genome of *the* isolate representing *Methylobacterium* HAM ASV 2569b is designated with a red label. Each row in the heatmap represents one of the seven *Hpa* metagenomes. Cells are colored based on the percentage of reads within that metagenome that pseudo-aligned to that specific genome (dendrogram leaf), as a proportion of the total number of reads that were pseudo-aligned to all genomes present in the dendrogram. Colors are a gradient from 0% (white) to the maximum observed percentage within a metagenome (69%, red).

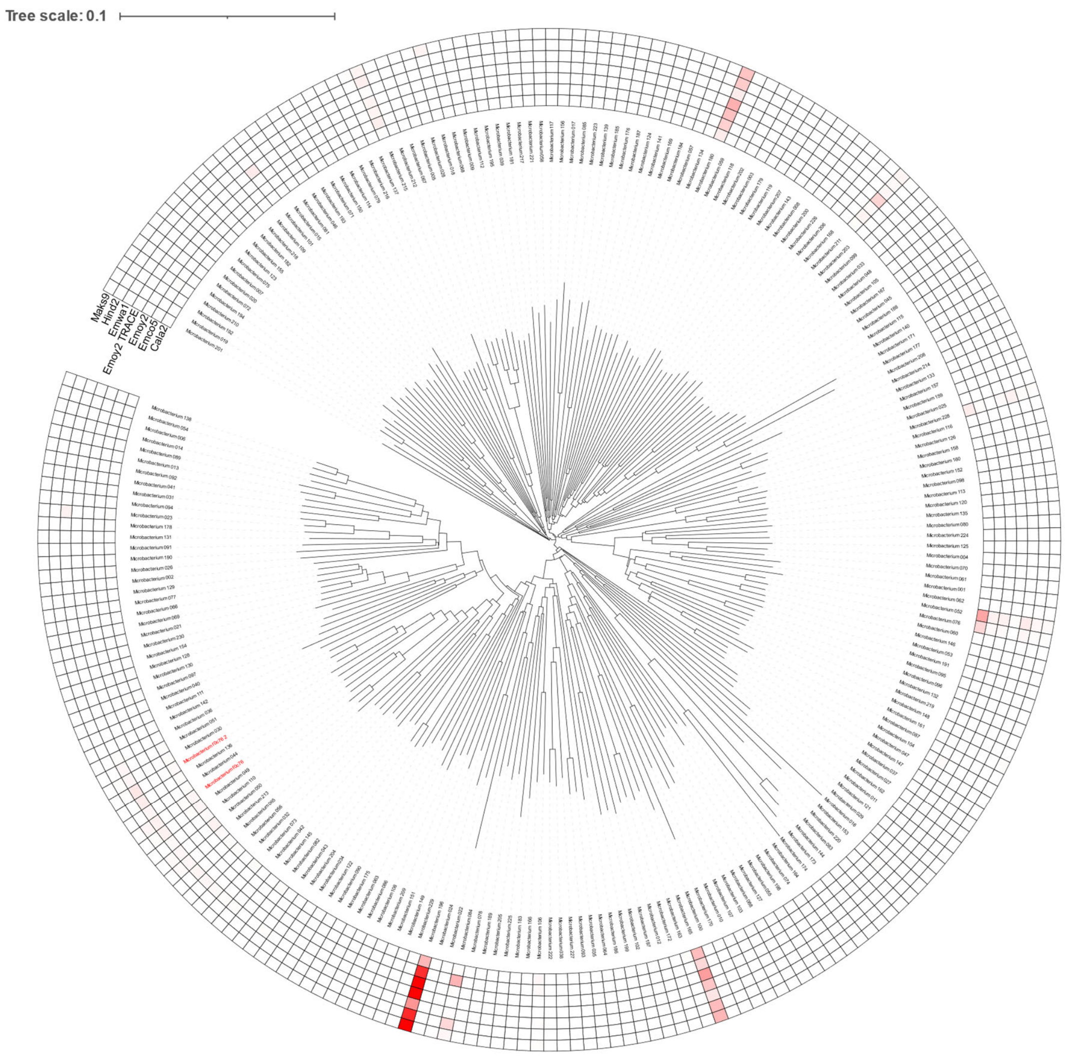


Figure S11 **Assignment of reads from *Hpa* metagenomes to unique *Microbacterium* genomes.** Mashtree dendrogram of non-redundant *Microbacterium* genomes and a heatmap of read-assignment from *Hpa* metagenomes to the bacterial genomes The genome of 2 isolates representing *Microbacterium* HAM ASV 2569b are designated with a red label. Each row in the heatmap represents one of the seven *Hpa* metagenomes. Cells are colored based on the percentage of reads within that metagenome that pseudo-aligned to that specific genome (dendrogram leaf), as a proportion of the total number of reads that were pseudo-aligned to all genomes present in the dendrogram. Colors are a gradient from 0% (white) to the maximum observed percentage within a metagenome (52%, red).

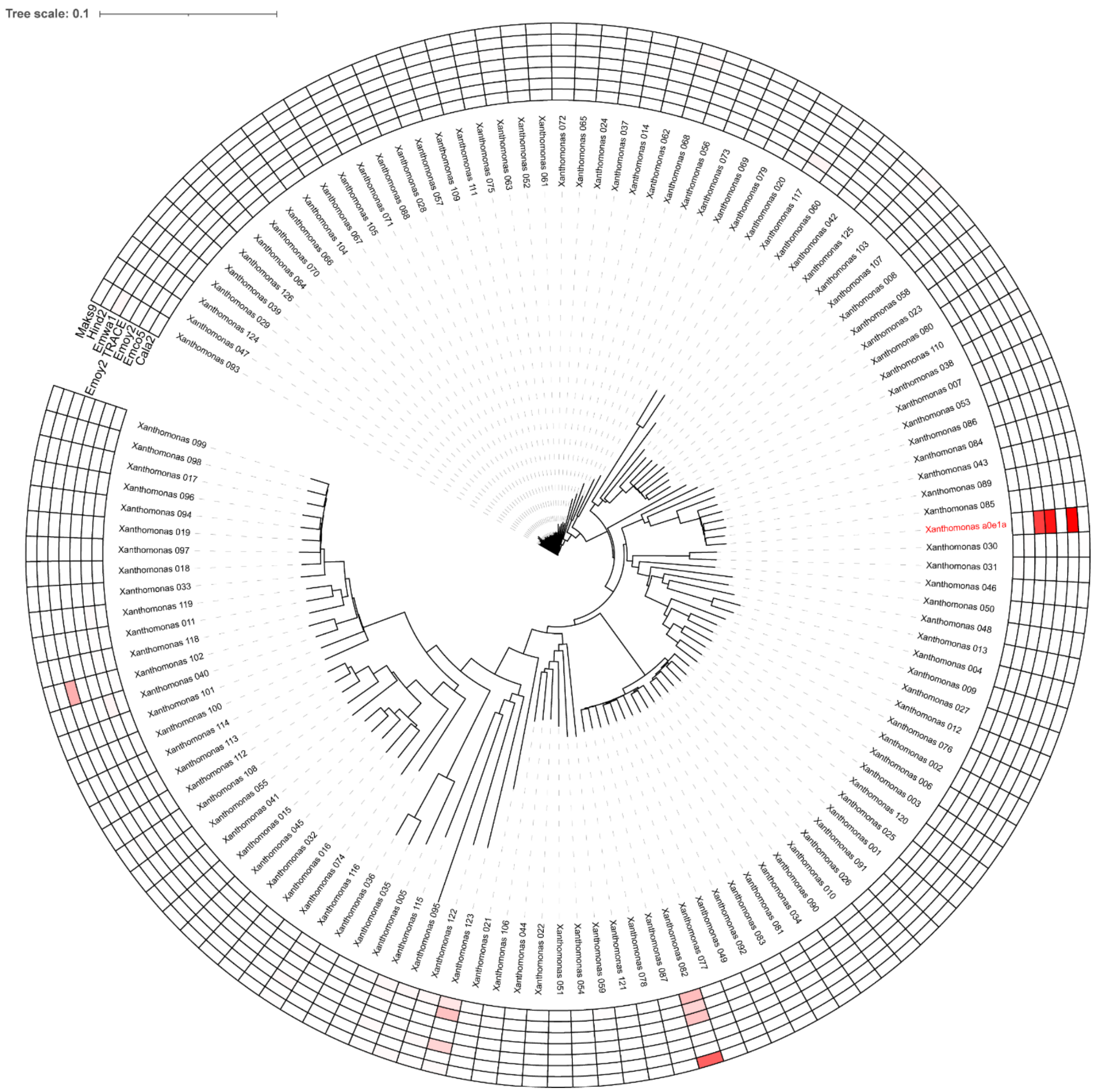


Figure S12 **Assignment of reads from *Hpa* metagenomes to unique *Xanthomonas* genomes.** Mashtree dendrogram of non-redundant *Xanthomonas* genomes and a heatmap of read-assignment from *Hpa* metagenomes to the bacterial genomes The genome of  the isolate representing *Xanthomonas* HAM ASV a0e1a are designated with a red label. Each row in the heatmap represents one of the seven *Hpa* metagenomes. Cells are colored based on the percentage of reads within that metagenome that pseudo-aligned to that specific genome (dendrogram leaf), as a proportion of the total number of reads that were pseudo-aligned to all genomes present in the dendrogram. Colors are a gradient from 0% (white) to the maximum observed percentage within a metagenome (97%, red).

| | HAM genome | genus | HAM genome | genus | HAM genome | genus | HAM genome | genus | HAM genome | genus | HAM genome | genus | HAM genome | genus |
|---|---|---|---|---|---|---|---|---|---|---|---|---|---|---|
| *Xanthomonas* a0e1a | 128 | 32314 | 267 | 19876 | 44635 | 60910 | 4180 | 4723 | 786 | 110392 | 193314 | 198928 | 148 | 15507 |
| *Aeromicrobium* d93fb | 93 | 2019 | 3069 | 4698 | 122 | 521 | 53 | 3948 | 8094 | 12194 | 1671 | 2342 | 148 | 1129 |
| *Sphingobium* ed6be | 18084 | 77564 | 3038 | 40898 | 10356 | 16137 | 1597 | 2738 | 424 | 70179 | 3387 | 12744 | 2579 | 17975 |
| *Rhizobium* 2569b | 87445 | 179698 | 1958 | 65448 | 5357 | 15646 | 164 | 2016 | 141949 | 342440 | 23837 | 49505 | 114 | 29250 |
| *Acidovorax* a4065 | 819 | 55882 | 33045 | 54600 | 46 | 12949 | 12 | 1857 | 150684 | 368391 | 26687 | 37850 | 137 | 19614 |
| *Microbacterium* f0c76 1 | 1072 | 62399 | 1006 | 97644 | 346 | 79077 | 13 | 4044 | 3519 | 125111 | 865 | 43926 | 82 | 42232 |
| *Microbacterium* f0c76 2 | 1009 | 62399 | 609 | 97644 | 241 | 79077 | 1 | 4044 | 6117 | 125111 | 419 | 43926 | 114 | 42232 |
| *Methylobacterium* 15da8 | 188 | 30350 | 35 | 20233 | 13 | 3313 | 0 | 996 | 213 | 50693 | 24 | 2806 | 14 | 12640 |
| *Arthrobacter* 42fbd | 36 | 57196 | 83 | 136175 | 7 | 108681 | 17 | 12786 | 48 | 93114 | 12 | 51484 | 28 | 73728 |
| | Cala2 | | Emco5 | | Emoy2 | | Emoy2_BAC | | Emwa1 | | Hind2 | | Maks9 | |

Figure S13. **Summary of *Hpa* metagenome read mapping against HAM isolate genomes and non-redundant genomes of genera**. For each *Hpa* metagenome (indicated in bottom row) the numbers indicate how many reads were mapped on the genome of interest (left) and on the all genomes withing the corresponding genomes (right, this number includes the number of reads mapped on the corresponding HAM genome).

| | Emoy2_TRACE_assembly | WCS2017Cala2-12 | WCS2019Cala2-53 | WCS2017Noco2-62 | WCS2018Noco2-28 | WCS2018Noco2-18 | WCS2018Cala2-21 | WCS2018Cala2-20 | WCS2018Cala2-7 | WCS2018Cala2-13 | WCS2018Noco2-14 | WCS2018Noco2-15 | WCS2018Noco2-27 | WCS2018Cala2-18 | X. WCS2014-23 |
|---|---|---|---|---|---|---|---|---|---|---|---|---|---|---|---|
| Emoy2_BAC_assembly | 100.00% | 99.96% | 99.97% | 99.96% | 99.97% | 99.98% | 99.97% | 99.97% | 99.97% | 99.96% | 99.98% | 99.97% | 99.99% | 99.94% | 99.99% |
| WCS2017Cala2-12 (Cologne) | 99.96% | 100.00% | 100.00% | 100.00% | 100.00% | 99.99% | 100.00% | 100.00% | 99.99% | 100.00% | 100.00% | 100.00% | 100.00% | 99.99% | 99.99% |
| WCS2019Cala2-53 (Cologne) | 99.97% | 100.00% | 100.00% | 100.00% | 100.00% | 99.99% | 100.00% | 100.00% | 100.00% | 100.00% | 100.00% | 100.00% | 100.00% | 99.99% | 99.99% |
| WCS2017Noco2-62 (Cologne) | 99.96% | 100.00% | 100.00% | 100.00% | 100.00% | 99.99% | 100.00% | 100.00% | 100.00% | 100.00% | 100.00% | 100.00% | 100.00% | 99.99% | 99.99% |
| WCS2018Noco2-28 (Utrecht) | 99.97% | 100.00% | 100.00% | 100.00% | 100.00% | 99.99% | 100.00% | 100.00% | 100.00% | 99.99% | 100.00% | 100.00% | 100.00% | 99.99% | 99.99% |
| WCS2018Noco2-18 (Utrecht) | 99.98% | 99.99% | 99.99% | 99.99% | 99.99% | 100.00% | 99.99% | 99.99% | 99.99% | 99.99% | 99.99% | 99.99% | 99.99% | 99.99% | 99.99% |
| WCS2018Cala2-21 (Utrecht) | 99.97% | 100.00% | 100.00% | 100.00% | 100.00% | 99.99% | 100.00% | 100.00% | 100.00% | 99.99% | 100.00% | 100.00% | 100.00% | 99.99% | 99.99% |
| WCS2018Cala2-20 (Utrecht) | 99.97% | 100.00% | 100.00% | 100.00% | 100.00% | 99.99% | 100.00% | 100.00% | 100.00% | 99.99% | 100.00% | 100.00% | 100.00% | 100.00% | 99.99% |
| WCS2018Cala2-7 (Utrecht) | 99.97% | 99.99% | 100.00% | 100.00% | 100.00% | 99.99% | 100.00% | 100.00% | 100.00% | 100.00% | 100.00% | 100.00% | 100.00% | 100.00% | 99.99% |
| WCS2018Cala2-13 (Utrecht) | 99.96% | 100.00% | 100.00% | 100.00% | 99.99% | 99.99% | 99.99% | 99.99% | 100.00% | 100.00% | 100.00% | 99.99% | 100.00% | 100.00% | 99.99% |
| WCS2018Noco2-14 (Utrecht) | 99.98% | 100.00% | 100.00% | 100.00% | 100.00% | 99.99% | 100.00% | 100.00% | 100.00% | 100.00% | 100.00% | 100.00% | 100.00% | 99.99% | 99.99% |
| WCS2018Noco2-15 (Utrecht) | 99.97% | 100.00% | 100.00% | 100.00% | 100.00% | 99.99% | 100.00% | 100.00% | 100.00% | 99.99% | 100.00% | 100.00% | 100.00% | 99.99% | 99.99% |
| WCS2018Noco2-27 (Utrecht) | 99.99% | 100.00% | 100.00% | 100.00% | 100.00% | 99.99% | 100.00% | 100.00% | 100.00% | 100.00% | 100.00% | 100.00% | 100.00% | 100.00% | 99.99% |
| WCS2018Cala2-18 (Utrecht) | 99.94% | 99.99% | 99.99% | 99.99% | 99.99% | 99.99% | 99.99% | 100.00% | 100.00% | 100.00% | 99.99% | 99.99% | 100.00% | 100.00% | 99.99% |
| X. WCS2014-23 (Utrecht) | 99.99% | 99.99% | 99.99% | 99.99% | 99.99% | 99.99% | 99.99% | 99.99% | 99.99% | 99.99% | 99.99% | 99.99% | 99.99% | 99.99% | 100.00% |

Figure S14. **Genome comparisons between *Xanthomonas* a0e1a isolates and a *Hpa*-metagenome-derived assembly.** Heatmap with average nucleotide identities (%) between genomes of HAM *Xanthomonas* bacteria, *Xanthomonas* sp. WCS2014-23, and an assembly based on reads from Sanger sequencing of selected BAC clones derived from *Hpa* isolate Emoy2[29].

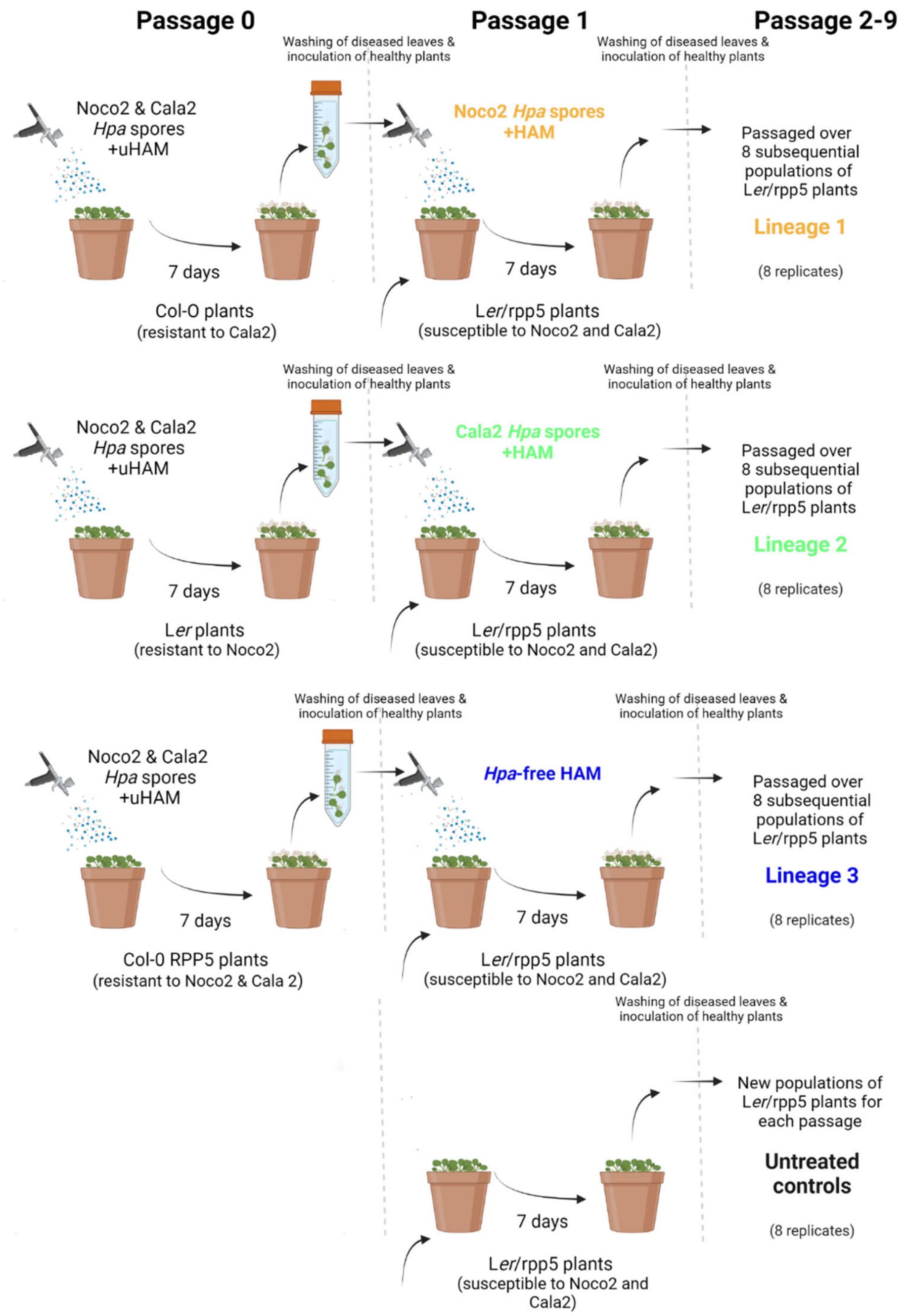


Figure S15. **Schematic overview of the '9-passages experiment', which tests the effect of the removal of *Hpa* on its associated microbiome**. A uniform HAM (uHAM) containing a mix of of Noco2 and Cala2 spore suspensions was spray-inoculated on different Arabidopsis genotypes to selectively remove Noco2 and/or Cala2 from the microbiome (HAM) that travels together with these *Hpa* isolates upon passaging to new host plants. One week post-inoculation, Col-0 (Lineage 1) and L*er* plants (Lineage 2) sporulated with Noco2 and Cala2, respectively, and Col-0/*RPP5* transgenic plants (Lineage 3) did not sporulate. From each Arabidopsis genotype, a leaf wash-off was obtained, containing Noco2, Cala2, or no *Hpa*, and sprayed on eight pots containing small fields of L*er*/*rpp5* mutant plants, which are susceptible to both Noco2 and Cala2. All pots were then placed in individual plastic containers, to prevent cross-contamination between pots. One week post-inoculation, the L*er*/*rpp5* plants that were inoculated with Noco2 (Lineage 1) or Cala2 (Lineage 2), sporulated, while the L*er*/*rpp5* plants that were inoculated with the leaf wash-off without *Hpa* spores (Lineage 3) did not display disease symptoms. From each individual pot, the leaf wash-off was sprayed on a new pot containing L*er*/*rpp5* plants, thereby propagating eight separate phyllosphere microbiomes or *Hpa* cultures per lineage. This process was maintained for nine consecutive weeks, allowing eight separate lineages of Noco2, Cala2, or the uHAM without *Hpa* to develop independently. Eight untreated control pots with L*er*/*rpp5* plants were included for all planting cycles.

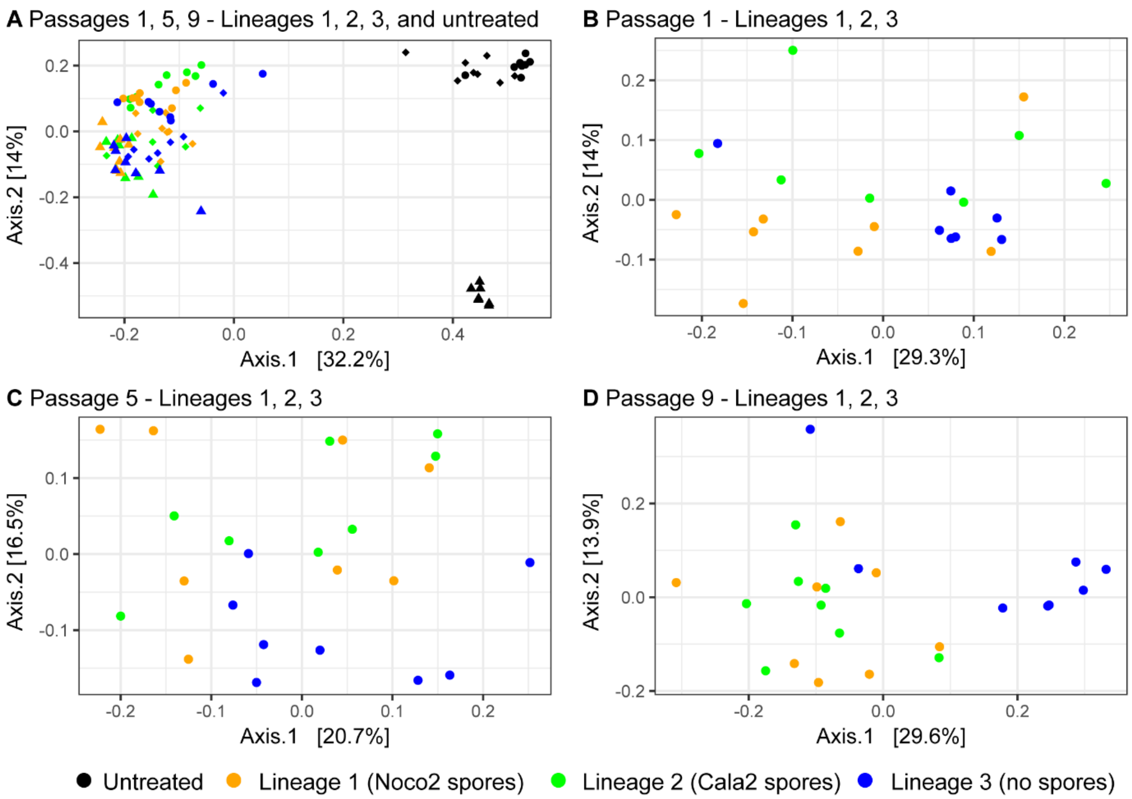


Figure S16. **The *Hpa*-culture bacterial community is largely unaffected by removal of *Hpa*, but nonetheless there are community shifts in the absence of *Hpa*.** PcoA plots based on Bray-Curtis dissimilarities of **A)** all samples from Lineages 1-3 and untreated plants of passages 1 (circles), passage 5 (triangles) and passage 9 (diamonds); and of all inoculated samples of Lineage 1-3 from **B)** passage 1, **C)** passage 5, and **D)** passage 9. Plants were left untreated (black symbols) or were inoculated with leaf wash-offs from Lineage 1 containing Noco2 (orange symbols), from Lineage 2 containing Cala2 (green symbols), or from Lineage 3 which remained *Hpa* free (blue symbols).

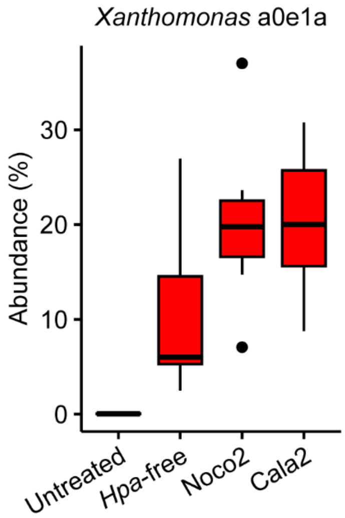


Figure S17**. Boxplot of the abundance of *Xanthomonas* ASV a0e1a in 16S rDNA amplicon sequencing data in the 9$^{th}$ plant population from the 9-passages experiment.** Plants were either untreated or inoculated with a leaf wash suspension from the 9$^{th}$ passage of lineage 1 (containing Noco2 spores), lineage 2 (containing Cala2 spores), or lineage 3 (containing no spores; '*Hpa*-free').

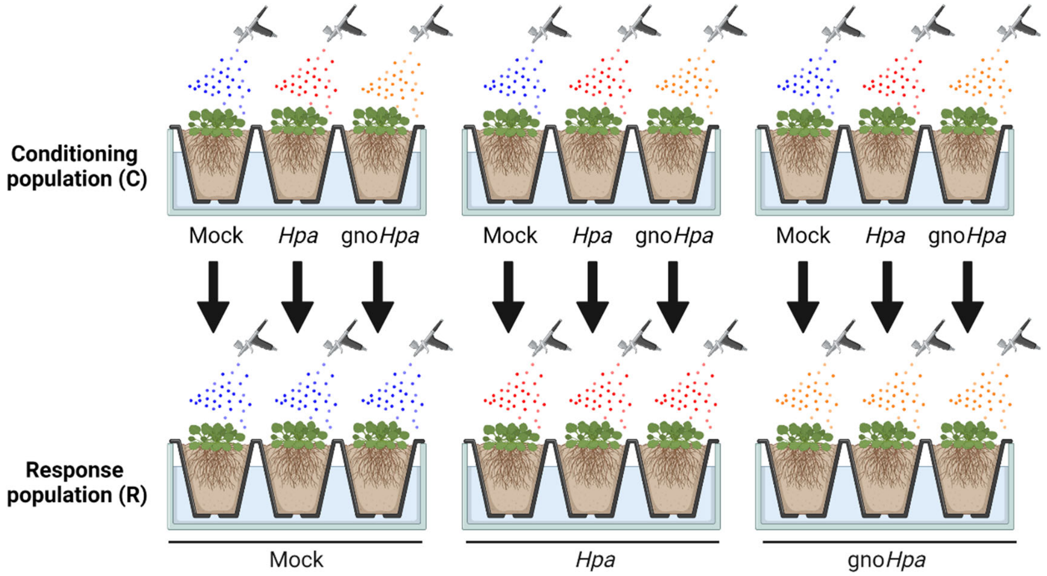


Figure S18. **Setup of soil-borne legacy experiments.** A conditioning population of two-week-old *Arabidopsis thaliana* Col-seedlings was inoculated with mock, *Hpa* Noco2 or gno*Hpa* Noco2. After one week of infection, shoots were cut-off and a response population of plants was directly sown on the conditioned soil and again mock- *Hpa*- or gno*Hpa*-inoculated. Figure created with BioRender.com.

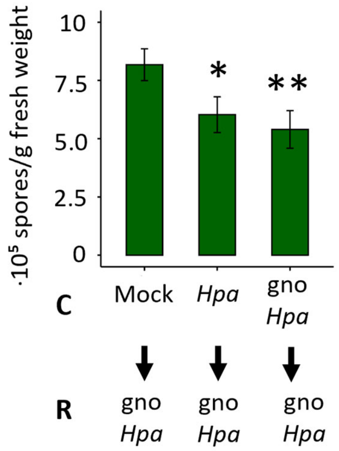


Figure S19. **Gno*Hpa* inoculated Col-0 plants grown on soils conditioned with *Hpa*- or gno*Hpa* inoculated Col-0 plants can perceive soil-borne legacy.** Gno*Hpa s*pore production on a response (R) population of Arabidopsis Col-0 plants growing on soil conditioned by a mock-, *Hpa-,* or gno*Hpa*-inoculated population of Col-0 plants. All bars indicate the average and error bars represent standard error, respectively, from 10 replicate pots. Asterisks indicate significant differences compared to the mock-conditioned population of Col-0 plants in a Student's *t*-test. *: $P < 0.05$, **: $P < 0.01$.

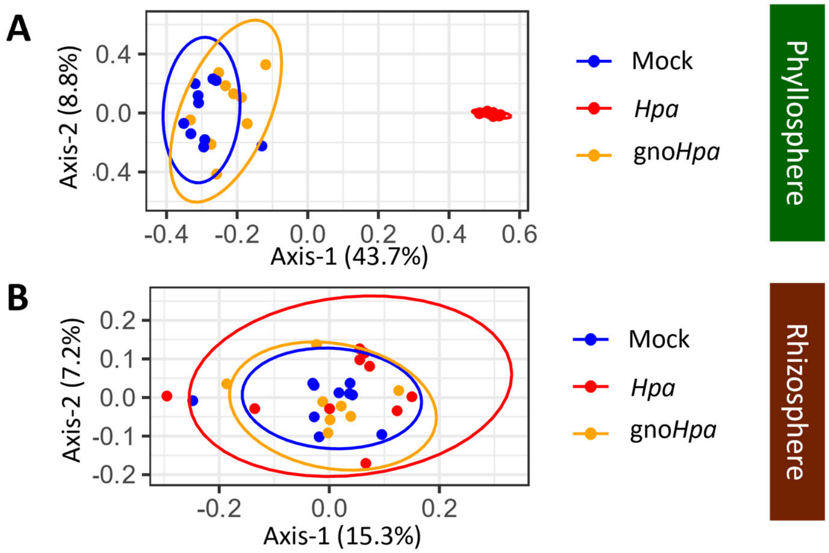


Figure S20. ***Hpa* inoculation of a conditioning population of Col-0 plants drastically alters the phyllosphere microbial community composition, whereas the rhizosphere remains largely unaffected.** PCoA ordination plots based on Bray-Curtis dissimilarities between all (**A**) phyllosphere samples and (**B**) rhizosphere samples from a conditioning population of Col-0 plants that was either mock- (blue), *Hpa*- (red), or gno*Hpa*-inoculated (orange). *Hpa*-inoculation significantly changes (**A**) phyllosphere microbial community composition (PERMANOVA, $R^2$ = 0.58, $P$ < 0.001) whereas (**B**) the rhizosphere remains largely unaffected (PERMANOVA, $R^2$ = 0.05, $P$ = 0.45).

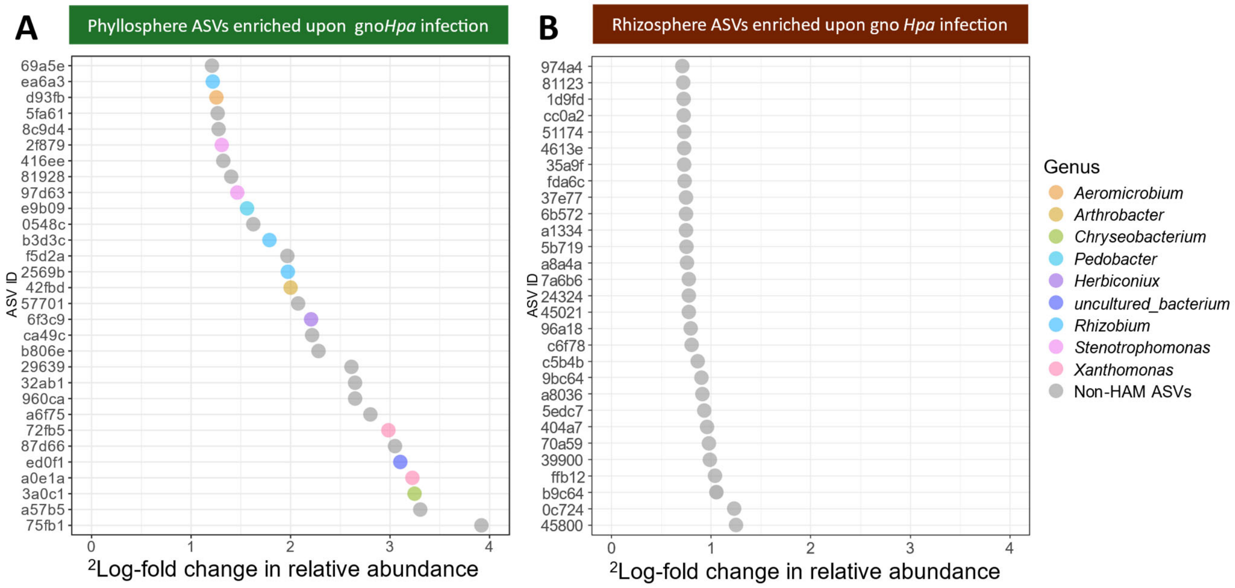


Figure S21. **HAM ASVs are represented among the top-30 most-strongly enriched ASVs in the phyllosphere, but not the rhizosphere of gno*Hpa*-infected plants. (A**) $^{2}$Log-fold change in relative abundance of the top-30 most-strongly enriched ASVs in the phyllosphere of gno*Hpa*-inoculated Col-0 plants grown on gno*Hpa*-conditioned soil compared to mock-inoculated Col-0 plants grown on mock-conditioned soil. Predefined HAM ASVs are colored by genus-level taxonomy whereas non-HAM ASVs are colored grey. Y-axis shows the first 5 letters of the unique identifier for each ASV. (**B**) $^{2}$Log-fold change in relative abundance of the top-30 most-strongly enriched ASVs in the rhizosphere of gno*Hpa*-inoculated Col-0 plants grown on gno*Hpa*-conditioned soil compared to mock-inoculated Col-0 plants grown on mock-conditioned soil. HAM ASVs were not among the top-30 most strongly enriched ASVs in the rhizosphere.

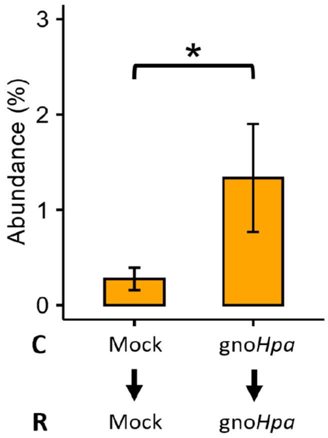


Figure S22. ***Aeromicrobium* ASV d93fb is promoted in the phyllosphere by gno*Hpa* infection.** Relative abundance of *Aeromicrobium* ASV d93fb in the phyllosphere of mock-inoculated Col-0 plants grown on mock-conditioned soils compared to gno*Hpa*-inoculated Col-0 plants grown on gno*Hpa*-conditioned soils. Asterisks indicate significance level based on ANCOM-BC differential abundance testing: *: $P < 0.05$.

Table S1. **Statistical differences (PERMANOVA) between bacterial phyllosphere communities in mock-, Noco2-, and Cala2-treated Arabidopsis plants.** PERMANOVA results of pairwise comparisons (permutations = 9999) between Noco2-inoculated plants, Cala2-inoculated plants, mix-inoculated plants and mock-inoculated plants. *q*-value represents *fdr*-corrected *P*-value. $R^2$ is a measure of effect size.

| **group 1** | **group 2** | $R^2$ | ***q*-value** |
|---|---|---|---|
| Mock | Cala2 | 0.544 | 0.0001 |
| Mock | Noco2 | 0.518 | 0.0001 |
| Mock | Mix | 0.548 | 0.0001 |
| Cala2 | Noco2 | 0.701 | 0.0001 |
| Cala2 | Mix | 0.363 | 0.0001 |
| Noco2 | Mix | 0.477 | 0.0001 |

Table S2. **PERMANOVA of treatment (mock, Noco2, Cala2) and Arabidopsis genotype (C24, Col-0, L*er*, Pro-0) on bacterial phyllosphere communities.** PERMANOVA test of 'Treatment' * 'Accession' with 9999 permutations for Bray-Curtis dissimilarities between all samples from Noco2, Cala2, and mix-inoculated samples in Utrecht. $R^2$ is a measure of effect size.

| | $R^2$ | ***P*-value** |
|---|---|---|
| Treatment | 0.64 | 0.0001 |
| Accession | 0.04 | 0.0066 |
| Treatment * Accession | 0.07 | 0.0559 |

Table S3. **Statistical differences (PERMANOVA) between bacterial phyllosphere communities in mock-, Noco2-, and Cala2-treatment per Arabidopsis genotype.** PERMANOVA results of pairwise comparisons (permutations = 9999) between teatments within Col-0 plants, L*er* plants, Pro-0 plants, and C24 plants. Absence of significant differences observed for C24 plants can be explained by lack of statistical power due to the availability of fewer replicates (indicated by 'x' after the respective treatment). *q*-value represents *fdr*-corrected *P*-value. $R^2$ is a measure of effect size.

| **Col-0 (Total *N* = 16)** | | | |
|---|---|---|---|
| **group 1** | **group 2** | $R^2$ | ***q*-value** |
| Cala2 | Mix | 0.379 | 0.0327 |
| Cala2 | Mock | 0.612 | 0.0281 |
| Cala2 | Noco2 | 0.802 | 0.0297 |
| Mix | Mock | 0.604 | 0.0307 |
| Mix | Noco2 | 0.661 | 0.0297 |
| Mock | Noco2 | 0.605 | 0.0309 |
| **L*er* (Total *N* = 16)** | | | |
| **group 1** | **group 2** | $R^2$ | ***q*-value** |
| Cala2 | Mix | 0.615 | 0.0305 |
| Cala2 | Mock | 0.671 | 0.0283 |
| Cala2 | Noco2 | 0.812 | 0.0279 |
| Mix | Mock | 0.676 | 0.027 |
| Mix | Noco2 | 0.68 | 0.0296 |
| Mock | Noco2 | 0.669 | 0.0289 |
| **C24 (Total *N* = 13)** | | | |
| **group 1** | **group 2** | $R^2$ | ***q*-value** |
| Cala2 | Mix | 0.602 | 0.0286 |
| Cala2 x | Mock x | 0.759 | 0.1 |
| Cala2 x | Noco2 x | 0.872 | 0.1 |
| Mix | Mock | 0.76 | 0.0286 |
| Mix | Noco2 | 0.677 | 0.0286 |
| Mock x | Noco2 x | 0.731 | 0.1 |
| **Pro-0 (Total *N* = 16)** | | | |
| **group 1** | **group 2** | $R^2$ | ***q*-value** |
| Cala2 | Mix | 0.562 | 0.0265 |
| Cala2 | Mock | 0.576 | 0.0277 |
| Cala2 | Noco2 | 0.779 | 0.0256 |
| Mix | Mock | 0.607 | 0.0276 |
| Mix | Noco2 | 0.621 | 0.0279 |
| Mock | Noco2 | 0.576 | 0.0309 |

Table S4. **Number of genomes included in genome index per genus, as used for metagenome analyses (Fig 3A).**

| **Genus** | **#** |
|---|---|
| *Xanthomonas* spp. | 127 |
| *Acidovorax* spp. | 80 |
| *Arthrobacter* spp. | 184 |
| *Sphingobium* spp. | 91 |
| *Rhizobium* spp. | 303 |
| *Aeromicrobium* spp. | 30 |
| *Methylobacterium* spp. | 82 |
| *Microbacterium* spp. | 231 |

Table S5.**Statistical differences (PERMANOVA) in bacterial phyllosphere communities of L*er rpp5* lineages 1-3 (Noco2, Cala2, *Hpa*-free) and untreated L*er rpp5* plants in the 9-passages experiment**. Effect size and significance of *Hpa* removal from its associated HAM microbiome in pairwise PERMANOVA tests of all Treatments with 9999 permutations for Bray-Curtis dissimilarities between all samples from the '9-passages' experiment. The *q*-values represent the *fdr*-corrected *P*-values. $R^2$ is a measure of effect size.

| **Comparison** | $R^2$ | ***q*-value** |
|---|---|---|
| Untreated – *Hpa-free* | 0.328 | 0.0001 |
| Untreated - Noco2 | 0.341 | 0.0001 |
| Untreated - Cala2 | 0.33 | 0.0001 |
| *Hpa-free* - Noco2 | 0.06 | 0.0081 |
| *Hpa-free* - Cala2 | 0.067 | 0.005 |
| Noco2 - Cala2 | 0.017 | 0.6099 |

Table S6. **Statistical differences (PERMANOVA) in bacterial phyllosphere communities of L*er rpp5* lineages 1-3 (Noco2, Cala2, *Hpa*-free) per passage (1, 5, and 9) in the 9-passages experiment**. PERMANOVA results of pairwise comparisons (permutations = 9999) after HAM passages 1, 5, and 9, between Noco2-infected plants, Cala2-infected plants, and plants inoculated with a phyllosphere wash-off from which *Hpa* was removed ('*Hpa*-free'). The *q*-values represent the *fdr*-corrected *P*-values. $R^2$ is a measure of effect size.

| | | **HAM passage 1** | | **HAM passage 5** | | **HAM passage 9** | |
|---|---|---|---|---|---|---|---|
| group 1 | group 2 | $R^2$ | *q*-value | $R^2$ | *q*-value | $R^2$ | *q*-value |
| Noco2 | Cala2 | 0.095 | 0.203 | 0.059 | 0.537 | 0.075 | 0.318 |
| Noco2 | *Hpa*-free | 0.16 | 0,027 | 0.142 | 0.02 | 0.212 | 0.02 |
| Cala2 | *Hpa*-free | 0.131 | 0.107 | 0.132 | 0.02 | 0.237 | 0.018 |

Table S7. **Bacterial ASVs that that are affected by the removal of *Hpa* from the HAM in the 9-passages experiment.** ASVs that were significantly differentially abundant (DESeq2) in the *Hpa*-infected (Noco2 and Cala2) plants compared to *Hpa*-free plants inoculated with passage 9 from the 9-passages experiment.

| **ASV** | **Status in *Hpa* infected plants** | ***fdr*-corrected *P*-value** | **Class** | **Genus** |
|---|---|---|---|---|
| 95a529b86d88a6107aee969a0900df7e | Enriched | 1,71E-03 | *Actinobacteria* | *Nocardioides* |
| 27674392fce5ef7ef32f30e810a0a7c2 | Enriched | 8,17E-04 | *Actinobacteria* | *Nocardioides* |
| f0c7648ba19e0e246a6fbc8fd575ab93 | Enriched | 2,23E-05 | *Actinobacteria* | *Microbacterium* |
| d93fb404358cbc6c5fb5c2c26f76bed6 | Enriched | 1,73E-03 | *Actinobacteria* | *Aeromicrobium* |
| 31c5d6d689fe24ec7157fa1c0783fbd4 | Enriched | 7,57E-17 | *Alphaproteobacteria* | *Methylobacterium* |
| 15da8b3417459a1a66342300e6968701 | Enriched | 4,95E-02 | *Alphaproteobacteria* | *Methylobacterium* |
| 4e248ecf1bc742964ad36d58dc1643e6 | Enriched | 8,17E-04 | *Alphaproteobacteria* | *Rhizobium* |
| 2569b4c4cad75aecc852532bc6bd3c10 | Enriched | 6,05E-04 | *Alphaproteobacteria* | *Rhizobium* |
| e64afb2ca7d2f51d117d7f8cb6fe5e7c | Depleted | 1,17E-19 | *Alphaproteobacteria* | *Rhizobium* |
| 3fa9470e4f41b6f34c5fc5d2a588af89 | Depleted | 1,90E-03 | *Alphaproteobacteria* | *Sphingobium* |
| 9cce057c75c4bf8792364e8776ec3161 | Enriched | 8,17E-04 | *Betaproteobacteria* | *Acidovorax* |
| 7e537c12b9af4d0e06d9c229f25d2176 | Enriched | 5,35E-20 | *Betaproteobacteria* | *Delftia* |
| 18a9493174a87b8ddfa73655faa37a88 | Enriched | 5,45E-20 | *Betaproteobacteria* | *Comamonas* |
| deaf8029eba24ec9cfc9f77e9689423e | Enriched | 3,43E-02 | *Betaproteobacteria* | *Comamonas* |
| fb1944c973cbfd41e00cf593920c788d | Enriched | 1,52E-12 | *Betaproteobacteria* | *Delftia* |
| 819286959c7d21780818a60b9f3e3481 | Depleted | 1,61E-03 | *Betaproteobacteria* | *Methylophilus* |
| 7bae764266c9fa3aa36b95ed1643b550 | Depleted | 4,08E-11 | *Betaproteobacteria* | *Comamonadaceae* |
| 5639089d02ce076046253e2233a57d7c | Depleted | 3,61E-07 | *Cytophagia* | *Arcicella* |
| cc141890f0c2cd9ce45e1ef255a92a05 | Enriched | 1,63E-02 | *Flavobacteriia* | *Flavobacterium* |
| 6ef37d5e731cdd976b90ac058b7c0492 | Depleted | 1,74E-12 | *Flavobacteriia* | *Flavobacterium* |
| 57b1624fb8fa7f753961385792b49ba1 | Enriched | 3,26E-02 | *Gammaproteobacteria* | *Stenotrophomonas* |
| be5f7561425c2fdf15e1347047e3a7ba | Enriched | 1,84E-02 | *Gammaproteobacteria* | *Pseudomonas* |
| a0e1a3759ad3462a28017d94b4432ee1 | Enriched | 8,66E-03 | *Gammaproteobacteria* | *Xanthomonas* |
| 1919e10bb7fe8700ae16e09e536adc7c | Depleted | 1,41E-04 | *Gammaproteobacteria* | *Hydrocarboniphaga* |
| 649c8b8d580e225a6282ad485178b6b8 | Enriched | 2,31E-25 | *Sphingobacteriia* | *Pedobacter* |

Table S8. **Bacterial isolates screened for modulation of *Hpa* reproduction and the effect of *Hpa* on bacterial abundance in a gnotobiotic system**. *Microbacterium* f0c76 #1 and #2 are isolates with identical 16S V3/V4 (ASV) sequences, but with an average nucleotide identity of ~88% based on their genomes.

| **Name** | **Isolate** | **Category** |
|---|---|---|
| *Acidovorax* a4065 | WCS2018Noco2-16 | *Hpa*-associated |
| *Arthrobacter* 42fbd | WCS2018Hpa-7 | *Hpa*-associated |
| *Sphingobium* ed6be | WCS2017Hpa-17 | *Hpa*-associated |
| *Xanthomonas* a0e1a | WCS2014-23R | *Hpa*-associated |
| *Aeromicrobium* d93fb | WCS2018Hpa-31 | *Hpa*-associated |
| *Rhizobium* 2569b | WCS2018Hpa-8 | *Hpa*-associated |
| *Microbacterium* f0c76 #1 | WCS2018Hpa-23 | *Hpa*-associated |
| *Microbacterium* f0c76 #2 | WCS2018Hpa-9 | *Hpa*-associated |
| *Methylobacterium* 15da8 | WCS2018Hpa-22 | *Hpa*-associated |
| *Pseudomonas* 7d105 | CN2-GNA-2 | Phyllosphere resident |
| *Asticcacaulis* 70cff | PN-R2A-26 | Phyllosphere resident |
| *Duganella* f90ae | PN-TSA-11 | Phyllosphere resident |
| *Pseudomonas* fb830 | PN-YEM-6 | Phyllosphere resident |

Table S9. **ASVs that were significantly enriched in *Hpa*-treated phyllosphere samples in the conditioning phase of the soil-borne legacy experiment (Fig. 6C).** Differential abundances testing was performed with DESeq2 (*fdr*-corrected Wald tests, indicated by '*P*-adj') for *Hpa*-inoculated samples compared to mock-treated samples.

| **Family** | **Genus** | **ASV** | **log2FoldChange** | ***P*-adj** | **Mean abundance (%) in *Hpa*-inoculated plants** |
|---|---|---|---|---|---|
| *Methylophilaceae* | *Methylophilus* | e50db | 8,32 | 1.00E-21 | 13,31 |
| *Nocardioidaceae* | *Aeromicrobium* | d93fb | 7,64 | 8,1E-16 | 13,06 |
| *Flavobacteriaceae* | *Chryseobacterium* | 3a0c1 | 7,81 | 1,7E-23 | 10,59 |
| *Xanthomonadaceae* | *Xanthomonas* | a0e1a | 6,37 | 4,8E-18 | 10,09 |
| *Flavobacteriaceae* | *Flavobacterium* | ef66d | 8,05 | 1,3E-17 | 9,59 |
| *Micrococcaceae* | *Arthrobacter* | 42fbd | 4,96 | 1,1E-16 | 4,74 |
| *Cytophagaceae* | *Dyadobacter* | eabab | 7,86 | 4,8E-18 | 3,66 |
| *Caulobacteraceae* | *Brevundimonas* | eb229 | 7,67 | 2,2E-12 | 3,50 |
| *Sphingomonadaceae* | *Sphingobium* | ed6be | 5,32 | 8,1E-12 | 2,94 |
| *Hyphomicrobiaceae* | *Devosia* | 29b65 | 5,11 | 3,8E-06 | 2,38 |
| *Comamonadaceae* | *Acidovorax* | 9cce0 | 5,53 | 0,00013 | 1,73 |
| *Rhizobiaceae* | *Rhizobium* | 2569b | 3,86 | 0,00208 | 1,53 |
| *Sphingobacteriaceae* | *Pedobacter* | f2a1b | 4,24 | 5,4E-06 | 1,27 |
| *Sphingobacteriaceae* | *Pedobacter* | e9b09 | 7,68 | 2,2E-11 | 0,99 |
| *Comamonadaceae* | *Acidovorax* | 9f6d2 | 5,91 | 3.00E-05 | 0,88 |
| *Xanthomonadaceae* | *Stenotrophomonas* | 77da9 | 6,07 | 5,8E-06 | 0,72 |
| *Comamonadaceae* | *Acidovorax* | a4065 | 3,9 | 1,8E-05 | 0,70 |
| *Microbacteriaceae* | *Agromyces* | efbd0 | 5,89 | 1,6E-07 | 0,67 |
| *Sphingobacteriaceae* | *Sphingobacterium* | 5d7e3 | 6,53 | 2,3E-06 | 0,59 |
| *uncultured_bacterium* | *uncultured_bacterium* | ed0f1 | 7,4 | 4,6E-11 | 0,57 |
| *Sphingomonadaceae* | *Sphingopyxis* | 41936 | 4,34 | 0,00025 | 0,50 |
| *Sphingobacteriaceae* | *Pedobacter* | 0795a | 16,45 | 1,6E-35 | 0,47 |
| *Sphingobacteriaceae* | *Pedobacter* | 57798 | 7,52 | 8,1E-08 | 0,43 |
| *Rhizobiaceae* | *Rhizobium* | f96ad | 1,96 | 0,00967 | 0,40 |
| *Comamonadaceae* | *Variovorax* | 3c699 | 6,29 | 4,3E-05 | 0,32 |
| *Comamonadaceae* | *Comamonas* | 18a94 | 6,68 | 1.00E-06 | 0,31 |
| *Comamonadaceae* | *Variovorax* | f6bf5 | 5,85 | 0,00019 | 0,29 |
| *Rhizobiaceae* | *Rhizobium* | aca76 | 7,67 | 5.00E-08 | 0,26 |
| *Sphingobacteriaceae* | *Sphingobacterium* | d0c4a | 5,78 | 3,4E-05 | 0,21 |
| *Methylobacteriaceae* | *Methylobacterium* | 15da8 | 0,54 | 4,6E-05 | 0,21 |
| *Xanthomonadaceae* | *Stenotrophomonas* | 57b16 | 1,01 | 0,00361 | 0,18 |
| *Xanthomonadaceae* | *Xanthomonas* | 72fb5 | 9,03 | 5.00E-17 | 0,18 |
| *Sphingobacteriaceae* | *Sphingobacterium* | d5416 | 9,06 | 2,9E-14 | 0,15 |
| *Nocardioidaceae* | *Aeromicrobium* | 7451e | 14,59 | 1,2E-24 | 0,15 |
| *Microbacteriaceae* | *Herbiconiux* | 6f3c9 | 0,59 | 0,00077 | 0,12 |
| *Sphingobacteriaceae* | *Sphingobacterium* | aec8d | 14,1 | 1,2E-08 | 0,12 |
| *Brucellaceae* | *Ochrobactrum* | 609fb | 0,07 | 1,2E-05 | 0,12 |
| *Comamonadaceae* | *NA* | 1da8b | 8,46 | 6,4E-16 | 0,10 |
| *uncultured_bacterium* | *uncultured_bacterium* | 13100 | 12,93 | 1,8E-07 | 0,06 |
| *Nocardioidaceae* | *Aeromicrobium* | 484fb | 12,48 | 2.00E-15 | 0,04 |
| *Xanthomonadaceae* | *Stenotrophomonas* | 97d63 | 0,93 | 0,0147 | 0,04 |

| | | | | | |
|---|---|---|---|---|---|
| *Nocardioidaceae* | *Nocardioides* | 27674 | 12,2 | 2,9E-14 | 0,04 |
| *Sphingomonadaceae* | *Sphingomonas* | 2f935 | 11,56 | 6,8E-05 | 0,03 |
| *Caulobacteraceae* | *Brevundimonas* | 973cd | 3,55 | 0,018 | 0,03 |
| *Streptomycetaceae* | *Streptomyces* | 658ee | 5,29 | 0,0002 | 0,03 |
| *Comamonadaceae* | *NA* | 8accf | 5,44 | 2.00E-05 | 0,02 |
| *Comamonadaceae* | *NA* | 5538f | 5,16 | 3,8E-06 | 0,02 |
| *Sphingobacteriaceae* | *Sphingobacterium* | f56f2 | 10,68 | 0,00011 | 0,02 |
| *Sphingobacteriaceae* | *Pedobacter* | d96e0 | 2,12 | 0,047 | 0,02 |
| *Coxiellaceae* | *uncultured* | f3109 | 4,46 | 0,00338 | 0,02 |
| *Xanthomonadaceae* | *Xanthomonas* | 983fc | 7,99 | 0,00011 | 0,02 |
| *Moraxellaceae* | *Alkanindiges* | b8f0f | 7,14 | 0,00013 | 0,02 |

Table S10. **Primers used in this study.**

| Name | Sequence (5' - 3') | Used for | Reference |
|---|---|---|---|
| AtActFwd | AATCACAGCACTTGCACCA | qPCR quantification of *Hpa* abundance | Anderson *et al.* (2015)[55] |
| ActActRv | GAGGGAAGCAAGAATGGAAC | qPCR quantification of *Hpa* abundance | Anderson *et al.* (2015)[55] |
| HpaActFwd | GTGTCGCACACTGTACCCATTTAT | qPCR quantification of *Hpa* abundance | Anderson *et al.* (2015)[55] |
| HpaActFwd | ATCTTCATCATGTAGTCGGTCAAGT | qPCR quantification of *Hpa* abundance | Anderson *et al.* (2015)[55] |
| 16S-Fw-ill | TCGTCGGCAGCGTCAGATGTGTATAAGAGACAG CCTACGGGNGGCWGCAG | PCR1 - Illumina 16S rDNA library preparation | Illumina |
| 16S-Rv-ill | GTCTCGTGGGCTCGGAGATGTGTATAAGAGACA GGACTACHVGGGTATCTAATCC | PCR1 - Illumina 16S rDNA library preparation | Illumina |
| NGS1-16s-N701 | TCGTCGGCAGCGTCAGATGTGTATAAGAGACAG TCGCCTTACCTGTGGCCTACGGGNGGCWGCAG | PCR1 - Illumina 16S rDNA library preparation | This study - with heterogeneity spacers based on de Muinck *et al*. (2017)[56] |
| NGS1-16s-N702 | TCGTCGGCAGCGTCAGATGTGTATAAGAGACAG CTAGTACGGAGTGGCCTACGGGNGGCWGCAG | PCR1 - Illumina 16S rDNA library preparation | This study - with heterogeneity spacers based on de Muinck *et al.* (2017)[56] |
| NGS1-16s-N703 | TCGTCGGCAGCGTCAGATGTGTATAAGAGACAG TTCTGCCTTGCGACCTACGGGNGGCWGCAG | PCR1 - Illumina 16S rDNA library preparation | This study - with heterogeneity spacers based on de Muinck *et al.* (2017)[56] |
| NGS1-16s-N704 | TCGTCGGCAGCGTCAGATGTGTATAAGAGACAG GCTCAGGAATGACCTACGGGNGGCWGCAG | PCR1 - Illumina 16S rDNA library preparation | This study - with heterogeneity spacers based on de Muinck *et al.* (2017)[56] |
| NGS1-16s-N705 | TCGTCGGCAGCGTCAGATGTGTATAAGAGACAG AGGAGTCCCGACCTACGGGNGGCWGCAG | PCR1 - Illumina 16S rDNA library preparation | This study - with heterogeneity spacers based on de Muinck *et al.* (2017)[56] |
| NGS1-16s-N706 | TCGTCGGCAGCGTCAGATGTGTATAAGAGACAG CATGCCTACGACCTACGGGNGGCWGCAG | PCR1 - Illumina 16S rDNA library preparation | This study - with heterogeneity spacers based on de Muinck *et al.* (2017)[56] |
| NGS1-16s-N707 | TCGTCGGCAGCGTCAGATGTGTATAAGAGACAG GTAGAGAGGTCCTACGGGNGGCWGCAG | PCR1 - Illumina 16S rDNA library preparation | This study - with heterogeneity spacers based on de Muinck *et al.* (2017)[56] |
| NGS1-16s-N708 | TCGTCGGCAGCGTCAGATGTGTATAAGAGACAG CCTCTCTGGTCCTACGGGNGGCWGCAG | PCR1 - Illumina 16S rDNA library preparation | This study - with heterogeneity spacers based on de Muinck *et al.* (2017)[56] |
| NGS1-16s-N709 | TCGTCGGCAGCGTCAGATGTGTATAAGAGACAG AGCGTAGCTCCTACGGGNGGCWGCAG | PCR1 - Illumina 16S rDNA library preparation | This study - with heterogeneity spacers based on de Muinck *et al.* (2017)[56] |
| NGS1-16s-N710 | TCGTCGGCAGCGTCAGATGTGTATAAGAGACAG CAGCCTCGTCCTACGGGNGGCWGCAG | PCR1 - Illumina 16S rDNA library preparation | This study - with heterogeneity spacers based on de Muinck *et al.* (2017)[56] |
| NGS1-16s-N711 | TCGTCGGCAGCGTCAGATGTGTATAAGAGACAG TGCCTCTTCCTACGGGNGGCWGCAG | PCR1 - Illumina 16S rDNA library preparation | This study - with heterogeneity spacers based on de Muinck *et al.* (2017)[56] |
| NGS1-16s-N712 | TCGTCGGCAGCGTCAGATGTGTATAAGAGACAG TCCTCTACCCTACGGGNGGCWGCAG | PCR1 - Illumina 16S rDNA library preparation | This study - with heterogeneity spacers based on de Muinck *et al.* (2017)[56] |
| NGS1-16s-N501 | GTCTCGTGGGCTCGGAGATGTGTATAAGAGACA GTAGATCGCCACTTCTGACTACHVGGGTATCTAA TCC | PCR1 - Illumina 16S rDNA library preparation | This study - with heterogeneity spacers based on de Muinck *et al.* (2017)[56] |
| NGS1-16s-N502 | GTCTCGTGGGCTCGGAGATGTGTATAAGAGACA GCTCTCTATTTCTCTGACTACHVGGGTATCTAATC C | PCR1 - Illumina 16S rDNA library preparation | This study - with heterogeneity spacers based on de Muinck *et al.* (2017)[56] |

| | | | |
|---|---|---|---|
| NGS1-16s-N503 | GTCTCGTGGGCTCGGAGATGTGTATAAGAGACAGTATCCTCTACTCAGACTACHVGGGTATCTAATCC | PCR1 - Illumina 16S rDNA library preparation | This study - with heterogeneity spacers based on de Muinck *et al.* (2017)[56] |
| NGS1-16s-N504 | GTCTCGTGGGCTCGGAGATGTGTATAAGAGACAGAGAGTAGAGATAGACTACHVGGGTATCTAATCC | PCR1 - Illumina 16S rDNA library preparation | This study - with heterogeneity spacers based on de Muinck *et al.* (2017)[56] |
| NGS1-16s-N505 | GTCTCGTGGGCTCGGAGATGTGTATAAGAGACAGGTAAGGAGCTAGACTACHVGGGTATCTAATCC | PCR1 - Illumina 16S rDNA library preparation | This study - with heterogeneity spacers based on de Muinck *et al.* (2017)[56] |
| NGS1-16s-N506 | GTCTCGTGGGCTCGGAGATGTGTATAAGAGACAGACTGCATATCGACTACHVGGGTATCTAATCC | PCR1 - Illumina 16S rDNA library preparation | This study - with heterogeneity spacers based on de Muinck *et al.* (2017)[56] |
| NGS1-16s-N507 | GTCTCGTGGGCTCGGAGATGTGTATAAGAGACAGAAGGAGTAAGACTACHVGGGTATCTAATCC | PCR1 - Illumina 16S rDNA library preparation | This study - with heterogeneity spacers based on de Muinck *et al.* (2017)[56] |
| NGS1-16s-N508 | GTCTCGTGGGCTCGGAGATGTGTATAAGAGACAGCTAAGCCTGACTACHVGGGTATCTAATCC | PCR1 - Illumina 16S rDNA library preparation | This study - with heterogeneity spacers based on de Muinck *et al.* (2017)[56] |
| S501 | AATGATACGGCGACCACCGAGATCTACACTAGATCGCTCGTCGGCAGCGTC | PCR2 - Illumina 16S rDNA library preparation | Illumina |
| S502 | AATGATACGGCGACCACCGAGATCTACACCTCTCTATTCGTCGGCAGCGTC | PCR2 - Illumina 16S rDNA library preparation | Illumina |
| S503 | AATGATACGGCGACCACCGAGATCTACACTATCCTCTTCGTCGGCAGCGTC | PCR2 - Illumina 16S rDNA library preparation | Illumina |
| S504 | AATGATACGGCGACCACCGAGATCTACACAGAGTAGATCGTCGGCAGCGTC | PCR2 - Illumina 16S rDNA library preparation | Illumina |
| S505 | AATGATACGGCGACCACCGAGATCTACACGTAAGGAGTCGTCGGCAGCGTC | PCR2 - Illumina 16S rDNA library preparation | Illumina |
| S506 | AATGATACGGCGACCACCGAGATCTACACACTGCATATCGTCGGCAGCGTC | PCR2 - Illumina 16S rDNA library preparation | Illumina |
| S507 | AATGATACGGCGACCACCGAGATCTACACAAGGAGTATCGTCGGCAGCGTC | PCR2 - Illumina 16S rDNA library preparation | Illumina |
| S508 | AATGATACGGCGACCACCGAGATCTACACCTAAGCCTTCGTCGGCAGCGTC | PCR2 - Illumina 16S rDNA library preparation | Illumina |
| N701 | CAAGCAGAAGACGGCATACGAGATTCGCCTTAGTCTCGTGGGCTCGG | PCR2 - Illumina 16S rDNA library preparation | Illumina |
| N702 | CAAGCAGAAGACGGCATACGAGATCTAGTACGGTCTCGTGGGCTCGG | PCR2 - Illumina 16S rDNA library preparation | Illumina |
| N703 | CAAGCAGAAGACGGCATACGAGATTTCTGCCTGTCTCGTGGGCTCGG | PCR2 - Illumina 16S rDNA library preparation | Illumina |
| N704 | CAAGCAGAAGACGGCATACGAGATGCTCAGGAGTCTCGTGGGCTCGG | PCR2 - Illumina 16S rDNA library preparation | Illumina |
| N705 | CAAGCAGAAGACGGCATACGAGATAGGAGTCCGTCTCGTGGGCTCGG | PCR2 - Illumina 16S rDNA library preparation | Illumina |
| N706 | CAAGCAGAAGACGGCATACGAGATCATGCCTAGTCTCGTGGGCTCGG | PCR2 - Illumina 16S rDNA library preparation | Illumina |

| | | | |
|---|---|---|---|
| N707 | CAAGCAGAAGACGGCATACGAGATGTAGAGAG GTCTCGTGGGCTCGG | PCR2 - Illumina 16S rDNA library preparation | Illumina |
| N708 | CAAGCAGAAGACGGCATACGAGATCCTCTCTGG TCTCGTGGGCTCGG | PCR2 - Illumina 16S rDNA library preparation | Illumina |
| N709 | CAAGCAGAAGACGGCATACGAGATAGCGTAGC GTCTCGTGGGCTCGG | PCR2 - Illumina 16S rDNA library preparation | Illumina |
| N710 | CAAGCAGAAGACGGCATACGAGATCAGCCTCGG TCTCGTGGGCTCGG | PCR2 - Illumina 16S rDNA library preparation | Illumina |
| N711 | CAAGCAGAAGACGGCATACGAGATTGCCTCTTG TCTCGTGGGCTCGG | PCR2 - Illumina 16S rDNA library preparation | Illumina |
| N712 | CAAGCAGAAGACGGCATACGAGATTCCTCTACG TCTCGTGGGCTCGG | PCR2 - Illumina 16S rDNA library preparation | Illumina |
| pPNA | GGCTCAACCCTGGACAG | PCR1 PCR clamp | Lundberg *et al*. (2013)[57] |
| mPNA | GGCAAGTGTTCTTCGGA | PCR1 PCR clamp | Lundberg *et al*. (2013)[57] |
| fITS7 | TCGTCGGCAGCGTCAGATGTGTATAAGAGACAG GTGARTCATCGAATCTTTG | PCR1 ITS2 fw primer | Ihrmark *et al*. (2012)[58] |
| ITS4 | GTCTCGTGGGCTCGGAGATGTGTATAAGAGACA GTCCTCCGCTTATTGATATGC | PCR2 ITS2 fw primer | Ihrmark *et al*. (2012)[58] |
| cl1ITS2-F | CGTCTGCCTGGGTGTCACAAATCGTCGTCC | ITS2 blocking oligonucleotide | Agler *et al*. (2016a)[59] |
| clITS2-R | CCTGGTGTCGCTATATGGACTTTGGGTCAT | ITS2 blocking oligonucleotide | Agler *et al*. (2016a)[59] |
